# Exposure of *Mycobacterium tuberculosis* to human alveolar lining fluid shows temporal and strain-specific adaptation to the lung environment

**DOI:** 10.1101/2023.09.27.559381

**Authors:** Anna Allué-Guardia, Andreu Garcia-Vilanova, Alyssa M. Schami, Angélica M. Olmo-Fontánez, Amberlee Hicks, Jay Peters, Diego J. Maselli, Mark D. Wewers, Yufeng Wang, Jordi B. Torrelles

**Author notes:** **Corresponding authors**: Drs. Anna Allué-Guardia, PhD and Jordi B. Torrelles, PhD.

## Abstract

Upon infection, *Mycobacterium tuberculosis* (*M.tb*) reaches the alveolar space and comes in close contact with human alveolar lining fluid (ALF) for an uncertain period of time prior to its encounter with alveolar cells. We showed that homeostatic ALF hydrolytic enzymes modify the *M.tb* cell envelope, driving *M.tb*-host cell interactions. Still, the contribution of ALF during *M.tb* infection is poorly understood. Here, we exposed 4 *M.tb* strains with different levels of virulence, transmissibility, and drug resistance (DR) to physiological concentrations of human ALF for 15-min and 12-h, and performed RNA sequencing. Gene expression analysis showed a temporal and strain-specific adaptation to human ALF. Differential expression (DE) of ALF-exposed *vs.* unexposed *M.tb* revealed a total of 397 DE genes associated with lipid metabolism, cell envelope and processes, intermediary metabolism and respiration, and regulatory proteins, among others. Most DE genes were detected at 12-h post-ALF exposure, with DR-*M.tb* strain W-7642 having the highest number of DE genes. Interestingly, genes from the KstR2 regulon, which controls the degradation of cholesterol C and D rings, were significantly upregulated in all strains post-ALF exposure. These results indicate that *M.tb*-ALF contact drives initial metabolic and physiologic changes in *M.tb*, with potential implications in infection outcome.

**IMPORTANCE:** Tuberculosis, caused by airborne pathogen *Mycobacterium tuberculosis* (*M.tb*), is one of the leading causes of mortality worldwide. Upon infection, *M.tb* reaches the alveoli and gets in contact with human alveolar lining fluid (ALF), where ALF hydrolases modify the *M.tb* cell envelope driving subsequent *M.tb*-host cell interactions. Still, the contributions of ALF during infection are poorly understood. We exposed 4 *M.tb* strains to ALF for 15-min and 12-h and performed RNA sequencing, demonstrating a temporal and strain-specific adaptation of *M.tb* to ALF. Interestingly, genes associated with cholesterol degradation were highly upregulated in all strains. This study shows for the first time that ALF drives global metabolic changes in *M.tb* during the initial stages of the infection, with potential implications in disease outcome. Biologically relevant networks and common and strain-specific bacterial determinants derived from this study could be further investigated as potential therapeutic candidates.

## INTRODUCTION

Airborne pathogen *Mycobacterium tuberculosis* (*M.tb*) is the causative agent of tuberculosis (TB), one of the top leading causes of death worldwide due to a single infectious agent, with 11 million cases and ∼1.6 million attributed deaths in 2021 and many underreported diagnoses due to the coronavirus disease 2019 (COVID-19) pandemic disruptions in global healthcare (WHO 2022). Indeed, a 20% raise in the number of TB deaths is predicted over the next 5 years, resulting in a setback of at least 5 to 8 years in End-TB Strategy (Cilloni et al. 2020; Hogan et al. 2020). Further, relocation of resources and access limitations to healthcare during the COVID-19 pandemic predict an increase in the number of drug-resistant (DR-) TB cases worldwide (Wingfield et al. 2021).

During the process of natural infection, *M.tb* is initially deposited in the lung alveolar space where it first encounters the lung mucosa, composed of a surfactant lipid layer and an aqueous hypophase (called alveolar lining fluid or ALF). *M.tb* is thought to be in close contact with ALF for an undefined period (from min to h) before encountering host alveolar resident cells and after its escape from dying cells during the initial phases of infection, where it can interact with ALF innate soluble components (Torrelles and Schlesinger 2017). Indeed, homeostatic hydrolytic activities (hydrolases) present in human ALF, whose main function is ALF degradation and recycling, alter the *M.tb* cell envelope upon ALF-*M.tb* exposure in as little as 15 min by significantly reducing the total amount of two major *M.tb* cell surface virulence factors, the mannose-capped lipoarabinomannan (ManLAM) and trehalose dimycolate (TDM), among others (Arcos et al. 2011). These ALF-induced cell envelope alterations in *M.tb* are shown to influence subsequent interactions with immune host cells *in vitro* and *in vivo,* ultimately determining the infection outcome (Arcos et al. 2011; Arcos et al. 2015; Arcos et al. 2017; Scordo et al. 2017; Moliva et al. 2019; Scordo et al. 2019; Olmo-Fontánez et al. 2021). In addition to ALF hydrolases, levels and functionality of other ALF soluble components are also altered significantly in some human populations such as the elderly and people living with HIV (Moliva et al. 2014; Moliva et al. 2019; Olmo-Fontánez et al. 2021). Thus, the host ALF status and functionality might play an important role in shaping the *M.tb* cell envelope during the initial stages of the infection (Moliva et al. 2014; Moliva et al. 2019). However, the overall impact of the ALF on the adaptation of *M.tb* to the alveolar space prior to being recognized by host cells and subsequent infection is still largely unknown.

Biochemical studies have also determined that the *M.tb* cell envelope composition is quite diverse between phylogenetic lineages and/or geographical regions, and even among strains with different drug resistance patterns (Reed et al. 2007; Huet et al. 2009; Velayati et al. 2009; Pal et al. 2017; Howard et al. 2018; Birhanu et al. 2019), which can lead to different interactions with host immune cells (Allue-Guardia et al. 2021). Still, it is uncertain if different *M.tb* strains/lineages exposed to human ALF will have distinct metabolic responses prior to encountering host cells.

In this study, we aimed to determine the overall and strain-specific effect of ALF exposure on different *M.tb* genomic backgrounds driving subsequent *M.tb*-host interactions (Arcos et al. 2011; Arcos et al. 2015; Arcos et al. 2017; Scordo et al. 2017; Moliva et al. 2019; Scordo et al. 2019; Olmo-Fontánez et al. 2021). To define *M.tb* metabolic adaptations to the human ALF environment, we performed RNA sequencing of four different *M.tb* strains (with different levels of virulence, transmissibility, and drug resistance) exposed to human ALF, and determined shared and strain-specific DE genes between strains when compared to unexposed bacteria (**Supplemental Figure S1**). Our results indicate that time-dependent exposure of *M.tb* to ALF drives global metabolic and physiologic changes in *M.tb* with potential implications in the establishment and outcome of the infection.

## RESULTS

### Alignment statistics, PCA and clustering of samples

After 4 different *M.tb* strains (highly transmissible CDC1551, virulent H_37_R_v_, hypervirulent HN878, and drug-resistant W-7642) were exposed for different lengths of time to human ALF at its physiological concentrations within the lung, bacterial RNA was extracted and sequenced. The total average number of reads/sample after removing low quality reads during the initial quality control (QC) was 6.15 million, ranging from ∼ 4.1 to 11.3 millions. The number of aligned reads to the *M.tb* reference genome was ∼ 90% on average. Samples with less than 70% of aligned reads were analyzed using Kraken (Davis et al. 2013) to discard potential sample contamination (data not shown). All samples had a coverage of more than 97% of their genome after alignment, with an average coverage depth of 193x and GC content of 62%. A Pearson correlation matrix was computed, showing a correlation of more than r = 0.97 between sample replicates (**Supplemental Figure S2**). Our quality post-alignment statistics and other quality control metrics are found in **Supplemental Table S1**.

Raw aligned data was then transformed and used for sample clustering and Principal Component Analysis (PCA). PCA grouped the samples based on *M.tb* strain (CDC1551, H_37_R_v_, HN878, W-7642) and ALF exposure time (15-min and 12-h), indicating temporal and strain-specific changes in transcriptional profiles of *M.tb* after ALF exposure (**Figure 1A**). Strain CDC1551 was furthest from the others at both exposure time points tested, suggesting a more distinct adaptation to the ALF lung environment. Hierarchical clustering of normalized gene expression (shown as a heatmap) confirmed the PCA groups, and classified the samples in four main clusters (**Figure 1B**). Cluster 1 (HN878 and W-7642 at 15-min) was closest to cluster 2 (HN878 and W-7642 at 12-h), indicating that those two strains presented the most similar transcriptional profile at each of the time points tested. They were followed by H_37_R_v_ at both time points (cluster 3) and CDC1551 (cluster 4), which was again separate from the others.

**Figure 1.**
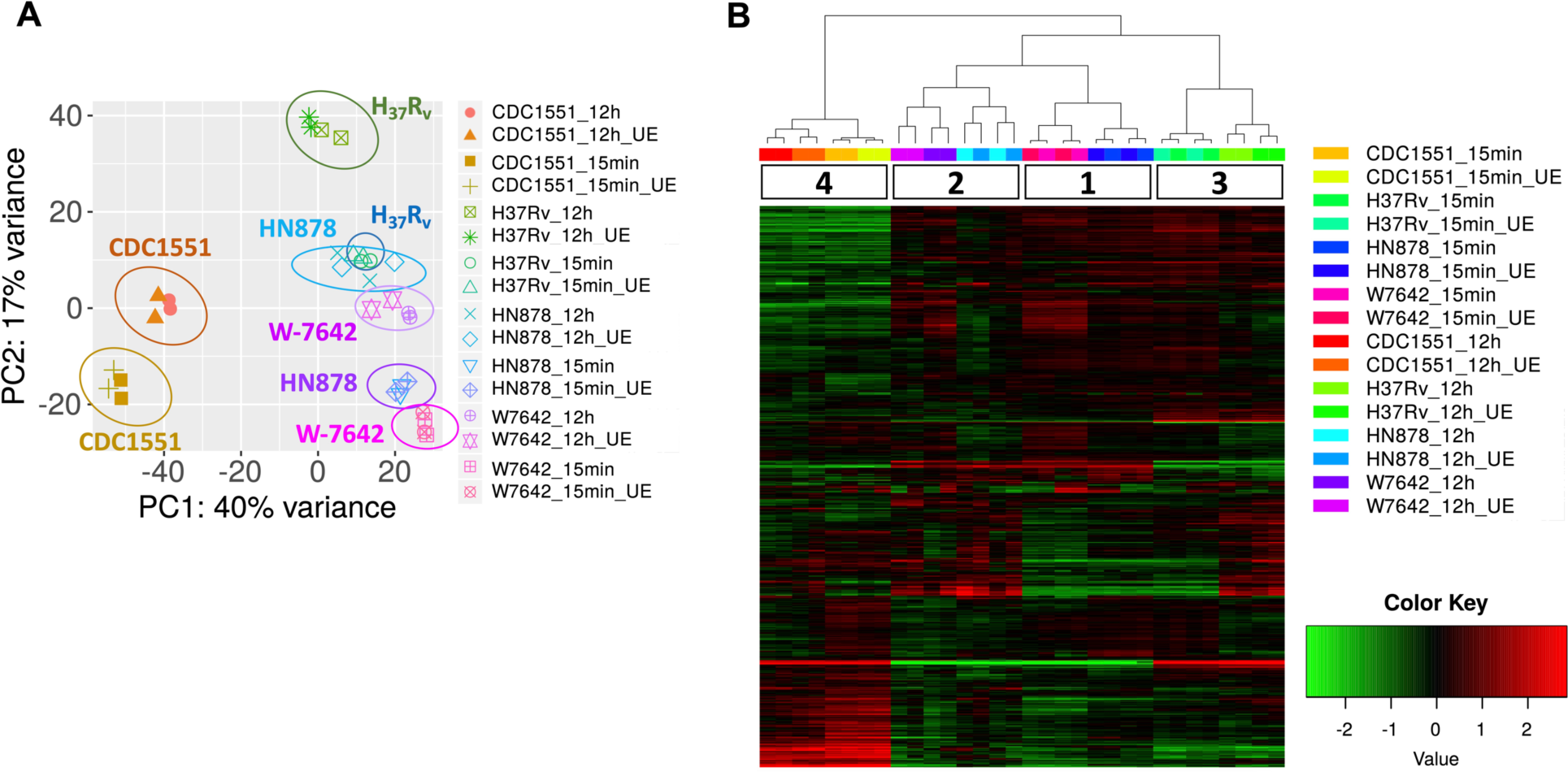
**A**) Principal Component Analysis and **B**) heatmap of normalized gene expression (red: upregulated, green: downregulated) with hierarchical clustering of samples analyzed in this study: four *M.tb* strains (CDC1551, H_37_R_v_, HN878, W-7642), each exposed to a human ALF pool at its physiological concentration or unexposed (UE) for 15-min or 12-h. Replicates (n=2) are shown in the same symbol and/or color. Graphs were generated in iDEP.

### Functional categories and enrichment of Differentially Expressed Genes (DEGs)

To assess the contribution of ALF in the adaptation of *M.tb* to the lung environment in the early stages of the infection, we performed DE analysis of ALF-exposed *vs.* unexposed *M.tb* for each of the strains at both time points. A total of 397 DEGs were identified, with DR-*M.tb* strain W-7642 being the one with the most transcriptional changes after exposure to ALF for 12-h (152 downregulated and 87 upregulated genes) compared to unexposed bacteria (**Figure 2A**). Exposure for 15-min to ALF resulted in a few DEGs: 32 in CDC1551, 12 in H_37_R_v_, 9 in HN878, and 6 in W-7642. Hierarchical clustering using DEGs clearly classified our samples in two main clusters based on the ALF exposure time (15-min *vs.* 12-h) (**Figure 2B**), suggesting a temporal adaptation to this early lung microenvironment.

**Figure 2.**
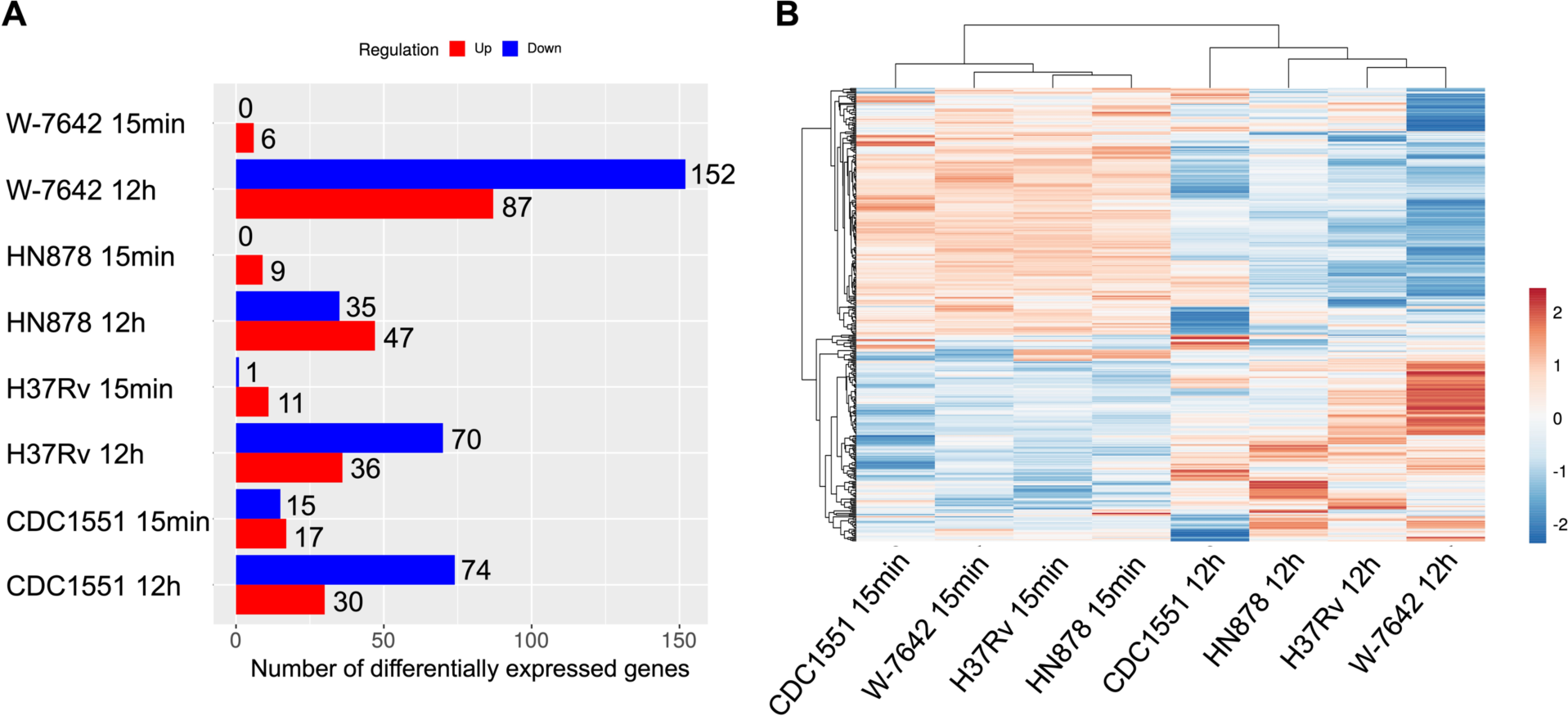
Differential expression analysis (ALF-exposed *M.tb vs.* unexposed *M.tb*) was performed for each of the strains and time points (15-min and 12-h) using the DeSeq2 method in the iDEP software, with the settings: log_2_FC greater than an absolute value of 1, FDR >0.1. **A**) Number of differentially expressed genes (upregulated in red, downregulated in blue) for each of the strains. **B**) A heatmap of DEGs was constructed using log_2_FC values in ClustVis (https://biit.cs.ut.ee/clustvis/) (Metsalu and Vilo 2015), showing separation of strains based on ALF exposure time. Upregulated genes: orange; downregulated genes: blue.

Identified DEGs belonged to different functional categories, including but not restricted to (from most to least number of DEGs): conserved hypotheticals (#109), intermediary metabolism and respiration (#82), cell wall and cell processes (#65), virulence, detoxification and adaptation (#36), lipid metabolism (#35), PE/PPE (#22), information pathways (#18), and regulatory proteins (#17) (**Figure 3A, Supplemental Table S2**). At 15-min post-ALF exposure, the majority of the DEGs were associated to either lipid or intermediate metabolism, respiration, regulatory proteins or cell wall processes, except for strain CDC1551 that included other categories such as information pathways and PE/PPE (**Figure 3B**). However, most transcriptional changes occurred at 12-h after ALF exposure, where we found DEGs corresponding to additional categories like virulence, detoxification and host adaptation, as well as insertion sequences and mobile genetic elements (MGEs) (**Figure 3B**).

**Figure 3.**
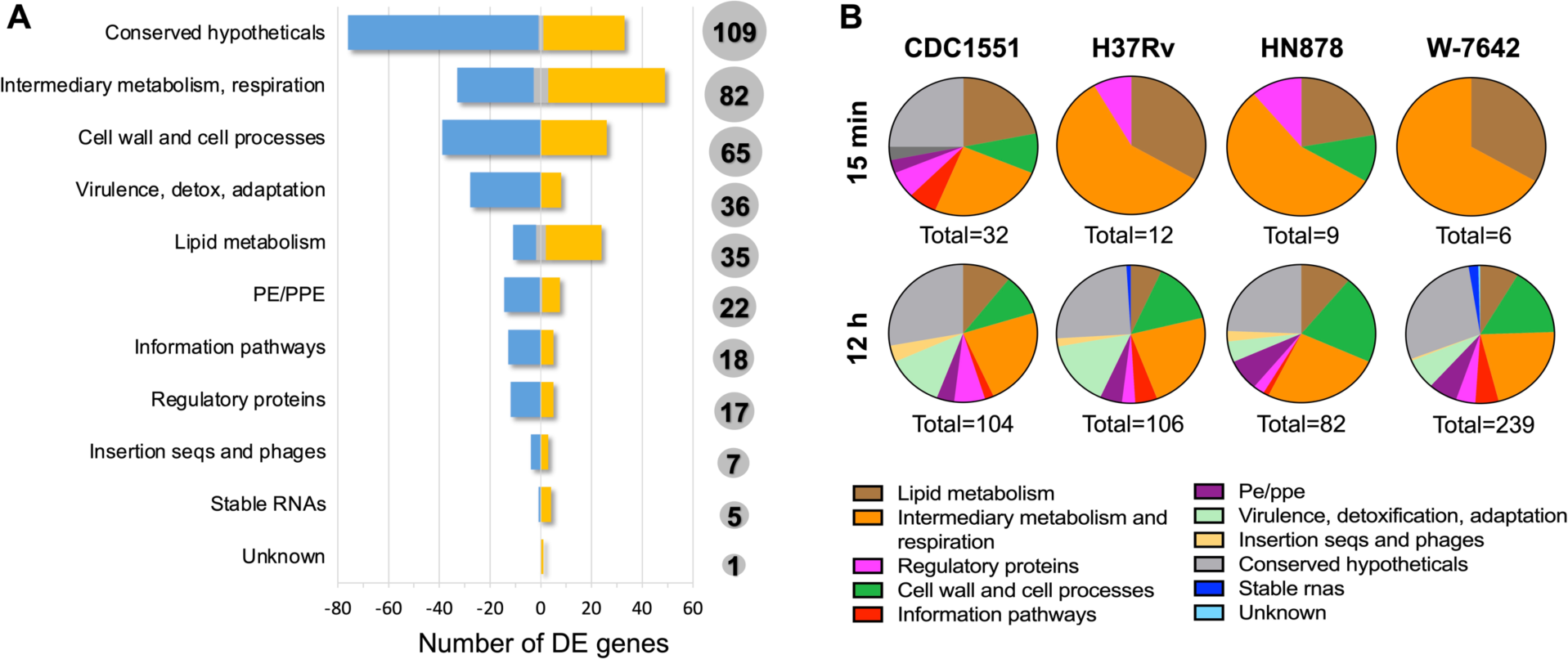
Functional categories of differentially expressed genes in ALF-exposed *vs.* unexposed *M.tb*. **A**) Total number of DEGs (in grey circles) separated by functional category, as defined in Mycobrowser. Genes upregulated in all strains: yellow; downregulated in all strains: blue; up or downregulated depending on the strain: grey. **B**) Functional categories of DEGs separated by strain and ALF exposure time. Graph generated in Graphpad Prism v9.1.1.

We next performed gene enrichment analysis for the 397 total identified DEGs after ALF exposure, and found 17 significantly enriched pathways, including response to environmental pH, fatty acid biosynthetic process and lipid transport and associated beta-ketoacyl synthase activity, acyl transferase activity, cholesterol metabolic process, and mycofactocin and heme activity (**Figure 4**). Other enriched pathways that did not pass the FDR cutoff of 0.05 were sulfolipid metabolic process, protein secretion by type VII secretion system, PPE superfamily, lipid biosynthesis, response to temperature and hypoxia, sigma factor activity, and phenolic phthiocerol biosynthetic process, among others (**Supplemental Table S3**).

**Figure 4.**
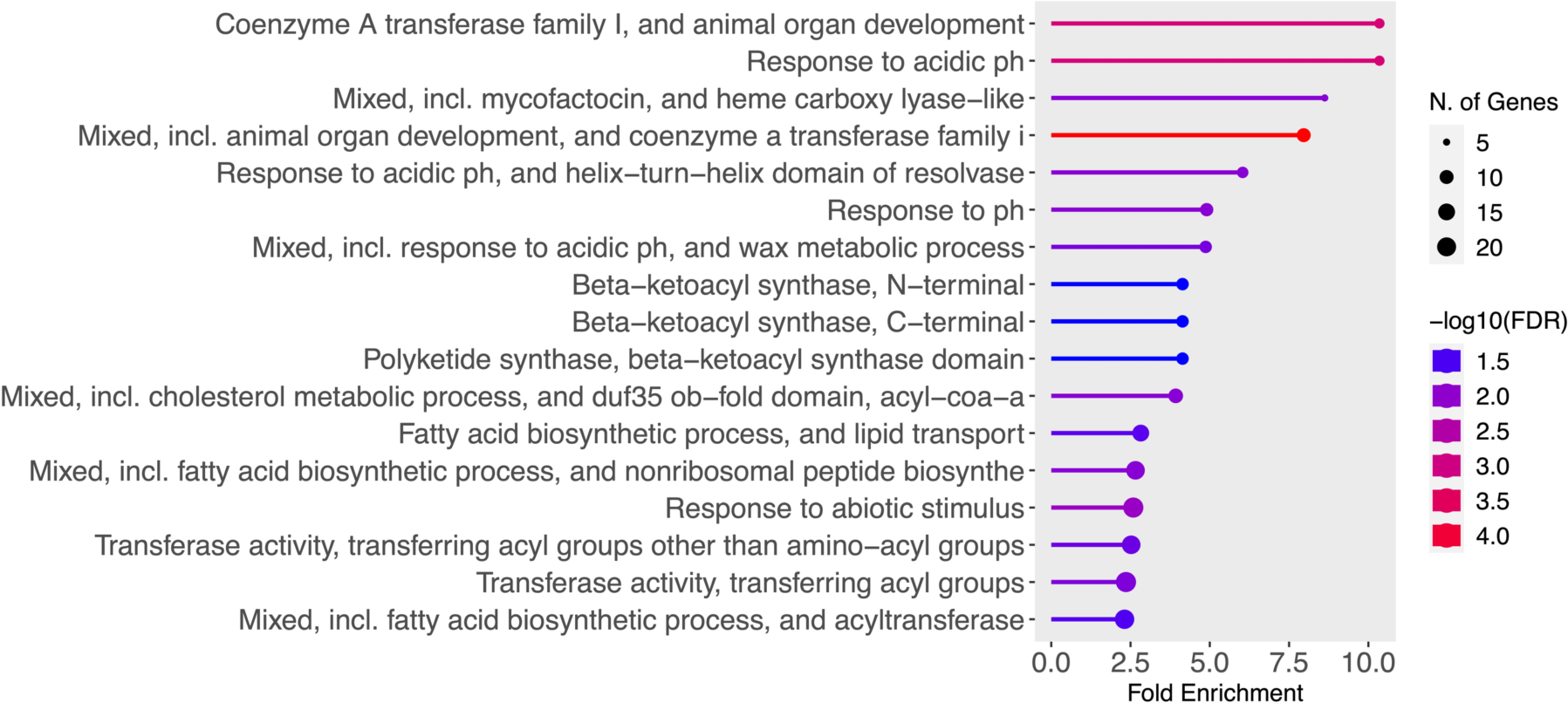
Gene enrichment analysis was performed in ShinyGO 0.77. Significant enriched pathways (FDR <0.05) were sorted by fold enrichment. Graph color depicts significance [-log10 (FDR)] and the number of DEGs belonging to that pathway are shown in proportional circles.

### Exposure to ALF activates the KstR2 regulon associated with cholesterol utilization in *M.tb*

The ability of *M.tb* to utilize host-derived cholesterol as a source of carbon is critical for its growth and virulence during both early (intra-macrophage growth) and late (chronic) stages of infection (Pandey and Sassetti 2008; Griffin et al. 2012; Wipperman et al. 2014). Indeed, deletion of genes associated with cholesterol catabolism result in attenuated *M.tb* strains (Hu et al. 2010; Crowe et al. 2017). However, many molecular aspects of the initiation and maintenance of cholesterol catabolism in *M.tb* remain unknown.

The cholesterol catabolic pathway is highly conserved in mycolic acid-containing actinomycetes, and is encoded by more than 100 genes in *M.tb* (Pawelczyk et al. 2021). There are three distinct sub-pathways or processes during cholesterol catabolism, according to the three parts of the molecule: side chain degradation, A & B ring degradation, and C & D ring degradation (Crowe et al. 2017). Two major regulons, controlled by two TetR-type transcriptional regulators, have been associated with these processes: KstR1 regulates the transcription of side chain and A & B ring degradation genes, while KstR2 controls the transcription of C & D ring degradation genes (Kendall et al. 2010). Other genes though to be regulated by cholesterol belong to the mammalian cell-entry Mce3R regulon, implicated in stress resistance (Santangelo et al. 2009; Yang et al. 2019), and the SigE regulon, essential for virulence (Fontan et al. 2008).

Our results indicate that most KstR2 genes (12/15) were significantly upregulated (Log_2_FC >1 and FDR < 0.1) starting at 15-min post-ALF exposure and maintained at 12-h in all *M.tb* strains tested in this study, independent of their transmissibility, virulence, or antibiotic resistance status, indicating a general ALF-driven response rather than a strain-specific response to ALF (**Figure 5**). This included the genes for C & D ring degradation (3 out of 5 genes were significantly upregulated in at least one of the strains/conditions tested). In contrast, genes associated with the KstR1 regulon, including ATP-dependent cholesterol import genes encoded by the Mce4 transport system (Mohn et al. 2008) and side chain and A&B ring degradation genes, were not differentially expressed when compared to unexposed *M.tb* at any of the ALF-exposure periods of time tested, although there was a trend of slight downregulation for several of the genes at 12-h after ALF exposure (**Figure 5**). Only *ltp3* from the KstR1 regulon, encoding a keto acyl-CoA thiolase involved in the cholesterol side chain degradation (Gilbert et al. 2018), was significantly downregulated in strain CDC1551 at 12-h post-ALF exposure. Other significant DEGs associated with cholesterol side chain degradation were *fadE30*, *fadE32*, and *echA20* (all three upregulated and controlled by KstR2), *fadA6* (upregulated), and *fadB2* (downregulated in all four strains, especially after 12-h of ALF exposure). In addition, of the 23 genes associated with the Mce3R regulon, only two were significantly differentially expressed: *yrbE3B*, encoding a hypothetical membrane protein with phospholipid transport activity, was upregulated in strain HN878 at 12 h post-exposure, while *mce3F* was downregulated in H_37_R_v_ and W-7642 at that same time point (**Figure 5**). None of the cholesterol-associated *sigE* regulon genes were differentially expressed in our dataset under the experimental conditions tested.

**Figure 5.**
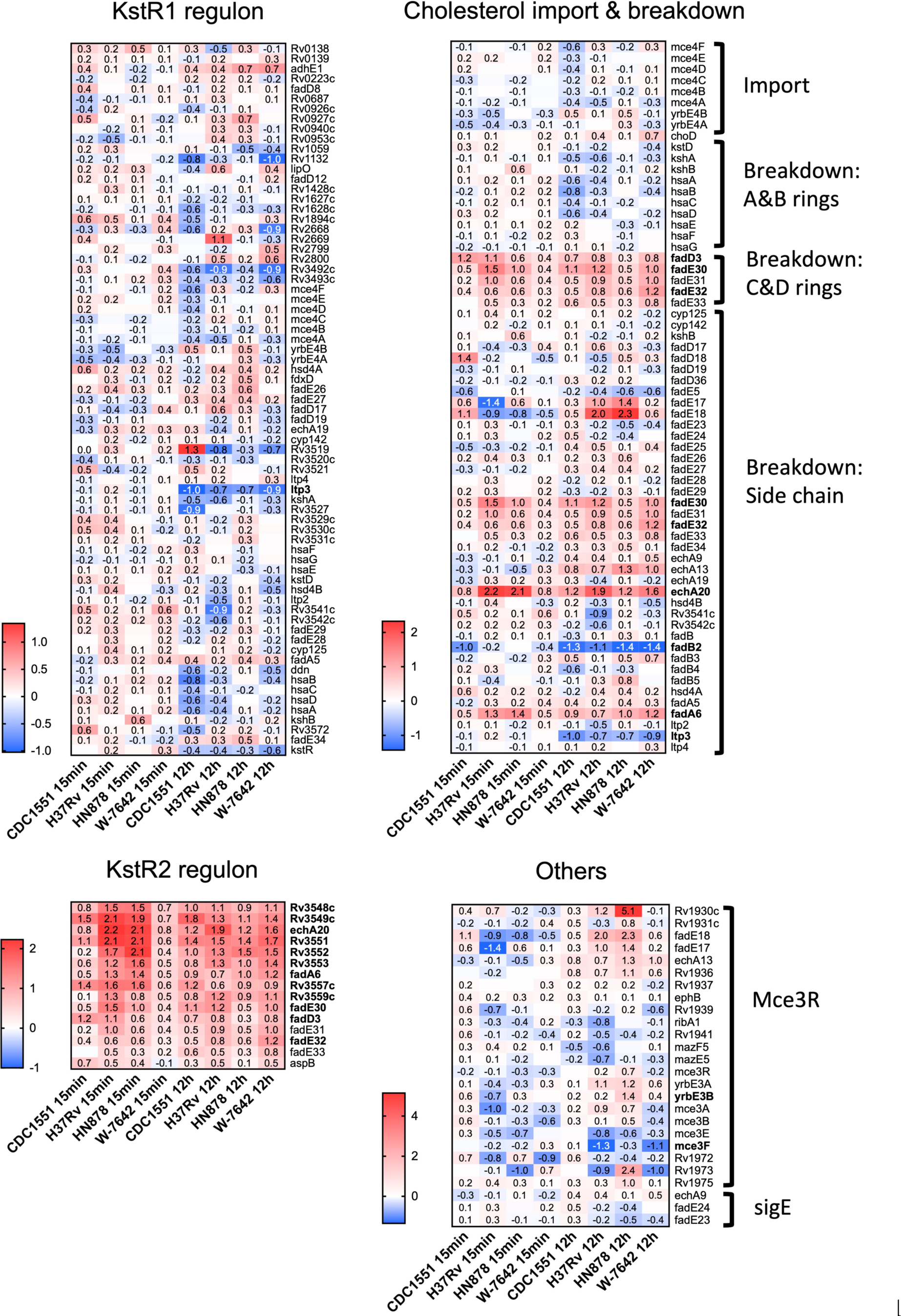
A heatmap of cholesterol utilization genes associated with the KstR1, KstR2, and other regulons (Mce3R and SigE), as well as cholesterol import and breakdown genes was generated in Graphpad Prism v9.1.1. Cells depict log_2_FC values (ALF-exposed *M.tb vs.* unexposed bacteria), upregulated: red, downregulated: blue. Genes in bold indicate significant DEGs (log_2_FC greater than an absolute value of 1, FDR >0.1) in one or more of the strains and/or time points.

### Exposure to ALF shows a temporal adaptation of *M.tb* to the lung environment

In addition to the cholesterol utilization pathways, most *M.tb* strains showed upregulation of genes required for maintaining an appropriate mycolic acid layer composition and permeability of the *M.tb* cell envelope at 15 min post-ALF exposure, achieving statistical significance in strains CDC1551 and W-7642 (**Figure 6**, **Supplemental Table S2**). This included shared genes *Rv3083* (*mymA*), *Rv3084* (*lipR*), and *Rv3085* (*sadH*), as well as *Rv3086* (*adhD*), *Rv3087*, and *Rv3088* (*tgs4*), only significant in W-7642. In contrast, these genes were downregulated after 12 h of ALF exposure (**Figure 7A**). All these genes belong to the *mymA* operon, associated with maintaining the *M.tb* cell envelope under harsh conditions (Singh et al. 2003; Saraav et al. 2017).

**Figure 6.**
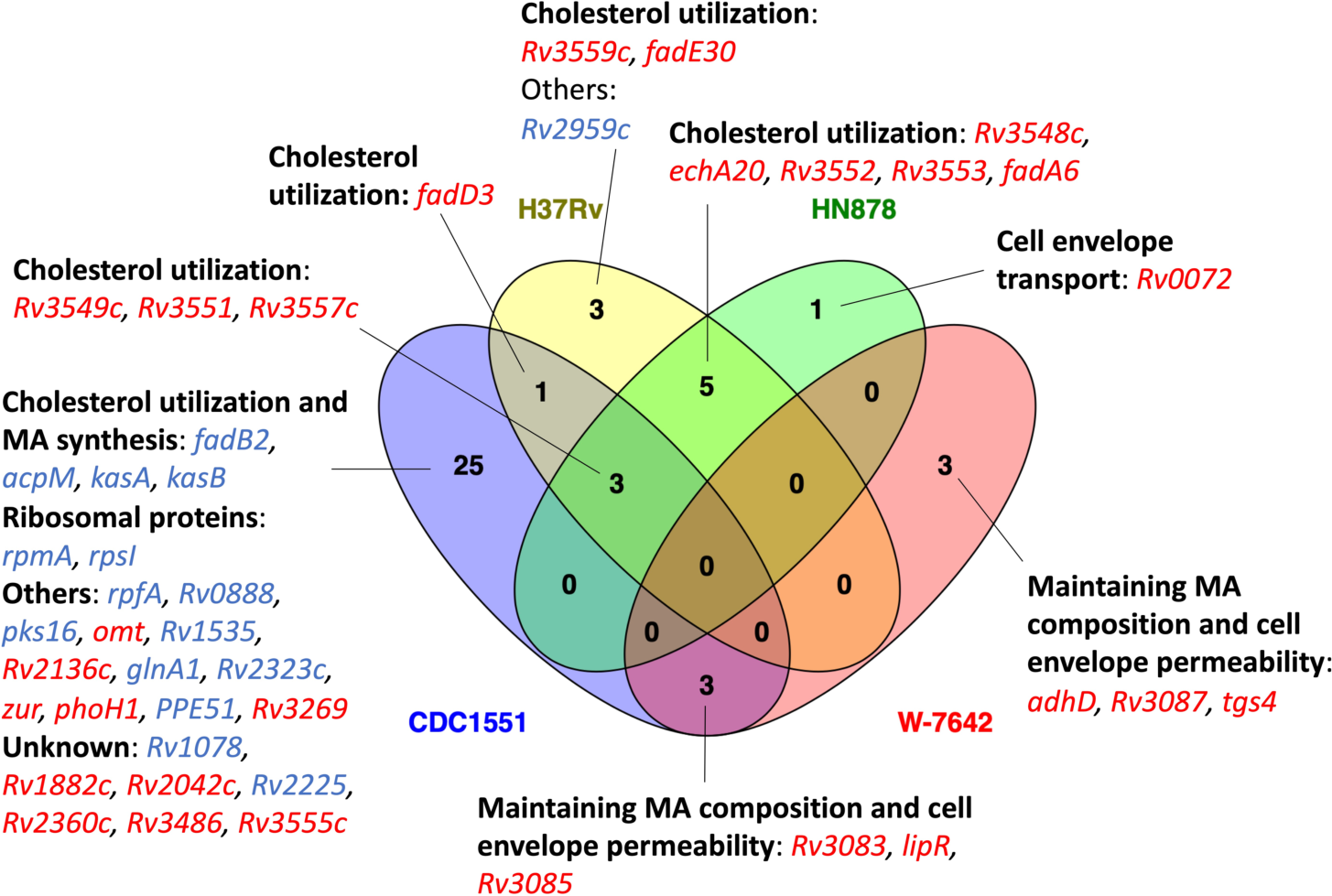
Venn diagram showing shared and strain-specific DEGs (upregulated in red, downregulated in blue) between ALF-exposed and unexposed *M.tb* after 15-min of exposure. Venn diagram was constructed using Venny 2.1.0 (Oliveros 2007).

**Figure 7.**
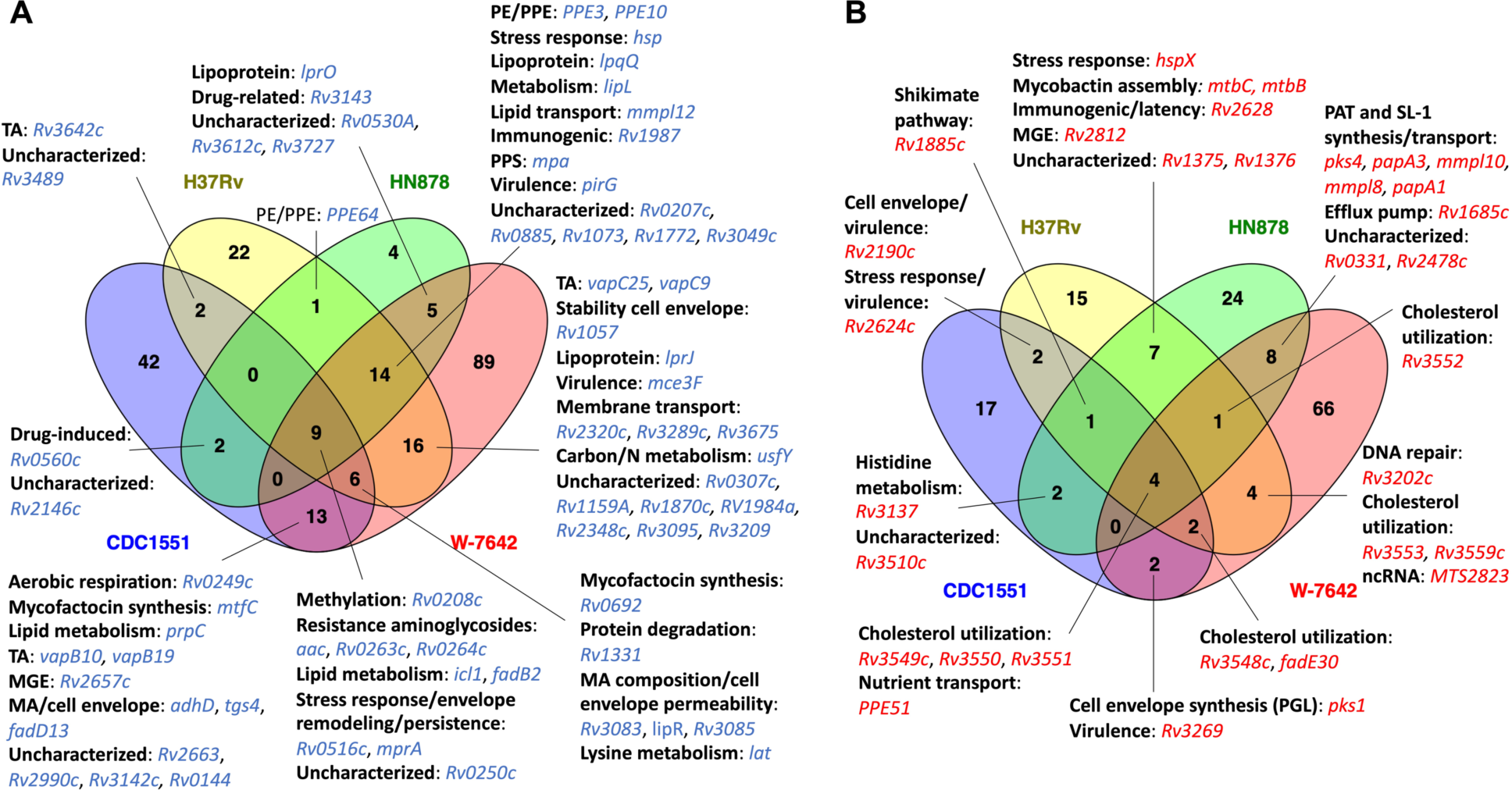
Venn diagram showing shared DEGs (ALF-exposed *vs.* unexposed *M.tb*) among two or more strains at 12-h post-ALF exposure: **A**) downregulated genes (blue); **B**) upregulated genes (red). Venn diagrams were constructed using Venny 2.1.0 (Oliveros 2007).

As mentioned earlier, most of the differential expression happened at 12-h post-ALF exposure. At this time point, we found several up and downregulated genes shared among two or more of the *M.tb* strains evaluated. Interestingly, both hypervirulent HN878 and drug resistant W-7642 showed significant upregulation of diacyltrehalose (DAT), penta-acyltrehalose (PAT) and sulfolipid-1 (SL-1) biosynthesis and transport genes *pks4*, *papA3*, *mmpL10, papA1*, and *mmpL8* (**Figure 7B**). These glycolipids are major *M.tb* cell envelope components (Garcia-Vilanova et al. 2019), where some of them are described as being significantly reduced in content on the *M.tb* cell envelope surface after only 15-min of exposure to human ALF (Arcos et al. 2011), indicating a potential compensatory upregulation of these cell envelope biosynthetic and transport genes to restore these components on the *M.tb* cell surface. Indeed, *pks1*, involved in the synthesis of immunomodulatory phenolic glycolipids (PGLs) produced by some *M.tb* strains and associated with cell envelope impermeability, phagocytosis, defense against oxidative stress and biofilm formation (Pang et al. 2012; Garcia-Vilanova et al. 2019; Ramos et al. 2020), was upregulated in all *M.tb* strains in this study, although was only significant in CDC1551 and W-7642 (**Figure 7B, Supplemental Table S2**). We note here, though, that many clinical isolates (including CDC1551), as well as laboratory strain H_37_R_v_, contain a *pks1-15* polymorphism associated with the inability of these strains to synthesize PGL (Constant et al. 2002).

*Rv1685c*, a TetR-like transcriptional regulator that controls an efflux pump with a potential role in the response to acid nitrosative stress (Perrone et al. 2017), was also significantly expressed in HN878 and W-7642 (**Figure 7B**). Other upregulated genes at 12 h post-ALF exposure were associated to mycobactin assembly (*mtbB*, *mtbC*), involved in iron acquisition, growth in macrophages and virulence (Quadri et al. 1998; Krithika et al. 2006), and stress response gene *hspX* in H_37_R_v_ and HN878, as well as potential immunogenic and virulence-associated genes (*Rv2190c*, *Rv2624c*) in CDC1551 and H_37_R_v_.

We also found several shared downregulated genes after 12 h of ALF exposure (**Figure 7A**). Genes associated with resistance to aminoglycosides (*acc*, *Rv0263c*, *Rv0264c*) (Bassenden et al. 2021), lipid metabolism (*icl1*, *fadB2*), and osmotic stress response and persistence (*Rv0516c*, *mprA*) (Banerjee et al. 2016) were consistently downregulated in all four *M.tb* strains. Further, fatty acid (FA) transporter *mmpL12* was down in all *M.tb* strains but CDC1551. We also observed downregulation of mycofactocin synthesis genes, which have a potential multifaceted role in alcohol metabolism of *M.tb*, hypoxia adaptation, and redox homeostasis (Krishnamoorthy et al. 2021), as well as several lipoproteins and toxin-antitoxin (TA) systems in most of the *M.tb* strains, among others (**Figure 7A**).

### Strain-specific transcriptional profiles after ALF exposure

In addition to shared transcriptional responses among *M.tb* strains due to ALF exposure, we also observed unique gene expression signatures attributed to the *M.tb* genomic background at both time points tested, shown in **Supplemental Table S4**. At 15 min post-ALF exposure, most of the strain-specific DEGs were still related to the KstR2 regulon and maintaining the mycolic acid layer composition and permeability of the *M.tb* cell envelope. We found a gene encoding a probable transmembrane protein responsible for the transport of glutamine across the membrane (*Rv0072*) significantly increased in the hypervirulent HN878 (log_2_FC >5). The highly transmissible CDC1551 showed the most changes at this time point compared to the other strains, with 25 unique DEGs (**Figure 6**). It included downregulation of essential mycolic acid biosynthesis genes *acpM*, *kasA*, and *kasB* (Slayden and Barry 2002), *pks16* involved in biofilm formation (Islam et al. 2012), and cell envelope-associated genes such as *rpfA* (stimulates resuscitation of dormant cells), *Rv0888* (induces formation of neutrophil extracellular traps or NETs) (Dang et al. 2018), and *glnA1* (regulates nitrogen levels during poly-L-glutamine or PLG synthesis in the cell envelope) (Tripathi et al. 2013). Interestingly, a global transcriptional regulator (*Rv2359* or *zur*) that represses transcription of genes involved in zinc homeostasis was upregulated in CDC1551 (**Figure 6**). These data suggest a potentially decreased pathogenicity of CDC1551 when in contact with ALF as little as 15-min.

At 12-h post-ALF exposure, CDC1551 downregulated several cell envelope genes, including *Rv0559c* and *phoY*, with suggested roles in drug resistance (Wang et al. 2013; Gamngoen et al. 2018). Sigma factor E (*sigE*), involved in persistence to anti-TB drugs and essential for virulence and phagosome maturation in macrophages, was downregulated (Casonato et al. 2014; Pisu et al. 2017), as well as *msrB*, which encodes a repair enzyme for proteins that are inactivated by oxidation and reported as upregulated in the resuscitation of dormant *M.tb* (Salina et al. 2019). In addition, several PE/PPE genes, involved in virulence and the activation of proinflammatory responses, and stress-induced genes (*e.g*. *hspR*, *clpB*) were also downregulated. In contrast, only a few unique genes were upregulated in CDC1551, including *Rv2958c* and *Rv2959c*, involved in the glycosylation of cell envelope PGLs, and some fatty acid biosynthesis genes, which were initially downregulated at 15 min ALF post-exposure.

Interestingly, the virulent H_37_R_v_ strain showed upregulation of several latency-associated *dosR* regulon genes after 12-h of ALF exposure (*Rv2623* or *TB31.7*, *Rv2625c*, *hrp1*) (Garcia-Morales et al. 2017), as well as *pncB2*, involved in NAD biosynthesis during host invasion that was found increased during hypoxia (Boshoff et al. 2008), indicating a potential role during persistence (**Supplemental Table S4)**. Gene *irtB*, involved in iron homeostasis (Zhang et al. 2020), and a couple of genes associated with resistance to oxidative stress (*lpdA*, *Rv1050*) were also upregulated. Similar to CDC1551, we found two sigma factors with decreased expression (*sigB* and *sigD*), potentially involved in the regulation of the stationary phase of *M.tb* (Chauhan et al. 2016). Other unique downregulated genes included *pup*, with a role in proteasomal degradation, and TA-associated genes (**Supplemental Table S4)**.

Of the 28 unique DEGs in hypervirulent HN878 at 12-h post-ALF exposure, 24 showed significant increased expression (log_2_FC >1) (**Supplemental Table S4)**. Within the cell envelope functional category, we found *esxI* upregulated, which encodes an ESAT-6 protein secreted by the ESX-5 system that is involved in the virulence of *M.tb* (Shah et al. 2015). We also found upregulated the lipoprotein *lppJ* of unknown function, and the cluster *Rv1686c/Rv1687c*, a putative efflux pump responsible for the transport of undetermined substrates (potentially drugs) across the membrane that is activated under several environmental stressors (Birhanu et al. 2022). Further, all unique intermediate metabolism and respiration genes were upregulated in HN878, including *glbN* that encodes an oxygen transport protein upregulated during macrophage infection (Sethi et al. 2016), molybdenum cofactor biosynthesis protein MoaA1, strongly induced under hypoxia (Levillain et al. 2017), and *dxs2*, involved in the mycobacterial ability to prevent acidification of the phagosome (Pethe et al. 2004). Contrary to CDC1551 and H_37_R_v_, in HN878 two PE/PPE genes showed increased expression. The only unique downregulated genes in HN878 were two transmembrane proteins of unknown function, an uncharacterized hypothetical protein, and *fadE8*, involved in lipid degradation (**Supplemental Table S4)**.

Finally, MDR-*M.tb* W-7642 showed the most strain-specific DEGs (#155) after 12 h post-ALF exposure compared to unexposed bacteria, with 66 upregulated and 89 downregulated genes (**Figure 7**). We found upregulation of several genes associated with the ESX-3 system and other genes related to iron acquisition and utilization (*esxH*, *eccD3*, *mycP3*, *Rv0285* or *PE5*, *hemE*, *hemL*, *Rv2047c*), essential for mycobacterial growth *in vitro* and virulence (Tufariello et al. 2016) (**Supplemental Table S4)**. In contrast, ESX-1 genes *espD*, *espC*, and *espA* were downregulated. The ESX-1 system secretes host membrane-targeting proteins such as ESAT-6 that interrupt the innate immune response against *M.tb* (Wong 2017). Other ESAT-6-like proteins were found downregulated (EsxJ, EsxP, EsxD) in W-7642, with the exception of EsxL that was upregulated. Overall, genes generally induced during starvation, persistence, and/or during the stationary phase of *M.tb* showed decreased expression (*e.g. rsbW*, *vapC44*, *mce1R*, *lipU, Rv3241c*) (Hampshire et al. 2004), as well as sigma factors *sigF* and *sigI* (**Supplemental Table S4)**. However, we observed increased expression of *M.tb* cell envelope biosynthesis genes and lipid metabolism (*Rv2974c, aftD, murE, murF, Rv3807c, accD6, Rv3031, kasB, acpS, fas, ppsC, ppsD, pks15*), amino acid metabolism (*leuC, leuD, argD, trpA*), and DNA repair genes (*ruvB, Rv3201c, Rv3433c, radA*). Interestingly, W-7642 was the only strain with decreased expression of *ethA* (Dover et al. 2007), but increased expression of *Rv2989* (Li 2016), both associated with drug resistance. Other unique DEGs in strain W-7642 are shown in **Supplemental Table S4**.

RNA-seq results for selected genes of interest discussed in the results section were validated by qPCR, showing similar fold change values (**Supplemental Table S5**, **Supplemental Figure S3**).

### Exposure to ALF results in a slight increase of *M.tb* intracellular growth in primary human macrophages

To determine if exposure to ALF had any short-term effects on *M.tb* uptake and intracellular growth, we infected human primary monocyte-derived macrophages (MDMs) with 4 strains of *M.tb* (CDC1551, H_37_R_v_, HN878, and W-7642) previously exposed for 12-h to our ALF pool (or unexposed), and calculated the bacterial growth rate over a 24-h period. We choose to infect MDMs with *M.tb* after 12-h of ALF exposure since it was the exposure time that resulted in the most transcriptional changes in all strains tested. We observed an increase of up to 53% in the uptake of ALF-exposed *M.tb* compared to unexposed bacteria, with the exception of CDC1551 that showed a 41% reduction (**Figure 8A**). In addition, there was a trend of increased growth rate in all strains after ALF exposure compared to unexposed bacteria (**Figure 8B**). These results suggest that the ALF environment is priming and promoting metabolic changes in *M.tb*, affecting bacterial uptake by human macrophages and with potential consequences for *M.tb* intracellular growth. Further studies are needed to determine if these changes have long-term consequences during infection *in vivo*.

**Figure 8.**
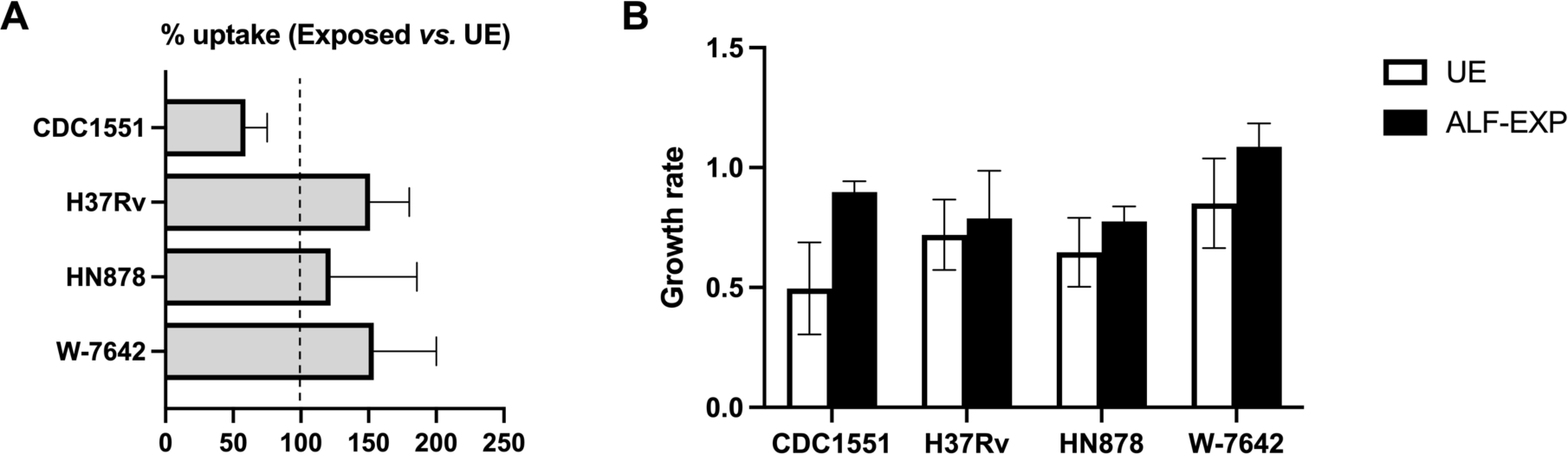
*In vitro* infection of human macrophages: **A**) % uptake of ALF-exposed *M.tb vs*. unexposed *M.tb* was calculated for each of the strains and plotted in Graphpad Prism v9.1.1. Uptake was defined as CFUs 2-h post infection (hpi) *vs.* CFUs inoculum. Data are represented as the mean ± SEM of n = 3 independent experiments (3 human donors). Values of > 100% indicate an increased uptake in ALF-exposed *M.tb* compared to unexposed (UE) *M.tb*, while values < 100% indicate decreased uptake. **B**) Growth rate between 2 and 24 hpi in ALF-exposed (ALF-EXP) and unexposed (UE) *M.tb* was calculated for each of the strains as in (Scordo et al. 2019). Graph was generated in Graphpad Prism v9.1.1 and plotted as mean ± SEM of n = 3 independent experiments (3 human donors).

## CONCLUSIONS

The continuous development of Next Generation Sequencing (NGS) technologies in the past 15 to 20 years has greatly contributed to an increased knowledge of host-pathogen interactions, including in the TB field. Genome-wide transcriptomic studies are critical in deciphering molecular determinants of TB disease progression and development of drug resistance in response to particular drug treatments, defining bacterial interactions with the host immune system, or studying the *M.tb* adaptation to different lung microenvironments, among others (Tang et al. 2020; Dow et al. 2021; Tabone et al. 2021).

Our published studies on the role of ALF during *M.tb* infection indicate that human ALF hydrolases alter the *M.tb* cell envelope. These alterations have an impact in subsequent interactions with immune host cells *in vitro* and *in vivo* (Arcos et al. 2011; Arcos et al. 2015; Arcos et al. 2017; Scordo et al. 2017; Moliva et al. 2019; Scordo et al. 2019; Olmo-Fontánez et al. 2021), suggesting that physiological levels of ALF plays an important role in shaping the *M.tb* cell envelope during the very early stages of infection. Here, we used RNA-seq to depict the transcriptional responses to ALF exposure of four *M.tb* strains (CDC1551, H_37_R_v_, HN878, and W-7642) with different degrees of transmissibility, virulence, and drug resistance. Our results define for the first time the overall impact of human ALF, first lung microenvironment encountered by *M.tb* during the establishment of the infection, on the adaptation of this pathogen to the human host prior to infection of host cells in the alveolar space. We also determined whether exposure to ALF has different transcriptional effects based on the *M.tb* genomic background, which might further define phenotypic differences among *M.tb* strains. Indeed, we observed temporal shared and strain-specific transcriptional responses in *M.tb* after exposure to human ALF for different lengths of time, demonstrating that *M.tb* can metabolically adapt to this lung microenvironment prior to encountering and infecting host cells.

The most striking effect of ALF exposure shared by all *M.tb* strains studied was the activation of cholesterol utilization pathways, specifically the KstR2 regulon, induced at 15-min post-ALF exposure and maintained at 12-h (**Figure 5**). The KstR2 regulon includes genes for C & D ring degradation, as well as some genes associated with degradation of the cholesterol side chain (*e.g. fadE30*, *fadE32*, and *echA20*) (Wipperman et al. 2014). Conversely, expression of genes under the KstR1 regulon remained practically unaltered after exposure to ALF, as well as most of the genes belonging to the Mce3R and SigE regulons (**Figure 5**), also associated to cholesterol utilization. Priming of cholesterol uptake and utilization is essential for survival of *M.tb* in the human host, especially in the nutrient-deprived macrophage environment, where cholesterol significantly contributes to *M.tb* intracellular growth and pathogenesis (Brzostek et al. 2009; Pawelczyk et al. 2021). Indeed, metabolites originated from cholesterol degradation such as pyruvate, acetyl-CoA or succinyl-CoA, among others, can be incorporated into *M.tb* metabolic pathways for cell envelope lipid biosynthesis and production of energy. Further, *M.tb* uses a cholesterol-dependent pathway to infect host cells, induces a foamy macrophage phenotype, and extracts cholesterol and lipids from caseous granulomas to persist, demonstrating the importance of host cholesterol utilization during both early and latent stages of the infection (Pandey and Sassetti 2008; Peyron et al. 2008). Our data suggest that the metabolic adaptation of *M.tb* to use host cholesterol is initiated even before its intracellular stage in host cells, where extracellular *M.tb* adapts its metabolism and starts consuming host cholesterol present in the ALF, activating cholesterol degradation pathways that will later be key for infection and survival within host cells. Indeed, we observed a trend of increased intracellular growth rate the first 24 hpi after *M.tb* was exposed to ALF for 12-h in all *M.tb* strains tested (**Figure 8B**).

We also found changes in the expression of genes associated to cell envelope biosynthesis and remodeling in ALF-exposed *M.tb* when compared to unexposed bacteria. In fact, *M.tb* exposure to human ALF alters the *M.tb* cell envelope after solely 15-min, by reducing, for example, the amount of two major *M.tb* cell surface virulence factors ManLAM (by ∼80%) and TDM (by ∼40%), both implicated in promoting phagosome maturation arrest and *M.tb* intracellular growth (Arcos et al. 2011; Torrelles and Schlesinger 2017; Garcia-Vilanova et al. 2019). In response to this reduction, *M.tb* exposed to ALF upregulates genes involved in the synthesis of TDM and mannose-containing biomolecules (e.g. ManLAM) (Allue-Guardia et al. 2022). However, although these cell envelope biosynthesis genes are quickly upregulated by ALF exposure and their restoration might be associated with the increased uptake and intracellular growth rate trend observed for most of the strains tested, their lasting effects of replenishing TDM and ManLAM on the *M.tb* surface are still uncertain.

Further, most *M.tb* strains exposed to ALF showed increased expression of some of the genes in the *mymA* operon (*Rv3083-Rv3089*, **Supplemental Table S2**) after 15-min of contact with ALF (**Figure 6)**. This operon is induced when *M.tb* is exposed to acidic pH and during macrophage infection, demonstrating a role in the modification of fatty acids required for maintaining the appropriate permeability of the cell envelope and mycolic acid layer composition under harsh conditions (Singh et al. 2003; Saraav et al. 2017). Furthermore, *mymA* mutants are more susceptible to known anti-TB drugs and detergents, and infection with those mutants results in decreased bacterial load in the spleen of guinea pigs, showing that this operon is also important for *M.tb* pathogenesis and drug resistance (Singh et al. 2005). This could explain why MDR W-7642 was the only strain in our dataset that had significantly increased expression of all genes from the *mymA* operon after 15-min exposure to ALF. In contrast, *mymA* genes were downregulated at 12-h post-ALF exposure (**Figure 7A**). These results suggest that, early after contact, *M.tb* upregulates genes that help maintain the cell envelope integrity to compensate for the changes generated upon contact with ALF, but this effect is only temporal and expression of these genes is decreased after a certain period, maybe due to a certain degree of *M.tb* adaptation to the ALF environment. What we known is that ALF exposure to *M.tb* does not affect bacterial viability (Arcos et al. 2011; Scordo et al. 2019).

Several other cell envelope-associated genes were significantly downregulated after exposure to ALF for 12-h, including multiple protein export genes in CDC1551 (**Supplemental Table S2**) and lipoproteins (*lprJ, lpqQ, lprO, Rv3354, Rv2293c*) in H_37_R_v_, HN878, and especially W-7642. Lipoproteins are membrane-anchored proteins involved in virulence and immunity in *M.tb*, with a dual role, some promote survival of *M.tb*, while others induce host protective immune responses (Becker and Sander 2016). The ones found here are some of the less studied, and their specific function/s are still unknown, although our MDM infection data show a slight increase in intracellular growth when *M.tb* is exposed to ALF (**Figure 8B**), which could be explained by the downregulation of lipoproteins that potentially induce immune responses in the host. In contrast, only hypervirulent HN878 and MDR W-7642 strains, both from the Beijing lineage, showed significantly increased expression of DAT/PAT, and SL-1 biosynthesis and transport genes, including MmpL proteins (*mmpL10* and *mmpL8*) important for the physiology and pathogenesis of *M.tb* (**Figure 7B**) (Melly and Purdy 2019). Together with ManLAM and TDM, these cell envelope glycolipids have the ability to modulate host cell immune responses (Zhang et al. 1991), as well as blocking phagosome maturation (e.g. fusion with lysosomes) in infected phagocytes (Brodin et al. 2010; Gilmore et al. 2012), supporting the hypervirulent phenotype found in most Beijing lineage strains.

Further, other genes involved in the biosynthesis of the *M.tb* cell envelope were significantly upregulated in W-7642, such as *pks1* and *pks15* (PGL synthesis) (Ramos et al. 2020), *murE* and *murF* (peptidoglycan synthesis) (Maitra et al. 2019), *aftD* (biosynthesis of the arabinogalactan region of the *M.tb* cell envelope backbone known as mycolyl-arabinogalactan-peptidoglycan complex or mAGP) (Skovierova et al. 2009), *Rv2974c* (hypothetical protein potentially involved in glycerolipid and glycerophospholipid metabolism) (Vargas et al. 2021), and several lipid and amino acid metabolic genes (**Supplemental Table S2**) implicated in cell envelope biomass synthesis (Maitra et al. 2019). These results indicate that, even though exposure to functional ALF cleaves components of the cell envelope (Arcos et al. 2011), strains HN878 and W-7642 are capable of reprograming their metabolism and upregulating genes involved in the synthesis of cell surface components involved in *M.tb* survival and pathogenesis (DAT/PAT and SL-1).

Further, W-7642 was the only strain with decreased expression of *ethA*, which encodes the enzyme responsible for the activation of the pro-drug ethionamide (Dover et al. 2007), but increased expression of *Rv2989*, an IclR family transcriptional regulator involved in increased isoniazid tolerance (Li 2016). Thus, exposure to ALF might be further promoting the drug resistance ability of W-7642.

ESX or type VII secretion systems, responsible for the secretion of *M.tb* proteins to evade host immune responses (Groschel et al. 2016), were also impacted after 12 h of ALF exposure, especially in MDR W-7642. In this strain, we found significantly increased expression of ESX-3 genes, involved in iron and nutrient uptake and utilization (Tufariello et al. 2016), as well as other iron homeostasis genes (e.g. *mtbB*, *mtbC*) important for intracellular growth that were also upregulated in the rest of the strains. In contrast, genes from the ESX-1 system, involved in the lysis of the phagosomal membrane and phagosomal escape of the bacteria (Conrad et al. 2017), were downregulated in W-7642. Increase of ESX-3 and other iron homeostasis genes after human ALF exposure could be preparing *M.tb* for subsequent infection of macrophages by improving its ability to survive and grow in phagosomes in detriment of other functions (e.g. the ability to escape the phagolysosome), as suggested by a moderate, although not significant, increase in intracellular growth in MDMs after 24 hpi (**Figure 8B**). Associated with the ESX-5 secretion system, gene *esxI* (Bukka et al. 2011), suggested to modulate growth in intracellular bacteria, was uniquely upregulated in HN878, potentially related to its hypervirulent genotype.

Several sigma factors (*sigB, sigD, sigE, sigF, sigI*) were found downregulated after ALF exposure in all *M.tb* strains studied (significant in all but HN878). Sigma factors are known to contribute to *M.tb* pathogenesis by regulating the expression of genes involved in the adaptation of *M.tb* to stress conditions encountered during different stages of infection (Sachdeva et al. 2010), including the stationary phase. Accordingly, the majority of stress response genes detected after DE analysis, as well as genes associated to hypoxia and latency, were downregulated in most strains. H_37_R_v_ was the exception, showing upregulation of genes with potential roles during persistence and oxidative stress (*e.g. TB31.7*, *Rv2625c*, *hrp1, hspX, pncB2, lpdA*) after exposure to ALF for 12-h (**Supplemental Table S4)**. Overall, this suggests that exposure to ALF might be reducing the ability of some *M.tb* strains (CDC1551, HN878, W-7642) to adapt to long-term survival and persist in a latent state, which needs further investigation.

Finally, all DEGs from the PE/PPE functional category were downregulated in both CDC1551 and H_37_R_v_ after 12-h of exposure to ALF, while some were upregulated in HN878 and W-7642 (Beijing lineage strains) (**Supplemental Table S2**). While most PE/PPE proteins are still uncharacterized and their molecular mechanisms are still unknown, it is suggested that these are involved in *M.tb* virulence and modulation of host immune responses (Ates 2020). The fact that after exposure to ALF some of these are upregulated only in Beijing lineage strains could indicate a potential adaptive advantage of HN878 and W-7642 to the human host. Additional mechanistic studies on these particular PE/PPE proteins are needed to elucidate their exact function during infection. Conversely, TA systems, associated to *M.tb* pathogenicity and persistence (Yu et al. 2020), were downregulated in most of the *M.tb* strains at 12 h post-ALF exposure, which might be explained by the ALF trying to induce a protective response (Arcos et al. 2011).

In summary, this study shows that human ALF changes the metabolism of *M.tb* in a temporal and strain-specific manner, and that it is an important environment to consider when studying *M.tb* infection and pathogenesis. Most of the effects were observed at 12-h ALF post-exposure, with the exception of cholesterol utilization pathways that were rapidly upregulated starting at 15-min after exposure, defining the very early initial status of *M.tb* in the alveolar compartment prior to getting in contact with host cells. Early exposure (15-min) to ALF also drives upregulation of genes associated with maintaining the appropriate composition of the cell envelope. In addition, our transcriptional analyses show remodeling of the *M.tb* cell envelope, upregulation of genes associated with nutrient and iron uptake (e.g. ESX-3), and downregulation of sigma factors and other genes associated with persistence and pathogenesis. These results indicate that the ALF environment promotes changes in *M.tb*’s metabolism, preparing the bacteria for its encounter with host cells (e.g. by upregulating cholesterol utilization pathways and iron/nutrient uptake genes, key for intracellular survival), while at the same time *M.tb* is trying to compensate for ALF-driven changes on the cell envelope early after exposure. This *M.tb*-ALF interplay results in a slight increase in uptake as well as early (24 hpi) *M.tb* growth rate in macrophages in contrast to our previous findings, where *M.tb* exposure to ALF drives a decreased intracellular bacterial growth and slower replication *in vitro* and *in vivo* (Arcos et al. 2011; Arcos et al. 2015; Arcos et al. 2017; Scordo et al. 2017; Moliva et al. 2019; Scordo et al. 2019; Olmo-Fontánez et al. 2021). These differences could be explained by the fact that we are using unexposed bacteria as control (instead of NaCl-exposed *M.tb* used in previous studies, which can cause osmotic stress in the bacteria and subsequent transcriptional changes (Hatzios et al. 2013; Rebollo-Ramirez and Larrouy-Maumus 2019)) and that we are exposing *M.tb* to a pool of healthy adult ALFs instead of individual ALFs from different human sub-populations (e.g. elderly ALF), as well as using other *M.tb* strains rather than Erdman. In this regard, we observed significant differences within strains upon human ALF exposure. Indeed, Beijing lineage strains HN878 and W-7642 upregulated genes associated with the synthesis of cell envelope glycolipids, potentially linked to their hypervirulent phenotype (Reed et al. 2007; Huet et al. 2009), although there was only a slight increase in the growth rate of these two strains 24 hpi after ALF exposure compared to unexposed bacteria, similar to the other strains tested.

This study provides new insights into specific *M.tb* metabolic pathways affected by exposure to human ALF and further defines transcriptional differences among *M.tb* strains exposed to ALF, highlighting the importance of the host lung environment and its interactions with *M.tb* during the early infection process, potentially playing a major role in *M.tb* infection and pathogenesis outcomes.

## METHODS

### Human subjects and ethics statement

Blood and bronchoalveolar lavage fluid (BALF) was collected from healthy adult volunteers, in strict accordance with the US Code of Federal Regulations and approved Local Regulations (The Ohio State University Human Subjects IRB numbers 2008H0135 & 2008H0119 and Texas Biomedical Research Institute/UT-Health San Antonio Human Subjects IRB numbers HSC20170667H & HSC20170315H), and Good Clinical Practice as approved by the National Institutes of Health (NIAID/DMID branch), with written informed consent from all human subjects. Healthy adult volunteers (18–45 years old) were TST- or IGRA-negative, and were recruited from both sexes (50:50 male:female ratio) with no discrimination based on race or ethnicity. The following comorbidities were excluded: drug and alcohol users, smokers, asthma, acute pneumonia, upper/lower respiratory tract infections, acute illness/chronic condition, heart disease, diabetes, obesity, obstructive pulmonary disease (COPD), renal failure, liver failure, hepatitis, thyroid disease, rheumatoid arthritis, immunosuppression or taking nonsteroidal anti-inflammatory agents, human immunodeficiency virus (HIV)/AIDS, cancer requiring chemotherapy, leukemia/lymphoma, seizure history, blood disorders, depression, lidocaine allergies (used during the BAL procedure), pregnancy, nontuberculous mycobacterial infection, and TB.

### Collection of BALF and ALF isolation

BALF was collected from healthy adult donors in sterile endotoxin-free saline (0.9% NaCl), filtered through 0.2 μm filters, and concentrated 20-fold using Amicon Ultra Centrifugal Filter Units with a 10-kDa molecular mass cutoff membrane (Millipore Sigma, Burlington, MA, USA) to obtain the ALF physiological concentration within the lung (1 mg/mL of phospholipid), as we described (Arcos et al. 2011; Moliva et al. 2014; Arcos et al. 2015; Arcos et al. 2017; Scordo et al. 2017; Moliva et al. 2019; Scordo et al. 2019; Olmo-Fontánez et al. 2021; Allue-Guardia et al. 2022). ALF was aliquoted and stored at −80 °C until further use.

### Bacterial cultures and ALF exposure

*M.tb* strain selection was based on lineage, levels of transmissibility, virulence, and drug resistance. Both H_37_R_v_ and CDC1551 belong to lineage 4 (European, American, and African). H_37_R_v_ is a common reference laboratory strain that was derived from a patient with chronic pulmonary TB at the beginning of the 20^th^ century and selected for its virulence in animal models (Steenken et al. 1934). CDC1551 is a clinical strain isolated from a TB outbreak from 1994 to 1996 in a small rural area close to the Kentucky-Tennessee border. CDC1551 is considered of low virulence in humans but highly transmissible, since 72% of both close and casual contacts were infected after a brief exposure with the three original TB patients (Valway et al. 1998). Though highly immunogenic at early stages of infection, CDC1551 is efficiently controlled in the lungs of infected animals (Subbian et al. 2013), contrary to the hypervirulent isolate HN878 (Lineage 2/Beijing/East-Asian), associated with rapid growth and reduced survival due to a decreased Th1 host protective immunity that results in progressive cavitary disease (Subbian et al. 2011). The HN878 phenotype, which was isolated during a TB outbreak in Houston, Texas from 1995 to 1998, is associated with the presence of PGLs and triglycerides in the *M.tb* cell envelope (Reed et al. 2004; Reed et al. 2007). Finally, W-7642, which also belongs to lineage 2, was isolated in the early 1990s from a series of outbreaks in New York City caused by the spread of a multidrug resistant (MDR)-*M.tb* Beijing clone. Besides the MDR phenotype, it is also considered highly virulent, since it caused increased mortality rates compared to other clinical isolates (Bifani et al. 1996).

*M.tb* strains CDC1551, H_37_R_v_ (ATCC# 25618), HN878 (BEI Resources, NR-13647), and W-7642 were cultured from frozen stocks in 7H11 agar plates (BD BBL), supplemented with oleic acid, albumin, dextrose and catalase (OADC) for 14 days at 37°C. Single bacterial suspensions (∼ 1 x 10^9^ bacteria/ml) were obtained as described (Arcos et al. 2011; Arcos et al. 2015; Arcos et al. 2017; Scordo et al. 2017; Moliva et al. 2019; Scordo et al. 2019; Olmo-Fontánez et al. 2021; Allue-Guardia et al. 2022), and further exposed to a pool of ALFs (n=17 different donors, 50:50 male:female ratio, no discrimination in ethnicity and race) for 15 min or ∼12 h at 37°C. Using a pool instead of individual ALFs allows us to reduce human variability and focus on the overall effect of ALF in each *M.tb* strain rather than changes due to ALF-specific factors. After exposure, ALF was washed away and bacterial pellet incubated in RNAProtect (Qiagen, Hilden, Germany) for 10 min at room temperature (RT), centrifuged at 13,000 × *g* and stored at −80 °C until further use. For each of the strains, unexposed bacteria were used as controls and treated using the same conditions as the ALF-exposed *M.tb*.

### RNA extraction, library prep and RNA sequencing

Stored bacterial pellets were used for RNA extraction using the Quick-RNA Fungal/Bacterial Miniprep kit (Zymo Research, Irvine, CA, USA), following the manufacturer’s protocol. Briefly, bacterial pellet was resuspended in lysis buffer and transferred to a ZR BashingBead Lysis tube containing 0.1 mm and 0.5 mm ceramic beads. A bead beating procedure to break the mycobacterial cell envelope was performed in a BioSpec Mini-Beadbeater-24 (6 cycles of 45 s at 3,800 rpm, with 3 min intervals on a cooler rack). RNA in the supernatant was isolated using Zymo-spin columns, with an in-column DNAse I treatment, and eluted in nuclease-free water. A second DNAse treatment was performed on the isolated RNA using the TURBO DNAse reagent (Thermo Fisher Scientific, Waltham, MA, USA), and RNA was further purified using the RNA Clean & Concentrator kit (Zymo Research, Irvine, CA, USA). RNA final concentration was measured with a Qubit 4 Fluorometer using the HS RNA kit (Thermo Fisher Scientific, Waltham, MA, USA), and RIN numbers were obtained using the 4200 TapeStation System (Agilent, Santa Clara, CA, USA) to determine RNA quality before sequencing. A stranded whole transcriptome RNA library was constructed using the Zymo-Seq RiboFree Total RNA kit (Zymo Research) following the manufacturer’s guidelines. RNA (500 ng) was used as input to construct the library, which included a rRNA depletion step of 30 min followed by 14 cycles of PCR. A 75-bp paired-end (PE) sequencing was conducted using a NextSeq 500 Mid Output Illumina platform, obtaining 5-8 million PE reads per sample. We conducted two independent experiments for each of the strains, and all samples were sequenced in the same flow cell to avoid sequencing batch effects.

### Data analysis

Raw reads were trimmed (phred score of 30) and aligned to the reference genome H_37_R_v_ (Genbank NC_000962.3) (Cole et al. 1998; Camus et al. 2002; Lew et al. 2011) using STAR v2.5.3a with default values in the Partek Flow Genomic Analysis software (v9.0.20.0202, Partek Inc., Chesterfield, MO, USA). Transcripts were normalized and quantified as Transcripts per Kilobase Million (TPM) using the Partek E/M (Expectation/Maximization) algorithm. For further analyses, read count data was uploaded to the iDEP.94 software (http://bioinformatics.sdstate.edu/idep94/) (Ge et al. 2018). Raw data were filtered (0.5 CPM, n=1 library) to remove lowly expressed genes and transformed with EdgeR as log_2_ (CPM + c) using default settings (Robinson et al. 2010; McCarthy et al. 2012; Chen et al. 2016), and used for clustering and principal component Analysis (PCA). Hierarchical clustering of normalized gene expression levels was calculated using default values (correlation distance, average linkage, cutoff Z score of 4), and shown as a heatmap. Significant differentially expressed genes (DEGs) between ALF-exposed *M.tb* and unexposed *M.tb* (control) for each of the strains and time points were identified using the DESeq2 method (Love et al. 2014) with an FDR cutoff of 0.1 and a minimum fold change (FC) of 2 (corresponding to a log_2_ FC equal or greater than an absolute value of 1). The Benjamini-Hochberg procedure was used to adjust the false discovery rate (FDR), and fold changes were calculated using an Empirical Bayes shrinkage approach (Love et al. 2014). Functional categories for DEGs were extracted from Mycobrowser (Kapopoulou et al. 2011). Gene enrichment analysis was performed in ShinyGO 0.77 using default settings (FDR cutoff of 0.05 and minimum pathway size of 2, Pathway database: all available gene sets) (Ge et al. 2020). A summary of the experimental conditions and workflow used for RNA sequencing and data analysis is shown in **Supplemental Figure S1**.

### Validation of gene expression by RT-qPCR

Some of the genes that were differentially expressed were selected for validation using RT-qPCR (**Supplemental Table S5**). Briefly, 500 ng of RNA were used for the synthesis of cDNA using the RevertAid H Minus First Strand cDNA Synthesis kit (Thermo Fisher Scientific, Waltham, MA, USA) with random hexamer primers, following the manufacturer’s instructions. Then, cDNA and specific primers were used in a 20 μl qPCR reaction with PowerUp SYBR Green Master Mix (Applied Biosystems, Waltham, MA, USA) run in an Applied Biosystems 7500 Real-Time PCR instrument with the following settings: reporter SYBR Green, no quencher, passive reference dye ROX, standard ramp speed, and continuous melt curve ramp increment, as described (Allue-Guardia et al. 2022). Relative expression was calculated using the 2^−ΔΔCT^ method (Livak and Schmittgen 2001) with *sigA* as the housekeeping gene.

### *M.tb* infection of primary human monocyte-derived macrophages (MDMs)

Human MDMs were obtained from peripheral blood mononuclear cells (PBMCs) from healthy donors (n=3) after a 5-day differentiation period, as described (Schlesinger 1993; Arcos et al. 2011; Arcos et al. 2017; Moliva et al. 2019; Scordo et al. 2019). At day 6, MDM monolayers were generated (2.5×10^5^/well in a 24-well plate) and infected with ALF-exposed *M.tb* or unexposed bacteria in triplicate for 2 h using a multiplicity of infection (MOI) of 1:1, washed twice with warm RPMI medium, and further incubated in RPMI + 2% autologous serum for up to 24 h at 37°C with 5% CO_2_. At 2 h and 24 h post-infection (hpi), monolayers were lysed and colony-forming units (CFUs) were enumerated (Arcos et al. 2011; Arcos et al. 2017; Moliva et al. 2019; Scordo et al. 2019).

### Statistical analyses

Statistical significance for bacterial uptake and intracellular growth rate was determined by multiple paired Student’s *t* tests (ALF-exposed *vs.* unexposed *M.tb*). A 2-way ANOVA for multiple comparisons with an uncorrected Fisher’s LSD test was used for qPCR data.

## DATA AVAILABILITY

All raw and processed sequencing data generated in this study have been submitted to the NCBI Gene Expression Omnibus (GEO; https://www.ncbi.nlm.nih.gov/geo/) (Edgar et al. 2002) under accession number GSE228998.

## ACKNOWLEDGEMENTS

Strains CDC1551 (mc^2^ 7992 CDC1551 GFP) and W-7642 were a gift from Dr. William R. Jacobs Jr. and Drs. Barry N. Kreiswirth and Barun Mathema, respectively. The following reagent was obtained through BEI Resources, NIAID, NIH:*Mycobacterium tuberculosis*, Strain HN878, NR-13647. We thank Jeremy Glenn and Clinton Christensen at the Texas Biomedical Research Institute Molecular Core for technical support.

## AUTHOR CONTRIBUTIONS

AAG and JBT conceived and designed the study. AAG, AGV, AMS, and AH performed experiments. AAG and AGV analyzed the data. JP, DJM, and MDW performed the bronchoalveolar lavage (BAL). AMOF and AGV processed the BALs and obtained ALFs used in this study. AAG and JBT wrote the first draft and final version of the manuscript. YW provided critical analysis of the data and editing of the manuscript. All authors contributed to manuscript revisions and read and approved the submitted version.

## REFERENCES

Allue-Guardia A, Garcia JI, Torrelles JB. 2021. Evolution of Drug-Resistant Mycobacterium tuberculosis Strains and Their Adaptation to the Human Lung Environment. Front Microbiol 12: 612675.

Allue-Guardia A, Garcia-Vilanova A, Olmo-Fontanez AM, Peters J, Maselli DJ, Wang Y, Turner J, Schlesinger LS, Torrelles JB. 2022. Host- and Age-Dependent Transcriptional Changes in Mycobacterium tuberculosis Cell Envelope Biosynthesis Genes after Exposure to Human Alveolar Lining Fluid. Int J Mol Sci 23.

Arcos J, Diangelo LE, Scordo JM, Sasindran SJ, Moliva JI, Turner J, Torrelles JB. 2015. Lung Mucosa Lining Fluid Modification of Mycobacterium tuberculosis to Reprogram Human Neutrophil Killing Mechanisms. J Infect Dis 212: 948–958.

Arcos J, Sasindran SJ, Fujiwara N, Turner J, Schlesinger LS, Torrelles JB. 2011. Human lung hydrolases delineate Mycobacterium tuberculosis-macrophage interactions and the capacity to control infection. J Immunol 187: 372–381.

Arcos J, Sasindran SJ, Moliva JI, Scordo JM, Sidiki S, Guo H, Venigalla P, Kelley HV, Lin G, Diangelo L et al. 2017. Mycobacterium tuberculosis cell wall released fragments by the action of the human lung mucosa modulate macrophages to control infection in an IL-10-dependent manner. Mucosal Immunol 10: 1248–1258.

Ates LS. 2020. New insights into the mycobacterial PE and PPE proteins provide a framework for future research. Mol Microbiol 113: 4–21.

Banerjee SK, Kumar M, Alokam R, Sharma AK, Chatterjee A, Kumar R, Sahu SK, Jana K, Singh R, Yogeeswari P et al. 2016. Targeting multiple response regulators of Mycobacterium tuberculosis augments the host immune response to infection. Sci Rep 6: 25851.

Bassenden AV, Dumalo L, Park J, Blanchet J, Maiti K, Arya DP, Berghuis AM. 2021. Structural and phylogenetic analyses of resistance to next-generation aminoglycosides conferred by AAC(2’) enzymes. Sci Rep 11: 11614.

Becker K, Sander P. 2016. Mycobacterium tuberculosis lipoproteins in virulence and immunity - fighting with a double-edged sword. FEBS Lett 590: 3800–3819.

Bifani PJ, Plikaytis BB, Kapur V, Stockbauer K, Pan X, Lutfey ML, Moghazeh SL, Eisner W, Daniel TM, Kaplan MH et al. 1996. Origin and interstate spread of a New York City multidrug-resistant Mycobacterium tuberculosis clone family. JAMA 275: 452–457.

Birhanu AG, Gomez-Munoz M, Kalayou S, Riaz T, Lutter T, Yimer SA, Abebe M, Tonjum T. 2022. Proteome Profiling of Mycobacterium tuberculosis Cells Exposed to Nitrosative Stress. ACS Omega 7: 3470–3482.

Birhanu AG, Yimer SA, Kalayou S, Riaz T, Zegeye ED, Holm-Hansen C, Norheim G, Aseffa A, Abebe M, Tonjum T. 2019. Ample glycosylation in membrane and cell envelope proteins may explain the phenotypic diversity and virulence in the Mycobacterium tuberculosis complex. Sci Rep-Uk 9.

Boshoff HI, Xu X, Tahlan K, Dowd CS, Pethe K, Camacho LR, Park TH, Yun CS, Schnappinger D, Ehrt S et al. 2008. Biosynthesis and recycling of nicotinamide cofactors in mycobacterium tuberculosis. An essential role for NAD in nonreplicating bacilli. J Biol Chem 283: 19329–19341.

Brodin P, Poquet Y, Levillain F, Peguillet I, Larrouy-Maumus G, Gilleron M, Ewann F, Christophe T, Fenistein D, Jang J et al. 2010. High content phenotypic cell-based visual screen identifies Mycobacterium tuberculosis acyltrehalose-containing glycolipids involved in phagosome remodeling. PLoS Pathog 6: e1001100.

Brzostek A, Pawelczyk J, Rumijowska-Galewicz A, Dziadek B, Dziadek J. 2009. Mycobacterium tuberculosis is able to accumulate and utilize cholesterol. J Bacteriol 191: 6584–6591.

Bukka A, Price CT, Kernodle DS, Graham JE. 2011. Mycobacterium tuberculosis RNA Expression Patterns in Sputum Bacteria Indicate Secreted Esx Factors Contributing to Growth are Highly Expressed in Active Disease. Front Microbiol 2: 266.

Camus JC, Pryor MJ, Medigue C, Cole ST. 2002. Re-annotation of the genome sequence of Mycobacterium tuberculosis H37Rv. Microbiology (Reading) 148: 2967–2973.

Casonato S, Provvedi R, Dainese E, Palu G, Manganelli R. 2014. Mycobacterium tuberculosis requires the ECF sigma factor SigE to arrest phagosome maturation. Plos One 9: e108893.

Chauhan R, Ravi J, Datta P, Chen T, Schnappinger D, Bassler KE, Balazsi G, Gennaro ML. 2016. Reconstruction and topological characterization of the sigma factor regulatory network of Mycobacterium tuberculosis. Nat Commun 7: 11062.

Chen Y, Lun AT, Smyth GK. 2016. From reads to genes to pathways: differential expression analysis of RNA-Seq experiments using Rsubread and the edgeR quasi-likelihood pipeline. F1000Res 5: 1438.

Cilloni L, Fu H, Vesga JF, Dowdy D, Pretorius C, Ahmedov S, Nair SA, Mosneaga A, Masini E, Sahu S et al. 2020. The potential impact of the COVID-19 pandemic on the tuberculosis epidemic a modelling analysis. EClinicalMedicine 28: 100603.

Cole ST, Brosch R, Parkhill J, Garnier T, Churcher C, Harris D, Gordon SV, Eiglmeier K, Gas S, Barry CE, 3rd et al. 1998. Deciphering the biology of Mycobacterium tuberculosis from the complete genome sequence. Nature 393: 537–544.

Conrad WH, Osman MM, Shanahan JK, Chu F, Takaki KK, Cameron J, Hopkinson-Woolley D, Brosch R, Ramakrishnan L. 2017. Mycobacterial ESX-1 secretion system mediates host cell lysis through bacterium contact-dependent gross membrane disruptions. Proc Natl Acad Sci U S A 114: 1371–1376.

Constant P, Perez E, Malaga W, Laneelle MA, Saurel O, Daffe M, Guilhot C. 2002. Role of the pks15/1 gene in the biosynthesis of phenolglycolipids in the Mycobacterium tuberculosis complex. Evidence that all strains synthesize glycosylated p-hydroxybenzoic methyl esters and that strains devoid of phenolglycolipids harbor a frameshift mutation in the pks15/1 gene. J Biol Chem 277: 38148–38158.

Crowe AM, Casabon I, Brown KL, Liu J, Lian J, Rogalski JC, Hurst TE, Snieckus V, Foster LJ, Eltis LD. 2017. Catabolism of the Last Two Steroid Rings in Mycobacterium tuberculosis and Other Bacteria. mBio 8.

Dang G, Cui Y, Wang L, Li T, Cui Z, Song N, Chen L, Pang H, Liu S. 2018. Extracellular Sphingomyelinase Rv0888 of Mycobacterium tuberculosis Contributes to Pathological Lung Injury of Mycobacterium smegmatis in Mice via Inducing Formation of Neutrophil Extracellular Traps. Front Immunol 9: 677.

Davis MP, van Dongen S, Abreu-Goodger C, Bartonicek N, Enright AJ. 2013. Kraken: a set of tools for quality control and analysis of high-throughput sequence data. Methods 63: 41–49.

Dover LG, Alahari A, Gratraud P, Gomes JM, Bhowruth V, Reynolds RC, Besra GS, Kremer L. 2007. EthA, a common activator of thiocarbamide-containing drugs acting on different mycobacterial targets. Antimicrob Agents Chemother 51: 1055–1063.

Dow A, Sule P, O’Donnell TJ, Burger A, Mattila JT, Antonio B, Vergara K, Marcantonio E, Adams LG, James N et al. 2021. Zinc limitation triggers anticipatory adaptations in Mycobacterium tuberculosis. PLoS Pathog 17: e1009570.

Edgar R, Domrachev M, Lash AE. 2002. Gene Expression Omnibus: NCBI gene expression and hybridization array data repository. Nucleic Acids Res 30: 207–210.

Fontan PA, Aris V, Alvarez ME, Ghanny S, Cheng J, Soteropoulos P, Trevani A, Pine R, Smith I. 2008. Mycobacterium tuberculosis sigma factor E regulon modulates the host inflammatory response. J Infect Dis 198: 877–885.

Gamngoen R, Putim C, Salee P, Phunpae P, Butr-Indr B. 2018. A comparison of Rv0559c and Rv0560c expression in drug-resistant Mycobacterium tuberculosis in response to first-line antituberculosis drugs. Tuberculosis (Edinb) 108: 64–69.

Garcia-Morales L, Leon-Solis L, Monroy-Munoz IE, Talavera-Paulin M, Serafin-Lopez J, Estrada-Garcia I, Rivera-Gutierrez S, Cerna-Cortes JF, Helguera-Repetto AC, Gonzalez YMJA. 2017. Comparative proteomic profiles reveal characteristic Mycobacterium tuberculosis proteins induced by cholesterol during dormancy conditions. Microbiology (Reading) 163: 1237–1247.

Garcia-Vilanova A, Chan J, Torrelles JB. 2019. Underestimated Manipulative Roles of Mycobacterium tuberculosis Cell Envelope Glycolipids During Infection. Front Immunol 10: 2909.

Ge SX, Jung D, Yao R. 2020. ShinyGO: a graphical gene-set enrichment tool for animals and plants. Bioinformatics 36: 2628–2629.

Ge SX, Son EW, Yao R. 2018. iDEP: an integrated web application for differential expression and pathway analysis of RNA-Seq data. BMC Bioinformatics 19: 534.

Gilbert S, Hood L, Seah SYK. 2018. Characterization of an Aldolase Involved in Cholesterol Side Chain Degradation in Mycobacterium tuberculosis. J Bacteriol 200.

Gilmore SA, Schelle MW, Holsclaw CM, Leigh CD, Jain M, Cox JS, Leary JA, Bertozzi CR. 2012. Sulfolipid-1 biosynthesis restricts Mycobacterium tuberculosis growth in human macrophages. ACS Chem Biol 7: 863–870.

Griffin JE, Pandey AK, Gilmore SA, Mizrahi V, McKinney JD, Bertozzi CR, Sassetti CM. 2012. Cholesterol catabolism by Mycobacterium tuberculosis requires transcriptional and metabolic adaptations. Chem Biol 19: 218–227.

Groschel MI, Sayes F, Simeone R, Majlessi L, Brosch R. 2016. ESX secretion systems: mycobacterial evolution to counter host immunity. Nat Rev Microbiol 14: 677–691.

Hampshire T, Soneji S, Bacon J, James BW, Hinds J, Laing K, Stabler RA, Marsh PD, Butcher PD. 2004. Stationary phase gene expression of Mycobacterium tuberculosis following a progressive nutrient depletion: a model for persistent organisms? Tuberculosis (Edinb) 84: 228–238.

Hatzios SK, Baer CE, Rustad TR, Siegrist MS, Pang JM, Ortega C, Alber T, Grundner C, Sherman DR, Bertozzi CR. 2013. Osmosensory signaling in Mycobacterium tuberculosis mediated by a eukaryotic-like Ser/Thr protein kinase. Proc Natl Acad Sci U S A 110: E5069–5077.

Hogan AB, Jewell BL, Sherrard-Smith E, Vesga JF, Watson OJ, Whittaker C, Hamlet A, Smith JA, Winskill P, Verity R et al. 2020. Potential impact of the COVID-19 pandemic on HIV, tuberculosis, and malaria in low-income and middle-income countries: a modelling study. Lancet Glob Health 8: e1132–e1141.

Howard NC, Marin ND, Ahmed M, Rosa BA, Martin J, Bambouskova M, Sergushichev A, Loginicheva E, Kurepina N, Rangel-Moreno J et al. 2018. Mycobacterium tuberculosis carrying a rifampicin drug resistance mutation reprograms macrophage metabolism through cell wall lipid changes (vol 3, pg 1099, 2018). Nat Microbiol 3: 1327–1327.

Hu Y, van der Geize R, Besra GS, Gurcha SS, Liu A, Rohde M, Singh M, Coates A. 2010. 3-Ketosteroid 9alpha-hydroxylase is an essential factor in the pathogenesis of Mycobacterium tuberculosis. Mol Microbiol 75: 107–121.

Huet G, Constant P, Malaga W, Laneelle MA, Kremer K, van Soolingen D, Daffe M, Guilhot C. 2009. A lipid profile typifies the Beijing strains of Mycobacterium tuberculosis: identification of a mutation responsible for a modification of the structures of phthiocerol dimycocerosates and phenolic glycolipids. J Biol Chem 284: 27101–27113.

Islam MS, Richards JP, Ojha AK. 2012. Targeting drug tolerance in mycobacteria: a perspective from mycobacterial biofilms. Expert Rev Anti Infect Ther 10: 1055–1066.

Kapopoulou A, Lew JM, Cole ST. 2011. The MycoBrowser portal: A comprehensive and manually annotated resource for mycobacterial genomes. Tuberculosis 91: 8–13.

Kendall SL, Burgess P, Balhana R, Withers M, Ten Bokum A, Lott JS, Gao C, Uhia-Castro I, Stoker NG. 2010. Cholesterol utilization in mycobacteria is controlled by two TetR-type transcriptional regulators: kstR and kstR2. Microbiology (Reading) 156: 1362–1371.

Krishnamoorthy G, Kaiser P, Constant P, Abu Abed U, Schmid M, Frese CK, Brinkmann V, Daffe M, Kaufmann SHE. 2021. Role of Premycofactocin Synthase in Growth, Microaerophilic Adaptation, and Metabolism of Mycobacterium tuberculosis. mBio 12: e0166521.

Krithika R, Marathe U, Saxena P, Ansari MZ, Mohanty D, Gokhale RS. 2006. A genetic locus required for iron acquisition in Mycobacterium tuberculosis. Proc Natl Acad Sci U S A 103: 2069–2074.

Levillain F, Poquet Y, Mallet L, Mazeres S, Marceau M, Brosch R, Bange FC, Supply P, Magalon A, Neyrolles O. 2017. Horizontal acquisition of a hypoxia-responsive molybdenum cofactor biosynthesis pathway contributed to Mycobacterium tuberculosis pathoadaptation. PLoS Pathog 13: e1006752.

Lew JM, Kapopoulou A, Jones LM, Cole ST. 2011. TubercuList--10 years after. Tuberculosis (Edinb) 91: 1–7.

Li QF, T.; Li, C.; Fan, X.; Xie, J. 2016. Mycobacterial IclR family transcriptional factor *Rv2989* is specifically involved in isoniazid tolerance by regulating the expression of catalase encoding gene *katG*. RCS Advances 6: 54661–54667.

Livak KJ, Schmittgen TD. 2001. Analysis of relative gene expression data using real-time quantitative PCR and the 2(T)(-Delta Delta C) method. Methods 25: 402–408.

Love MI, Huber W, Anders S. 2014. Moderated estimation of fold change and dispersion for RNA-seq data with DESeq2. Genome Biol 15: 550.

Maitra A, Munshi T, Healy J, Martin LT, Vollmer W, Keep NH, Bhakta S. 2019. Cell wall peptidoglycan in Mycobacterium tuberculosis: An Achilles’ heel for the TB-causing pathogen. FEMS Microbiol Rev 43: 548–575.

McCarthy DJ, Chen Y, Smyth GK. 2012. Differential expression analysis of multifactor RNA-Seq experiments with respect to biological variation. Nucleic Acids Res 40: 4288–4297.

Melly G, Purdy GE. 2019. MmpL Proteins in Physiology and Pathogenesis of M. tuberculosis. Microorganisms 7.

Metsalu T, Vilo J. 2015. ClustVis: a web tool for visualizing clustering of multivariate data using Principal Component Analysis and heatmap. Nucleic Acids Res 43: W566–570.

Mohn WW, van der Geize R, Stewart GR, Okamoto S, Liu J, Dijkhuizen L, Eltis LD. 2008. The actinobacterial mce4 locus encodes a steroid transporter. J Biol Chem 283: 35368–35374.

Moliva JI, Duncan MA, Olmo-Fontanez A, Akhter A, Arnett E, Scordo JM, Ault R, Sasindran SJ, Azad AK, Montoya MJ et al. 2019. The Lung Mucosa Environment in the Elderly Increases Host Susceptibility to Mycobacterium tuberculosis Infection. J Infect Dis 220: 514–523.

Moliva JI, Rajaram MV, Sidiki S, Sasindran SJ, Guirado E, Pan XJ, Wang SH, Ross P, Jr., Lafuse WP, Schlesinger LS et al. 2014. Molecular composition of the alveolar lining fluid in the aging lung. Age (Dordr) 36: 9633.

Oliveros JC. 2007. VENNY. An interactive tool for comparing lists with Venn Diagrams.

Olmo-Fontánez AM, Scordo JM, Garcia-Vilanova A, Maselli DJ, Peters JI, Restrepo BI, Clemens DL, Turner J, Schlesinger LS, Torrelles JB. 2021. Human alveolar lining fluid from the elderly promotes Mycobacterium tuberculosis growth in alveolar epithelial cells and bacterial translocation into the cytosol. bioRxiv doi:10.1101/2021.05.12.443884:2021.2005.2012.443884.

Pal R, Hameed S, Kumar P, Singh S, Fatima Z. 2017. Comparative lipidomics of drug sensitive and resistant Mycobacterium tuberculosis reveals altered lipid imprints. *3* Biotech 7.

Pandey AK, Sassetti CM. 2008. Mycobacterial persistence requires the utilization of host cholesterol. Proc Natl Acad Sci U S A 105: 4376–4380.

Pang JM, Layre E, Sweet L, Sherrid A, Moody DB, Ojha A, Sherman DR. 2012. The polyketide Pks1 contributes to biofilm formation in Mycobacterium tuberculosis. J Bacteriol 194: 715–721.

Pawelczyk J, Brzostek A, Minias A, Plocinski P, Rumijowska-Galewicz A, Strapagiel D, Zakrzewska-Czerwinska J, Dziadek J. 2021. Cholesterol-dependent transcriptome remodeling reveals new insight into the contribution of cholesterol to Mycobacterium tuberculosis pathogenesis. Sci Rep 11: 12396.

Perrone F, De Siena B, Muscariello L, Kendall SL, Waddell SJ, Sacco M. 2017. A Novel TetR-Like Transcriptional Regulator Is Induced in Acid-Nitrosative Stress and Controls Expression of an Efflux Pump in Mycobacteria. Front Microbiol 8: 2039.

Pethe K, Swenson DL, Alonso S, Anderson J, Wang C, Russell DG. 2004. Isolation of Mycobacterium tuberculosis mutants defective in the arrest of phagosome maturation. Proc Natl Acad Sci U S A 101: 13642–13647.

Peyron P, Vaubourgeix J, Poquet Y, Levillain F, Botanch C, Bardou F, Daffe M, Emile JF, Marchou B, Cardona PJ et al. 2008. Foamy macrophages from tuberculous patients’ granulomas constitute a nutrient-rich reservoir for M. tuberculosis persistence. PLoS Pathog 4: e1000204.

Pisu D, Provvedi R, Espinosa DM, Payan JB, Boldrin F, Palu G, Hernandez-Pando R, Manganelli R. 2017. The Alternative Sigma Factors SigE and SigB Are Involved in Tolerance and Persistence to Antitubercular Drugs. Antimicrob Agents Chemother 61.

Quadri LE, Sello J, Keating TA, Weinreb PH, Walsh CT. 1998. Identification of a Mycobacterium tuberculosis gene cluster encoding the biosynthetic enzymes for assembly of the virulence-conferring siderophore mycobactin. Chem Biol 5: 631–645.

Ramos B, Gordon SV, Cunha MV. 2020. Revisiting the expression signature of pks15/1 unveils regulatory patterns controlling phenolphtiocerol and phenolglycolipid production in pathogenic mycobacteria. Plos One 15: e0229700.

Rebollo-Ramirez S, Larrouy-Maumus G. 2019. NaCl triggers the CRP-dependent increase of cAMP in Mycobacterium tuberculosis. Tuberculosis (Edinb) 116: 8–16.

Reed MB, Domenech P, Manca C, Su H, Barczak AK, Kreiswirth BN, Kaplan G, Barry CE. 2004. A glycolipid of hypervirulent tuberculosis strains that inhibits the innate immune response. Nature 431: 84–87.

Reed MB, Gagneux S, DeRiemer K, Small PM, Barry CE. 2007. The W-Beijing lineage of Mycobacterium tuberculosis overproduces triglycerides and has the DosR dormancy regulon constitutively upregulated. J Bacteriol 189: 2583–2589.

Robinson MD, McCarthy DJ, Smyth GK. 2010. edgeR: a Bioconductor package for differential expression analysis of digital gene expression data. Bioinformatics 26: 139–140.

Sachdeva P, Misra R, Tyagi AK, Singh Y. 2010. The sigma factors of Mycobacterium tuberculosis: regulation of the regulators. FEBS J 277: 605–626.

Salina EG, Grigorov AS, Bychenko OS, Skvortsova YV, Mamedov IZ, Azhikina TL, Kaprelyants AS. 2019. Resuscitation of Dormant “Non-culturable” Mycobacterium tuberculosis Is Characterized by Immediate Transcriptional Burst. Front Cell Infect Microbiol 9: 272.

Santangelo MP, Klepp L, Nunez-Garcia J, Blanco FC, Soria M, Garcia-Pelayo MDC, Bianco MV, Cataldi AA, Golby P, Jackson M et al. 2009. Mce3R, a TetR-type transcriptional repressor, controls the expression of a regulon involved in lipid metabolism in Mycobacterium tuberculosis. Microbiology (Reading) 155: 2245–2255.

Saraav I, Singh S, Pandey K, Sharma M, Sharma S. 2017. Mycobacterium tuberculosis MymA is a TLR2 agonist that activate macrophages and a TH1 response. Tuberculosis (Edinb) 106: 16–24.

Schlesinger LS. 1993. Macrophage phagocytosis of virulent but not attenuated strains of Mycobacterium tuberculosis is mediated by mannose receptors in addition to complement receptors. J Immunol 150: 2920–2930.

Scordo JM, Arcos J, Kelley HV, Diangelo L, Sasindran SJ, Youngmin E, Wewers MD, Wang SH, Balada-Llasat JM, Torrelles JB. 2017. Mycobacterium tuberculosis Cell Wall Fragments Released upon Bacterial Contact with the Human Lung Mucosa Alter the Neutrophil Response to Infection. Front Immunol 8: 307.

Scordo JM, Olmo-Fontanez AM, Kelley HV, Sidiki S, Arcos J, Akhter A, Wewers MD, Torrelles JB. 2019. The human lung mucosa drives differential Mycobacterium tuberculosis infection outcome in the alveolar epithelium. Mucosal Immunol 12: 795–804.

Sethi D, Mahajan S, Singh C, Lama A, Hade MD, Gupta P, Dikshit KL. 2016. Lipoprotein LprI of Mycobacterium tuberculosis Acts as a Lysozyme Inhibitor. J Biol Chem 291: 2938–2953.

Shah S, Cannon JR, Fenselau C, Briken V. 2015. A Duplicated ESAT-6 Region of ESX-5 Is Involved in Protein Export and Virulence of Mycobacteria. Infect Immun 83: 4349–4361.

Singh A, Gupta R, Vishwakarma RA, Narayanan PR, Paramasivan CN, Ramanathan VD, Tyagi AK. 2005. Requirement of the mymA operon for appropriate cell wall ultrastructure and persistence of Mycobacterium tuberculosis in the spleens of guinea pigs. J Bacteriol 187: 4173–4186.

Singh A, Jain S, Gupta S, Das T, Tyagi AK. 2003. mymA operon of Mycobacterium tuberculosis: its regulation and importance in the cell envelope. FEMS Microbiol Lett 227: 53–63.

Skovierova H, Larrouy-Maumus G, Zhang J, Kaur D, Barilone N, Kordulakova J, Gilleron M, Guadagnini S, Belanova M, Prevost MC et al. 2009. AftD, a novel essential arabinofuranosyltransferase from mycobacteria. Glycobiology 19: 1235–1247.

Slayden RA, Barry CE, 3rd. 2002. The role of KasA and KasB in the biosynthesis of meromycolic acids and isoniazid resistance in Mycobacterium tuberculosis. Tuberculosis (Edinb) 82: 149–160.

Steenken W, Oatway WH, Petroff SA. 1934. Biological Studies of the Tubercle Bacillus: Iii. Dissociation and Pathogenicity of the R and S Variants of the Human Tubercle Bacillus (H(37)). J Exp Med 60: 515–540.

Subbian S, Bandyopadhyay N, Tsenova L, O’Brien P, Khetani V, Kushner NL, Peixoto B, Soteropoulos P, Bader JS, Karakousis PC et al. 2013. Early innate immunity determines outcome of Mycobacterium tuberculosis pulmonary infection in rabbits. Cell Commun Signal 11: 60.

Subbian S, Tsenova L, Yang G, O’Brien P, Parsons S, Peixoto B, Taylor L, Fallows D, Kaplan G. 2011. Chronic pulmonary cavitary tuberculosis in rabbits: a failed host immune response. Open Biol 1: 110016.

Tabone O, Verma R, Singhania A, Chakravarty P, Branchett WJ, Graham CM, Lee J, Trang T, Reynier F, Leissner P et al. 2021. Blood transcriptomics reveal the evolution and resolution of the immune response in tuberculosis. J Exp Med 218.

Tang J, Liu Z, Shi Y, Zhan L, Qin C. 2020. Whole Genome and Transcriptome Sequencing of Two Multi-Drug Resistant Mycobacterium tuberculosis Strains to Facilitate Illustrating Their Virulence in vivo. Front Cell Infect Microbiol 10: 219.

Torrelles JB, Schlesinger LS. 2017. Integrating Lung Physiology, Immunology, and Tuberculosis. Trends Microbiol 25: 688–697.

Tripathi D, Chandra H, Bhatnagar R. 2013. Poly-L-glutamate/glutamine synthesis in the cell wall of Mycobacterium bovis is regulated in response to nitrogen availability. BMC Microbiol 13: 226.

Tufariello JM, Chapman JR, Kerantzas CA, Wong KW, Vilcheze C, Jones CM, Cole LE, Tinaztepe E, Thompson V, Fenyo D et al. 2016. Separable roles for Mycobacterium tuberculosis ESX-3 effectors in iron acquisition and virulence. Proc Natl Acad Sci U S A 113: E348–357.

Valway SE, Sanchez MP, Shinnick TF, Orme I, Agerton T, Hoy D, Jones JS, Westmoreland H, Onorato IM. 1998. An outbreak involving extensive transmission of a virulent strain of Mycobacterium tuberculosis. N Engl J Med 338: 633–639.

Vargas R, Freschi L, Marin M, Epperson LE, Smith M, Oussenko I, Durbin D, Strong M, Salfinger M, Farhat MR. 2021. In-host population dynamics of Mycobacterium tuberculosis complex during active disease. Elife 10.

Velayati AA, Farnia P, Ibrahim TA, Haroun RZ, Kuan HO, Ghanavi J, Farnia P, Kabarei AN, Tabarsi P, Omar AR et al. 2009. Differences in Cell Wall Thickness between Resistant and Nonresistant Strains of Mycobacterium tuberculosis: Using Transmission Electron Microscopy. Chemotherapy 55: 303–307.

Wang C, Mao Y, Yu J, Zhu L, Li M, Wang D, Dong D, Liu J, Gao Q. 2013. PhoY2 of mycobacteria is required for metabolic homeostasis and stress response. J Bacteriol 195: 243–252.

WHO. 2022. Global Tuberculosis Report 2022.

Wingfield T, Karmadwala F, MacPherson P, Millington KA, Walker NF, Cuevas LE, Squire SB. 2021. Challenges and opportunities to end tuberculosis in the COVID-19 era. Lancet Respir Med 9: 556–558.

Wipperman MF, Sampson NS, Thomas ST. 2014. Pathogen roid rage: cholesterol utilization by Mycobacterium tuberculosis. Crit Rev Biochem Mol Biol 49: 269–293.

Wong KW. 2017. The Role of ESX-1 in Mycobacterium tuberculosis Pathogenesis. Microbiol Spectr 5.

Yang X, Yuan T, Ma R, Chacko KI, Smith M, Deikus G, Sebra R, Kasarskis A, van Bakel H, Franzblau SG et al. 2019. Mce3R Stress-Resistance Pathway Is Vulnerable to Small-Molecule Targeting That Improves Tuberculosis Drug Activities. ACS Infect Dis 5: 1239–1251.

Yu X, Gao X, Zhu K, Yin H, Mao X, Wojdyla JA, Qin B, Huang H, Wang M, Sun YC et al. 2020. Characterization of a toxin-antitoxin system in Mycobacterium tuberculosis suggests neutralization by phosphorylation as the antitoxicity mechanism. Commun Biol 3: 216.

Zhang L, English D, Andersen BR. 1991. Activation of human neutrophils by Mycobacterium tuberculosis-derived sulfolipid-1. J Immunol 146: 2730–2736.

Zhang L, Hendrickson RC, Meikle V, Lefkowitz EJ, Ioerger TR, Niederweis M. 2020. Comprehensive analysis of iron utilization by Mycobacterium tuberculosis. PLoS Pathog 16: e1008337.

