## Supplemental Figures for "Exposure of *Mycobacterium tuberculosis* to human alveolar lining fluid shows temporal and strain-specific adaptation to the lung environment"

### Supplemental Figure S1

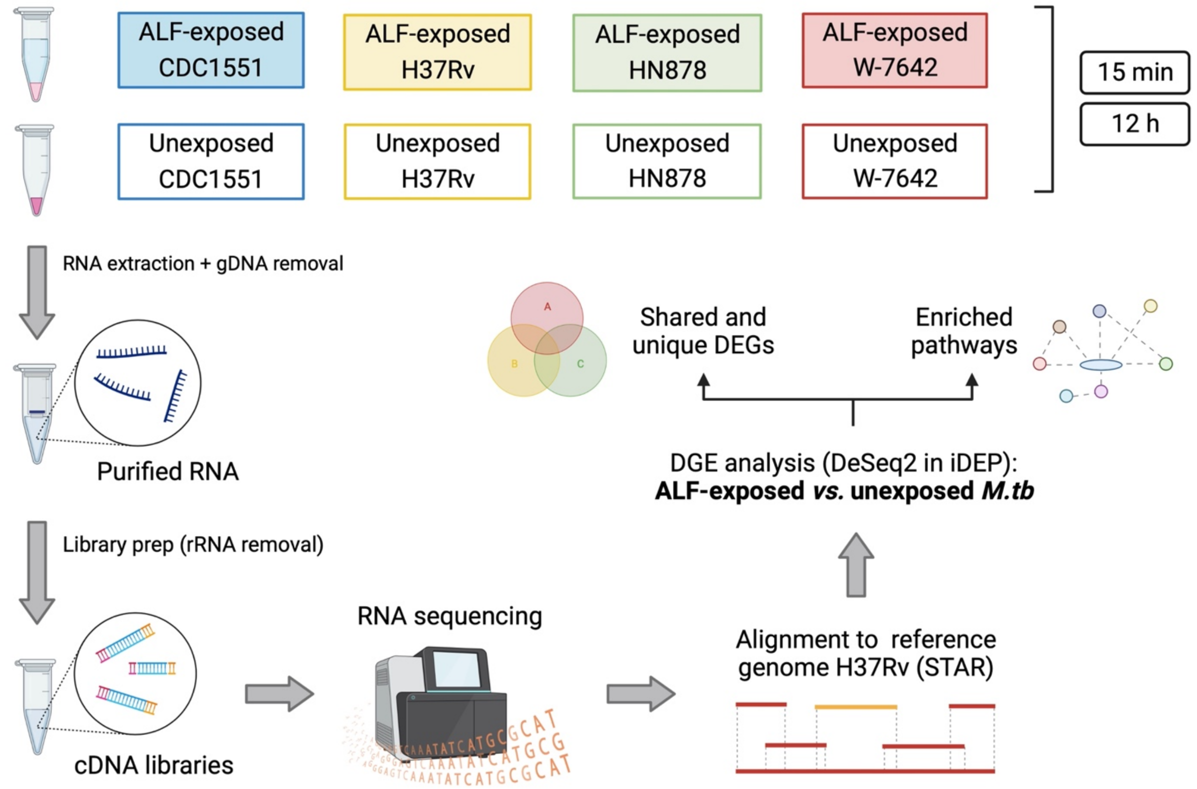


**Supplemental Figure S1**. Diagram of experimental conditions and strategy for RNA-seq and data analyses.

**Supplemental Figure S2**

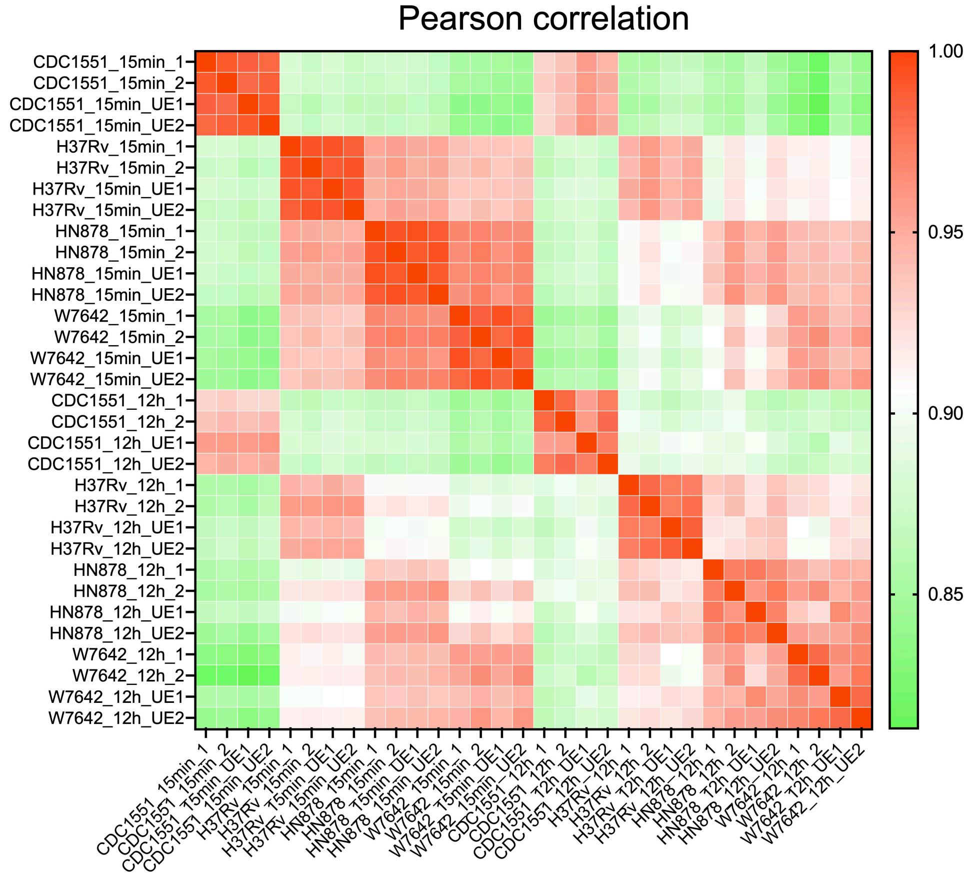


**Supplemental figure S2**. Pearson correlation calculated in iDEP.94 and represented as a heatmap. Biological replicates are shown as 1 and 2; UE (unexposed *M.tb*), otherwise indicates ALF-exposed bacteria.

### Supplemental Figure S3

A

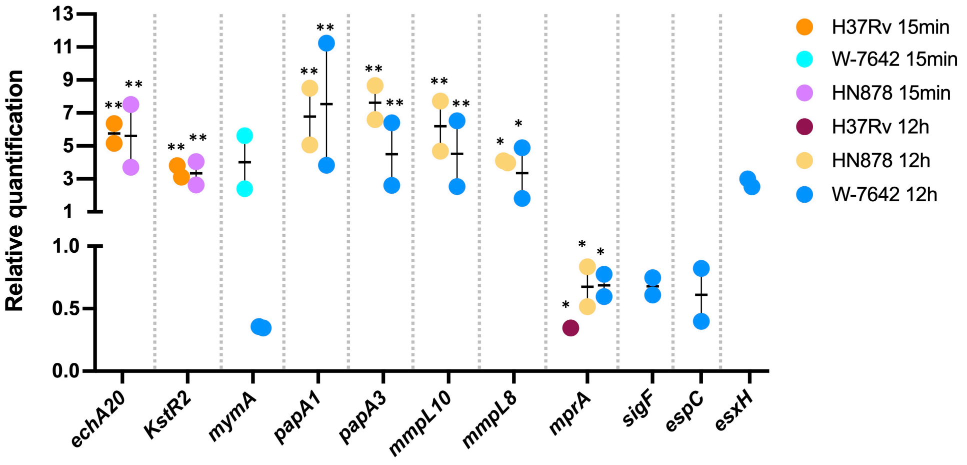


B

| | H37Rv 15min | | HN878 15min | | W-7642 15min | | H37Rv 12h | | HN878 12h | | W-7642 12h | |
|---|---|---|---|---|---|---|---|---|---|---|---|---|
| | qPCR | RNAseq | qPCR | RNAseq | qPCR | RNAseq | qPCR | RNAseq | qPCR | RNAseq | qPCR | RNAseq |
| ***echA20*** | 5.75 | 4.68 | 5.61 | 4.20 | | | | | | | | |
| ***KstR2*** | 3.45 | 3.05 | 3.34 | 3.00 | | | | | | | | |
| ***mymA*** | | | | | 4.01 | 3.52 | | | | | 0.35 | 0.37 |
| ***papA1*** | | | | | | | | | 6.78 | 4.57 | 7.53 | 4.63 |
| ***papA3*** | | | | | | | | | 7.63 | 4.44 | 4.50 | 3.10 |
| ***mmpL10*** | | | | | | | | | 6.20 | 3.66 | 4.53 | 3.06 |
| ***mmpL8*** | | | | | | | | | 4.04 | 3.32 | 3.35 | 3.20 |
| ***mprA*** | | | | | | | 0.35 | 0.36 | 0.68 | 0.46 | 0.68 | 0.29 |
| ***sigF*** | | | | | | | | | | | 0.68 | 0.40 |
| ***espC*** | | | | | | | | | | | 0.61 | 0.39 |
| ***esxH*** | | | | | | | | | | | 2.76 | 2.07 |

**Supplemental figure S3**. Validation of gene expression by RT-qPCR for selected targets and strains. **A**) RT-qPCR was performed as described in the methods and relative quantification was calculated using the $2^{-\Delta\Delta CT}$ method with *sigA* as the reference gene. Values are expressed as fold changes (ALF-exposed *vs.* unexposed *M.tb*) and plotted using Graphpad Prism v9.1.1. Statistical analysis: 2-way ANOVA for multiple comparisons with an uncorrected Fisher's LSD test; * *p*-value < 0.05; ** *p*-value < 0.005. **B**) Comparison of fold changes (n=2 replicates) in qPCR and RNA-seq data for selected genes (ALF-exposed vs. unexposed *M.tb*). $Log_2FC$ in RNA-seq data were converted to FC. Upregulated and downregulated genes are shown in yellow and blue, respectively.
