## Supplemental Table S1 for "Exposure of *Mycobacterium tuberculosis* to human alveolar lining fluid shows temporal and strain-specific adaptation to the lung environment"

**Supplemental Table S1**. Post-alignment stats, including number and % of aligned reads, read quality, genome coverage and sequencing depth, and %GC content.

| Strain | Time | ALF exposure | Replicate | Total reads | Aligned (%) | Total aligned (reads) | Coverage | Coverage depth | Quality | %GC |
|---|---|---|---|---|---|---|---|---|---|---|
| CDC1551 | 15min | Exposed | 1 | 5509415 | 82.4 | 4537310 | 98.4 | 166.3 | 33.6 | 62.0 |
| CDC1551 | 15min | Exposed | 2 | 5934111 | 93.1 | 5525863 | 99.1 | 202.3 | 33.7 | 62.2 |
| CDC1551 | 15min | Unexposed | 1 | 6455659 | 73.3 | 4734560 | 98.0 | 176.9 | 33.8 | 62.0 |
| CDC1551 | 15min | Unexposed | 2 | 6962857 | 93.4 | 6503758 | 98.5 | 242.6 | 33.7 | 62.3 |
| H37Rv | 15min | Exposed | 1 | 6846858 | 97.9 | 6699673 | 99.3 | 233.2 | 33.7 | 63.3 |
| H37Rv | 15min | Exposed | 2 | 6300460 | 98.1 | 6179577 | 99.2 | 214.7 | 33.8 | 63.3 |
| H37Rv | 15min | Unexposed | 1 | 6058552 | 98.1 | 5943270 | 99.6 | 206.4 | 33.8 | 63.3 |
| H37Rv | 15min | Unexposed | 2 | 5932530 | 97.6 | 5789050 | 99.2 | 200.8 | 33.4 | 63.1 |
| HN878 | 15min | Exposed | 1 | 5737808 | 97.4 | 5588822 | 98.4 | 197.7 | 33.8 | 62.9 |
| HN878 | 15min | Exposed | 2 | 11379796 | 82.9 | 9431019 | 99.0 | 329.7 | 34.0 | 62.9 |
| HN878 | 15min | Unexposed | 1 | 5824435 | 97.4 | 5675688 | 98.5 | 200.5 | 33.7 | 62.9 |
| HN878 | 15min | Unexposed | 2 | 5886080 | 69.9 | 4113780 | 97.8 | 145.6 | 33.7 | 62.6 |
| W-7642 | 15min | Exposed | 1 | 6209066 | 92.1 | 5721249 | 98.8 | 200.0 | 33.7 | 63.2 |
| W-7642 | 15min | Exposed | 2 | 5735475 | 93.5 | 5362804 | 98.6 | 187.2 | 33.8 | 63.0 |
| W-7642 | 15min | Unexposed | 1 | 5498289 | 91.6 | 5035154 | 98.7 | 175.9 | 33.4 | 63.1 |
| W-7642 | 15min | Unexposed | 2 | 6002067 | 93.2 | 5596776 | 98.8 | 195.2 | 33.5 | 63.0 |
| CDC1551 | 12h | Exposed | 1 | 5560129 | 85.7 | 4765889 | 99.4 | 170.2 | 33.5 | 60.2 |
| CDC1551 | 12h | Exposed | 2 | 5124697 | 93.0 | 4766215 | 99.4 | 172.0 | 33.6 | 61.3 |
| CDC1551 | 12h | Unexposed | 1 | 5272743 | 67.0 | 3534029 | 98.2 | 126.9 | 33.7 | 61.3 |
| CDC1551 | 12h | Unexposed | 2 | 5361190 | 92.6 | 4964967 | 99.4 | 180.9 | 33.8 | 61.9 |
| H37Rv | 12h | Exposed | 1 | 5435225 | 86.8 | 4717460 | 99.8 | 162.5 | 33.1 | 61.4 |
| H37Rv | 12h | Exposed | 2 | 5862354 | 92.3 | 5410172 | 99.3 | 187.3 | 33.3 | 61.4 |
| H37Rv | 12h | Unexposed | 1 | 5017770 | 94.7 | 4750163 | 99.0 | 164.9 | 33.7 | 61.7 |
| H37Rv | 12h | Unexposed | 2 | 6799187 | 95.1 | 6463956 | 99.6 | 223.1 | 33.8 | 61.8 |
| HN878 | 12h | Exposed | 1 | 6609944 | 94.6 | 6253247 | 99.1 | 218.1 | 33.8 | 61.2 |
| HN878 | 12h | Exposed | 2 | 5406542 | 70.3 | 3799992 | 99.0 | 132.3 | 33.6 | 61.0 |
| HN878 | 12h | Unexposed | 1 | 4106746 | 95.2 | 3910647 | 98.4 | 137.2 | 33.8 | 61.7 |
| HN878 | 12h | Unexposed | 2 | 6914330 | 61.5 | 4255521 | 98.0 | 149.7 | 33.8 | 61.7 |
| W-7642 | 12h | Exposed | 1 | 6369547 | 91.3 | 5817649 | 99.0 | 202.2 | 33.8 | 62.5 |
| W-7642 | 12h | Exposed | 2 | 6428728 | 91.7 | 5898287 | 99.2 | 204.3 | 33.9 | 62.1 |
| W-7642 | 12h | Unexposed | 1 | 6847018 | 90.7 | 6211375 | 99.1 | 215.5 | 33.8 | 62.2 |
| W-7642 | 12h | Unexposed | 2 | 7456592 | 92.7 | 6909877 | 99.2 | 239.2 | 33.8 | 62.4 |
