## Supplemental Table S2 for "Exposure of *Mycobacterium tuberculosis* to human alveolar lining fluid shows temporal and strain-specific adaptation to the lung environment"

**Supplemental Table S2**. Results of DE analysis (ALF-exposed vs. unexposed *M.tb* ) of 4 *M.tb* strains (CDC1551, H37Rv, HN878, and W-7642) at 15 min and 12 h. DE is shown as log2 FC; up- and downregulated genes in yellow and blue, respectively; significant DEGs (log2 FC equal or greater than an absolute value of 1 and FDR <1) are highlighted in bold. DEGs are listed based on functional category (Mycobrowser).

| | | CDC1551 15min | | H37Rv 15min | | HN878 15min | | W-7642 15min | | CDC1551 12h | | H37Rv 12h | | HN878 12h | | W-7642 12h | | | |
|---|---|---|---|---|---|---|---|---|---|---|---|---|---|---|---|---|---|---|---|
| Locus tag | Name | log2FC | padj | log2FC | padj | log2FC | padj | log2FC | padj | log2FC | padj | log2FC | padj | log2FC | padj | log2FC | padj | Product | Functional_Category |
| **Rv1987** | *Rv1987* | -0.170 | 0.858 | 0.077 | 0.992 | 0.076 | 1.000 | 0.135 | 1.000 | -0.416 | 0.441 | **-1.968** | **0.000** | **-1.175** | **0.003** | **-1.784** | **0.000** | Possible chitinase | cell wall and cell processes |
| **Rv1522c** | *mmpL12* | 0.077 | 0.951 | 0.197 | 0.990 | 0.040 | 1.000 | 0.151 | 1.000 | -0.221 | 0.604 | **-1.794** | **0.000** | **-1.437** | **0.000** | **-1.857** | **0.000** | MmpL12 | cell wall and cell processes |
| **Rv3289c** | *Rv3289c* | 0.537 | 0.385 | 0.334 | 0.990 | -0.116 | 1.000 | 0.364 | 1.000 | -0.938 | 0.019 | **-1.595** | **0.000** | -0.718 | 0.125 | **-1.657** | **0.000** | Possible transmembrane protein | cell wall and cell processes |
| **Rv1690** | *lprJ* | -0.811 | 0.346 | -0.914 | 0.712 | -0.368 | 1.000 | -0.083 | 1.000 | -0.632 | 0.382 | **-1.461** | **0.015** | -0.996 | 0.176 | **-1.223** | **0.025** | Probable lipoprotein LprJ | cell wall and cell processes |
| **Rv2115c** | *mpa* | 0.027 | 0.983 | 0.252 | 0.956 | 0.032 | 1.000 | 0.267 | 1.000 | -0.827 | 0.000 | **-1.451** | **0.000** | **-1.138** | **0.000** | **-1.322** | **0.000** | Mycobacterial proteasome ATPase Mpa | cell wall and cell processes |
| **Rv3675** | *Rv3675* | 0.086 | 0.982 | -0.328 | 0.990 | 0.163 | 1.000 | -0.402 | 1.000 | -0.735 | 0.416 | **-1.347** | **0.077** | -1.009 | 0.265 | **-1.340** | **0.042** | Possible membrane protein | cell wall and cell processes |
| **Rv0188** | *Rv0188* | -0.101 | 0.978 | 0.277 | 0.990 | -0.147 | 1.000 | 0.048 | 1.000 | -0.203 | 0.851 | **-1.309** | **0.076** | -1.011 | 0.244 | -0.904 | 0.177 | Probable conserved transmembrane protein | cell wall and cell processes |
| **Rv3810** | *pirG* | -0.082 | 0.968 | 0.013 | 0.994 | 0.170 | 1.000 | -0.220 | 1.000 | 0.356 | 0.555 | **-1.188** | **0.000** | **-1.122** | **0.000** | **-1.392** | **0.000** | Exported repetitive protein precursor PirG (cell surface protein) (EXP53) | cell wall and cell processes |
| **Rv0835** | *lpqQ* | -0.118 | 0.910 | 0.228 | 0.990 | 0.091 | 1.000 | -0.177 | 1.000 | -0.701 | 0.081 | **-1.181** | **0.000** | **-1.415** | **0.000** | **-1.334** | **0.000** | Possible lipoprotein LpqQ | cell wall and cell processes |
| **Rv2709** | *Rv2709* | 0.127 | 0.821 | 0.298 | 0.823 | 0.013 | 1.000 | 0.236 | 1.000 | -0.719 | 0.003 | **-1.109** | **0.000** | -0.360 | 0.272 | -0.565 | 0.010 | Probable conserved transmembrane protein | cell wall and cell processes |
| **Rv2320c** | *rocE* | -0.191 | 0.909 | 0.037 | 0.992 | 0.027 | 1.000 | 0.110 | 1.000 | -0.240 | 0.757 | **-1.095** | **0.054** | -0.935 | 0.156 | **-1.093** | **0.024** | Probable cationic amino acid transport integral membrane protein RocE | cell wall and cell processes |
| **Rv0888** | *Rv0888* | **-1.389** | **0.053** | -0.368 | 0.990 | 0.073 | 1.000 | -0.441 | 1.000 | -0.276 | 0.776 | -1.057 | 0.129 | -0.710 | 0.434 | **-1.439** | **0.014** | Probable exported protein | cell wall and cell processes |
| **Rv0309** | *Rv0309* | -0.137 | 0.894 | -0.315 | 0.976 | -0.078 | 1.000 | -0.026 | 1.000 | -0.097 | 0.865 | **-1.020** | **0.003** | -0.394 | 0.425 | -0.830 | 0.007 | Possible conserved exported protein | cell wall and cell processes |
| **Rv1038c** | *esxJ* | 0.736 | 0.436 | -0.115 | 0.992 | 0.150 | 1.000 | -0.046 | 1.000 | -0.509 | 0.509 | -1.015 | 0.113 | -0.271 | 0.779 | **-1.098** | **0.039** | ESAT-6 like protein EsxJ (ESAT-6 like protein 2) | cell wall and cell processes |
| **Rv0870c** | *Rv0870c* | 0.024 | 0.978 | 0.189 | 0.964 | -0.064 | 1.000 | 0.097 | 1.000 | -0.858 | 0.000 | **-1.004** | **0.000** | -0.526 | 0.012 | -0.726 | 0.000 | Possible conserved integral membrane protein | cell wall and cell processes |
| **Rv0179c** | *lprO* | -0.376 | 0.494 | -0.153 | 0.990 | 0.038 | 1.000 | -0.026 | 1.000 | -0.113 | 0.830 | -0.979 | 0.002 | **-1.015** | **0.003** | **-1.151** | **0.000** | Possible lipoprotein LprO | cell wall and cell processes |
| **Rv2163c** | *pbpB* | 0.004 | 1.000 | 0.015 | 0.994 | 0.048 | 1.000 | 0.047 | 1.000 | 0.132 | 0.775 | -0.959 | 0.000 | -0.778 | 0.007 | **-1.056** | **0.000** | Probable penicillin-binding membrane protein PbpB | cell wall and cell processes |
| **Rv2169c** | *Rv2169c* | 0.252 | 0.842 | 0.225 | 0.990 | 0.098 | 1.000 | 0.305 | 1.000 | -0.862 | 0.087 | -0.854 | 0.094 | -0.313 | 0.673 | **-1.007** | **0.018** | Probable conserved transmembrane protein | cell wall and cell processes |
| **Rv1517** | *Rv1517* | 0.226 | 0.867 | 0.107 | 0.992 | -0.086 | 1.000 | 0.199 | 1.000 | 0.420 | 0.474 | -0.811 | 0.081 | **-1.021** | **0.033** | -0.963 | 0.015 | Conserved hypothetical transmembrane protein | cell wall and cell processes |
| **Rv0522** | *gabP* | 0.281 | 0.429 | 0.238 | 0.964 | -0.029 | 1.000 | 0.190 | 1.000 | -0.225 | 0.456 | -0.757 | 0.001 | -0.340 | 0.249 | **-1.018** | **0.000** | Probable GABA permease GabP (4-amino butyrate transport carrier) (GAMA-aminobutyrate permease) | cell wall and cell processes |
| **Rv3679** | *Rv3679* | -0.671 | 0.140 | -0.323 | 0.985 | -0.020 | 1.000 | -0.355 | 1.000 | **-1.251** | **0.000** | -0.740 | 0.054 | -0.578 | 0.202 | -0.893 | 0.007 | Probable anion transporter ATPase | cell wall and cell processes |
| **Rv0559c** | *Rv0559c* | 0.156 | 0.785 | 0.131 | 0.990 | -0.001 | 1.000 | 0.285 | 1.000 | **-1.182** | **0.000** | -0.737 | 0.002 | -0.353 | 0.271 | -0.650 | 0.003 | Possible conserved secreted protein | cell wall and cell processes |
| **Rv2094c** | *tatA* | 0.091 | 0.863 | 0.152 | 0.990 | -0.012 | 1.000 | 0.161 | 1.000 | -0.703 | 0.000 | -0.714 | 0.000 | -0.635 | 0.007 | **-1.175** | **0.000** | Sec-independent protein translocase membrane-bound protein TatA | cell wall and cell processes |
| **Rv0821c** | *phoY2* | 0.567 | 0.284 | 0.318 | 0.990 | 0.050 | 1.000 | 0.275 | 1.000 | **-1.022** | **0.005** | -0.681 | 0.094 | -0.545 | 0.251 | -0.741 | 0.031 | Probable phosphate-transport system transcriptional regulatory protein PhoY2 | cell wall and cell processes |
| **Rv0655** | *mkl* | 0.103 | 0.863 | 0.158 | 0.990 | 0.075 | 1.000 | 0.185 | 1.000 | **-1.465** | **0.000** | -0.662 | 0.001 | -0.047 | 0.912 | -0.485 | 0.011 | Possible ribonucleotide-transport ATP-binding protein ABC transporter Mkl | cell wall and cell processes |
| **Rv1440** | *secG* | -0.093 | 0.950 | 0.070 | 0.992 | 0.083 | 1.000 | 0.004 | 1.000 | **-1.228** | **0.001** | -0.636 | 0.161 | -0.167 | 0.815 | -0.424 | 0.289 | Probable protein-export membrane protein (translocase subunit) SecG | cell wall and cell processes |
| **Rv2146c** | *Rv2146c* | -0.248 | NA | 0.091 | NA | 0.120 | 1.000 | -0.436 | 1.000 | **-1.392** | **0.020** | -0.611 | 0.248 | **-1.229** | **0.012** | -0.654 | 0.097 | Possible conserved transmembrane protein | cell wall and cell processes |
| **Rv3891c** | *esxD* | -0.805 | 0.218 | -0.394 | 0.990 | -0.058 | 1.000 | 0.199 | 1.000 | -0.095 | 0.906 | -0.519 | 0.423 | -0.543 | 0.437 | **-1.256** | **0.006** | Possible ESAT-6 like protein EsxD | cell wall and cell processes |
| **Rv3616c** | *espA* | -0.554 | 0.536 | -0.236 | 0.990 | -0.005 | 1.000 | -0.157 | 1.000 | -0.781 | 0.175 | -0.482 | 0.447 | -0.388 | 0.609 | **-1.075** | **0.020** | ESX-1 secretion-associated protein A, EspA | cell wall and cell processes |
| **Rv0072** | *Rv0072* | 0.241 | 0.702 | 0.080 | 0.990 | **5.728** | **0.048** | 0.000 | 1.000 | -0.361 | 0.321 | -0.477 | 0.120 | 0.000 | 1.000 | 0.000 | 1.000 | Probable glutamine-transport transmembrane protein ABC transporter | cell wall and cell processes |
| **Rv2347c** | *esxP* | -0.549 | 0.330 | -0.219 | 0.990 | -0.205 | 1.000 | -0.104 | 1.000 | 0.110 | 0.854 | -0.465 | 0.293 | -0.371 | 0.498 | **-1.144** | **0.001** | Putative ESAT-6 like protein EsxP (ESAT-6 like protein 7) | cell wall and cell processes |
| **Rv1362c** | *Rv1362c* | -0.308 | 0.658 | 0.015 | 0.998 | -0.267 | 1.000 | 0.006 | 1.000 | -0.501 | 0.230 | -0.447 | 0.350 | -0.604 | 0.244 | **-1.042** | **0.001** | Possible membrane protein | cell wall and cell processes |
| **Rv2080** | *lppJ* | -0.015 | NA | -0.177 | NA | 0.779 | 1.000 | -0.676 | 1.000 | -0.208 | 0.723 | -0.421 | 0.512 | **1.714** | **0.059** | -0.734 | 0.193 | Lipoprotein LppJ | cell wall and cell processes |
| **Rv3614c** | *espD* | -0.478 | 0.680 | -0.486 | 0.989 | -0.010 | 1.000 | -0.506 | 1.000 | -0.478 | 0.501 | -0.383 | 0.609 | -0.203 | 0.838 | **-1.426** | **0.004** | ESX-1 secretion-associated protein EspD | cell wall and cell processes |
| **Rv3615c** | *espC* | -0.803 | 0.297 | -0.310 | 0.990 | -0.056 | 1.000 | -0.478 | 1.000 | -0.595 | 0.362 | -0.355 | 0.626 | -0.152 | 0.882 | **-1.369** | **0.005** | ESX-1 secretion-associated protein EspC | cell wall and cell processes |
| **Rv3238c** | *Rv3238c* | -0.058 | 0.969 | 0.192 | 0.990 | -0.186 | 1.000 | 0.175 | 1.000 | -0.002 | 0.996 | -0.299 | 0.479 | -0.039 | 0.963 | **-1.104** | **0.000** | Probable conserved integral membrane protein | cell wall and cell processes |
| **Rv1342c** | *Rv1342c* | -0.039 | NA | 0.107 | NA | -0.304 | 1.000 | -0.096 | 1.000 | -0.271 | 0.743 | -0.282 | 0.689 | -0.638 | 0.358 | **-1.075** | **0.022** | Conserved membrane protein | cell wall and cell processes |
| **Rv2254c** | *Rv2254c* | -1.071 | NA | 0.018 | NA | 0.112 | 1.000 | -0.195 | 1.000 | -0.072 | 0.949 | -0.256 | 0.662 | **-1.104** | **0.086** | -0.710 | 0.067 | Probable integral membrane protein | cell wall and cell processes |
| **Rv1037c** | *esxI* | -0.639 | 0.350 | -0.235 | 0.990 | -0.024 | 1.000 | -0.280 | 1.000 | 0.569 | 0.295 | -0.206 | 0.834 | **1.515** | **0.059** | 0.087 | 0.933 | Putative ESAT-6 like protein EsxI (ESAT-6 like protein 1) | cell wall and cell processes |
| **Rv0261c** | *narK3* | -0.398 | 0.283 | -0.031 | 0.993 | -0.028 | 1.000 | -0.326 | 1.000 | 0.388 | 0.245 | -0.087 | 0.905 | -0.307 | 0.660 | **-1.551** | **0.000** | Probable integral membrane nitrite extrusion protein NarK3 (nitrite facilitator) | cell wall and cell processes |

| | | | | | | | | | | | | | | | | | | | |
|---|---|---|---|---|---|---|---|---|---|---|---|---|---|---|---|---|---|---|---|
| **Rv2136c** | *Rv2136c* | 1.015 | 0.002 | 0.284 | 0.987 | 0.086 | 1.000 | 0.258 | 1.000 | -0.366 | 0.383 | -0.009 | 0.988 | -0.041 | 0.960 | -0.380 | 0.246 | Possible conserved transmembrane protein | cell wall and cell processes |
| **Rv0531** | *Rv0531* | -0.014 | NA | 0.193 | NA | 0.002 | 1.000 | 0.155 | 1.000 | -1.102 | 0.095 | 0.004 | 0.996 | -0.230 | 0.800 | 0.437 | 0.373 | Possible conserved membrane protein | cell wall and cell processes |
| **Rv2157c** | *murF* | 0.036 | 0.982 | 0.222 | 0.990 | -0.119 | 1.000 | -0.009 | 1.000 | -0.514 | 0.094 | 0.025 | 0.955 | 0.248 | 0.585 | 1.155 | 0.000 | Probable UDP-N-acetylmuramoylalanyl-D-glutamyl-2,6-diaminopimelate-D-alanyl-D-alanyl ligase MurF | cell wall and cell processes |
| **Rv1183** | *mmpL10* | 0.205 | 0.886 | -0.024 | 0.994 | -0.033 | 1.000 | 0.011 | 1.000 | -0.763 | 0.138 | 0.054 | 0.944 | 1.873 | 0.000 | 1.613 | 0.000 | MmpL10 | cell wall and cell processes |
| **Rv0867c** | *rpfA* | -1.154 | 0.007 | -0.401 | 0.956 | -0.037 | 1.000 | -0.340 | 1.000 | 0.359 | 0.488 | 0.084 | 0.880 | -0.703 | 0.102 | 0.197 | 0.638 | Possible resuscitation-promoting factor RpfA | cell wall and cell processes |
| **Rv3823c** | *mmpL8* | 0.185 | 0.912 | 0.112 | 0.992 | 0.101 | 1.000 | 0.010 | 1.000 | -0.932 | 0.087 | 0.185 | 0.813 | 1.732 | 0.001 | 1.677 | 0.000 | Conserved integral membrane transport protein MmpL8 | cell wall and cell processes |
| **Rv2158c** | *murE* | -0.501 | 0.417 | 0.159 | 0.990 | -0.165 | 1.000 | 0.030 | 1.000 | -0.232 | 0.671 | 0.185 | 0.734 | 0.691 | 0.187 | 1.031 | 0.004 | Probable UDP-N-acetylmuramoylalanyl-D-glutamate-2,6-diaminopimelate ligase MurE | cell wall and cell processes |
| **Rv0114** | *gmhB* | -0.599 | 0.090 | -0.261 | 0.990 | -0.358 | 1.000 | 0.235 | 1.000 | 0.989 | 0.001 | 0.239 | 0.689 | 1.017 | 0.016 | 0.323 | 0.368 | phosphatase GmhB (D-glycero-D-manno-heptose 7-phosphate kinase) | cell wall and cell processes |
| **Rv3330** | *dacB1* | 1.196 | 0.300 | -0.068 | 0.992 | -0.092 | 1.000 | -0.068 | 1.000 | 1.307 | 0.011 | 0.303 | 0.614 | 0.466 | 0.469 | 0.482 | 0.196 | Probable penicillin-binding protein DacB1 (D-alanyl-D-alanine carboxypeptidase) (DD-peptidase) (DD-carboxypeptidase) (PBP) (DD-transpeptidase) (serine-type D-ala-D-ala carboxypeptidase) (D-amino acid hydrolase) | cell wall and cell processes |
| **Rv0288** | *esxH* | -0.163 | 0.824 | 0.077 | 0.992 | -0.008 | 1.000 | 0.223 | 1.000 | 0.715 | 0.010 | 0.368 | 0.255 | 0.522 | 0.128 | 1.049 | 0.000 | Low molecular weight protein antigen 7 EsxH (10 kDa antigen) (CFP-7) (protein TB10.4) | cell wall and cell processes |
| **Rv0488** | *Rv0488* | 0.610 | NA | -0.152 | NA | -0.134 | 1.000 | 0.411 | 1.000 | 1.140 | 0.039 | 0.467 | 0.521 | 0.658 | 0.391 | -0.150 | 0.815 | Probable conserved integral membrane protein | cell wall and cell processes |
| **Rv3807c** | *Rv3807c* | -0.102 | 0.930 | 0.039 | 0.992 | -0.082 | 1.000 | -0.033 | 1.000 | 0.366 | 0.330 | 0.491 | 0.073 | 0.036 | 0.959 | 1.081 | 0.000 | Possible conserved transmembrane protein | cell wall and cell processes |
| **Rv0290** | *eccD3* | -0.241 | 0.877 | 0.157 | 0.990 | 0.033 | 1.000 | 0.123 | 1.000 | 0.291 | 0.701 | 0.689 | 0.254 | 0.011 | 0.997 | 1.115 | 0.021 | ESX conserved component EccD3. ESX-3 type VII secretion system protein. Probable transmembrane protein. | cell wall and cell processes |
| **Rv0528** | *Rv0528* | 0.202 | 0.851 | 0.143 | 0.990 | 0.045 | 1.000 | 0.095 | 1.000 | -0.293 | 0.569 | 0.711 | 0.081 | 0.119 | 0.870 | 1.117 | 0.001 | Probable conserved transmembrane protein | cell wall and cell processes |
| **Rv1227c** | *Rv1227c* | 0.705 | 0.289 | -0.219 | 0.990 | 0.028 | 1.000 | 0.399 | 1.000 | 0.371 | 0.501 | 0.715 | 0.054 | 0.797 | 0.095 | 1.039 | 0.000 | Probable transmembrane protein | cell wall and cell processes |
| **Rv0236c** | *aftD* | -0.014 | 0.998 | 0.027 | 0.992 | -0.091 | 1.000 | 0.012 | 1.000 | 0.434 | 0.396 | 0.794 | 0.055 | 0.233 | 0.710 | 1.103 | 0.001 | Possible arabinofuranosyltransferase AftD | cell wall and cell processes |
| **Rv1198** | *esxL* | -0.350 | 0.680 | 0.022 | 0.994 | 0.085 | 1.000 | -0.035 | 1.000 | 0.734 | 0.088 | 0.808 | 0.057 | 0.751 | 0.112 | 1.100 | 0.002 | Putative ESAT-6 like protein EsxL (ESAT-6 like protein 4) | cell wall and cell processes |
| **Rv3270** | *ctpC* | 1.319 | 0.234 | 0.390 | 0.990 | 0.256 | 1.000 | 0.231 | 1.000 | 1.636 | 0.052 | 0.816 | 0.411 | 0.176 | 0.913 | 1.225 | 0.110 | Probable metal cation-transporting P-type ATPase C CtpC | cell wall and cell processes |
| **Rv0935** | *pstC1* | 0.093 | 0.946 | 0.058 | 0.992 | -0.122 | 1.000 | 0.043 | 1.000 | 0.102 | 0.854 | 0.837 | 0.016 | 0.076 | 0.912 | 1.037 | 0.001 | PstC1 | cell wall and cell processes |
| **Rv1226c** | *Rv1226c* | 0.735 | 0.086 | 0.190 | 0.990 | 0.089 | 1.000 | 0.137 | 1.000 | 0.534 | 0.151 | 0.888 | 0.004 | 0.647 | 0.097 | 1.080 | 0.000 | Probable transmembrane protein | cell wall and cell processes |
| **Rv2625c** | *Rv2625c* | -0.433 | 0.680 | -0.045 | 0.992 | 0.330 | 1.000 | 0.411 | 1.000 | 0.785 | 0.158 | 1.052 | 0.044 | 0.529 | 0.434 | 0.208 | 0.731 | Probable conserved transmembrane alanine and leucine rich protein | cell wall and cell processes |
| **Rv0267** | *narU* | -1.187 | 0.223 | -0.625 | 0.990 | 0.170 | 1.000 | -0.507 | 1.000 | 0.998 | 0.243 | 1.061 | 0.273 | 1.897 | 0.068 | 0.301 | 0.754 | Probable integral membrane nitrite extrusion protein NarU (nitrite facilitator) | cell wall and cell processes |
| **Rv1349** | *irtB* | 0.893 | 0.319 | -0.201 | 0.990 | -0.072 | 1.000 | -0.213 | 1.000 | 0.922 | 0.096 | 1.112 | 0.033 | 0.695 | 0.311 | 0.611 | 0.207 | Iron-regulated transporter IrtB | cell wall and cell processes |
| **Rv1686c** | *Rv1686c* | 0.621 | 0.846 | 0.018 | NA | -0.313 | 1.000 | -0.356 | 1.000 | 0.739 | 0.651 | 1.564 | 0.243 | 2.378 | 0.073 | 1.507 | 0.170 | transporter | cell wall and cell processes |
| **Rv1687c** | *Rv1687c* | 0.084 | 0.990 | 0.314 | 0.992 | 0.066 | 1.000 | 0.240 | 1.000 | 1.258 | 0.448 | 1.707 | 0.253 | 2.859 | 0.052 | 1.799 | 0.153 | Probable conserved ATP-binding protein ABC transporter | cell wall and cell processes |
| **Rv3354** | *Rv3354* | -0.176 | NA | -0.524 | NA | -0.747 | 1.000 | -0.391 | 1.000 | -0.197 | 0.878 | -1.854 | 0.016 | -0.819 | 0.419 | -1.177 | 0.101 | Conserved hypothetical protein | conserved hypotheticals |
| **Rv0250c** | *Rv0250c* | -0.054 | 0.983 | 0.118 | 0.992 | 0.293 | 1.000 | -0.146 | 1.000 | -2.218 | 0.000 | -1.726 | 0.000 | -1.449 | 0.003 | -1.603 | 0.000 | Conserved protein | conserved hypotheticals |
| **Rv3642c** | *Rv3642c* | -0.120 | 0.915 | 0.416 | 0.964 | 0.179 | 1.000 | 0.301 | 1.000 | -1.197 | 0.003 | -1.527 | 0.000 | -0.631 | 0.242 | -0.596 | 0.122 | Hypothetical protein | conserved hypotheticals |
| **Rv1772** | *Rv1772* | -0.401 | 0.776 | -0.131 | 0.992 | -0.078 | 1.000 | -0.153 | 1.000 | -0.968 | 0.141 | -1.505 | 0.010 | -1.275 | 0.060 | -1.375 | 0.010 | Hypothetical protein | conserved hypotheticals |
| **Rv0307c** | *Rv0307c* | 0.161 | 0.894 | 0.149 | 0.990 | -0.083 | 1.000 | 0.154 | 1.000 | -0.598 | 0.229 | -1.455 | 0.001 | -0.561 | 0.385 | -1.209 | 0.002 | Unknown protein | conserved hypotheticals |
| **Rv3209** | *Rv3209* | -0.025 | 0.993 | 0.322 | 0.686 | 0.100 | 1.000 | 0.302 | 1.000 | 0.111 | 0.896 | -1.450 | 0.000 | -0.715 | 0.038 | -1.549 | 0.000 | Conserved hypothetical threonine and proline rich protein | conserved hypotheticals |
| **Rv3288c** | *usfY* | 0.450 | 0.491 | 0.173 | 0.990 | -0.085 | 1.000 | 0.258 | 1.000 | -0.524 | 0.247 | -1.374 | 0.000 | -0.636 | 0.177 | -1.555 | 0.000 | Putative protein UsfY | conserved hypotheticals |
| **Rv1073** | *Rv1073* | -0.336 | 0.650 | -0.160 | 0.990 | -0.105 | 1.000 | -0.197 | 1.000 | -0.923 | 0.011 | -1.277 | 0.000 | -1.108 | 0.003 | -1.444 | 0.000 | Conserved hypothetical protein | conserved hypotheticals |
| **Rv3489** | *Rv3489* | 0.047 | 0.976 | 0.323 | 0.965 | -0.047 | 1.000 | 0.094 | 1.000 | -1.405 | 0.000 | -1.264 | 0.000 | -0.716 | 0.081 | -0.729 | 0.019 | Unknown protein | conserved hypotheticals |
| **Rv0264c** | *Rv0264c* | -0.383 | 0.724 | 0.208 | 0.990 | -0.031 | 1.000 | -0.156 | 1.000 | -1.023 | 0.046 | -1.247 | 0.006 | -1.600 | 0.000 | -2.033 | 0.000 | Conserved hypothetical protein | conserved hypotheticals |
| **Rv1331** | *Rv1331* | 0.231 | 0.636 | 0.311 | 0.855 | 0.118 | 1.000 | 0.442 | 1.000 | -1.183 | 0.000 | -1.206 | 0.000 | -0.647 | 0.021 | -1.037 | 0.000 | Conserved hypothetical protein | conserved hypotheticals |
| **Rv1057** | *Rv1057* | 0.248 | 0.707 | 0.159 | 0.990 | 0.029 | 1.000 | 0.185 | 1.000 | -0.375 | 0.302 | -1.193 | 0.000 | -0.968 | 0.002 | -1.447 | 0.000 | Conserved hypothetical protein | conserved hypotheticals |
| **Rv0692** | *Rv0692* | 0.026 | 0.993 | 0.461 | 0.990 | 0.274 | 1.000 | 0.247 | 1.000 | -2.097 | 0.000 | -1.167 | 0.035 | -0.632 | 0.372 | -1.174 | 0.015 | Conserved hypothetical protein | conserved hypotheticals |
| **Rv0207c** | *Rv0207c* | 0.001 | 1.000 | 0.125 | 0.990 | 0.034 | 1.000 | 0.001 | 1.000 | -0.928 | 0.021 | -1.166 | 0.003 | -1.042 | 0.014 | -1.495 | 0.000 | Conserved hypothetical protein | conserved hypotheticals |
| **Rv1159A** | *Rv1159A* | 0.163 | 0.893 | 0.390 | 0.966 | -0.052 | 1.000 | 0.378 | 1.000 | -0.554 | 0.231 | -1.146 | 0.004 | -0.454 | 0.419 | -1.203 | 0.001 | Unknown protein | conserved hypotheticals |
| **Rv0885** | *Rv0885* | -0.681 | 0.330 | -0.119 | 0.992 | -0.009 | 1.000 | -0.252 | 1.000 | -0.836 | 0.107 | -1.120 | 0.020 | -1.022 | 0.061 | -1.190 | 0.005 | Conserved hypothetical protein | conserved hypotheticals |
| **Rv0028** | *Rv0028* | 0.072 | NA | 0.270 | NA | 0.313 | 1.000 | 0.171 | 1.000 | -0.005 | NA | -1.113 | 0.140 | -0.572 | NA | -1.535 | 0.006 | Conserved hypothetical protein | conserved hypotheticals |
| **Rv1870c** | *Rv1870c* | 0.387 | 0.279 | 0.123 | 0.990 | -0.064 | 1.000 | 0.070 | 1.000 | 0.236 | 0.461 | -1.089 | 0.000 | -0.775 | 0.001 | -1.301 | 0.000 | Conserved hypothetical protein | conserved hypotheticals |
| **Rv0263c** | *Rv0263c* | -0.335 | 0.724 | 0.046 | 0.992 | 0.119 | 1.000 | -0.073 | 1.000 | -1.235 | 0.003 | -1.082 | 0.009 | -1.731 | 0.000 | -2.386 | 0.000 | Conserved hypothetical protein | conserved hypotheticals |
| **Rv1995** | *Rv1995* | -0.457 | 0.796 | -0.350 | 0.990 | 0.219 | 1.000 | 0.056 | 1.000 | -0.248 | 0.807 | -1.032 | 0.130 | -0.708 | 0.434 | -1.098 | 0.065 | Unknown protein | conserved hypotheticals |

| | | | | | | | | | | | | | | | | | | | |
|---|---|---|---|---|---|---|---|---|---|---|---|---|---|---|---|---|---|---|---|
| Rv0762c | Rv0762c | -0.136 | 0.937 | 0.035 | 0.992 | -0.131 | 1.000 | 0.059 | 1.000 | -0.356 | 0.558 | -1.016 | 0.042 | -0.013 | 0.995 | -0.634 | 0.157 | Conserved hypothetical protein | conserved hypotheticals |
| Rv0856 | Rv0856 | 0.204 | 0.775 | 0.255 | 0.990 | 0.013 | 1.000 | -0.003 | 1.000 | -0.604 | 0.072 | -0.996 | 0.001 | -0.570 | 0.130 | -1.050 | 0.000 | Conserved hypothetical protein | conserved hypotheticals |
| Rv3292 | Rv3292 | 0.245 | 0.753 | -0.001 | 0.999 | -0.076 | 1.000 | 0.011 | 1.000 | -0.285 | 0.504 | -0.983 | 0.000 | -0.485 | 0.183 | -1.039 | 0.000 | Conserved hypothetical protein | conserved hypotheticals |
| Rv2293c | Rv2293c | -0.242 | 0.894 | -0.362 | 0.990 | -0.068 | 1.000 | -0.402 | 1.000 | -0.343 | 0.708 | -0.919 | 0.205 | -0.593 | 0.533 | -1.038 | 0.082 | Conserved hypothetical protein | conserved hypotheticals |
| Rv2166c | Rv2166c | -0.259 | 0.762 | -0.083 | 0.992 | -0.040 | 1.000 | -0.002 | 1.000 | -0.178 | 0.757 | -0.918 | 0.004 | -0.793 | 0.033 | -1.158 | 0.000 | Conserved protein | conserved hypotheticals |
| Rv3412 | Rv3412 | 0.321 | 0.600 | 0.088 | 0.992 | -0.034 | 1.000 | 0.146 | 1.000 | -0.664 | 0.048 | -0.901 | 0.004 | -0.601 | 0.114 | -1.034 | 0.000 | Conserved hypothetical protein | conserved hypotheticals |
| Rv3142c | Rv3142c | 0.073 | 0.944 | 0.181 | 0.990 | -0.054 | 1.000 | 0.291 | 1.000 | -1.012 | 0.000 | -0.890 | 0.002 | -0.578 | 0.112 | -1.175 | 0.000 | Hypothetical protein | conserved hypotheticals |
| Rv2331 | Rv2331 | -0.133 | 0.958 | -0.194 | 0.990 | 0.171 | 1.000 | 0.355 | 1.000 | -1.468 | 0.020 | -0.876 | 0.385 | -0.196 | 0.888 | 0.147 | 0.870 | Hypothetical protein | conserved hypotheticals |
| Rv1904 | Rv1904 | -0.456 | 0.551 | -0.162 | 0.990 | -0.042 | 1.000 | -0.254 | 1.000 | -0.664 | 0.199 | -0.825 | 0.066 | -0.697 | 0.180 | -1.244 | 0.001 | Conserved hypothetical protein | conserved hypotheticals |
| Rv3237c | Rv3237c | 0.037 | 0.982 | 0.157 | 0.990 | 0.126 | 1.000 | 0.156 | 1.000 | -0.986 | 0.002 | -0.822 | 0.012 | -0.361 | 0.421 | -1.089 | 0.000 | Conserved protein | conserved hypotheticals |
| Rv3733c | Rv3733c | -0.310 | 0.697 | -0.084 | 0.992 | 0.088 | 1.000 | 0.105 | 1.000 | -0.622 | 0.141 | -0.745 | 0.031 | -0.464 | 0.308 | -1.387 | 0.000 | Conserved hypothetical protein | conserved hypotheticals |
| Rv2042c | Rv2042c | 1.001 | 0.000 | 0.485 | 0.175 | 0.333 | 1.000 | 0.210 | 1.000 | -0.892 | 0.000 | -0.710 | 0.003 | -0.501 | 0.057 | -0.942 | 0.000 | Conserved protein | conserved hypotheticals |
| Rv2143 | Rv2143 | 0.280 | 0.636 | -0.008 | 0.998 | 0.046 | 1.000 | 0.191 | 1.000 | -0.276 | 0.488 | -0.687 | 0.042 | -0.306 | 0.513 | -1.068 | 0.000 | Conserved hypothetical protein | conserved hypotheticals |
| Rv3005c | Rv3005c | 0.099 | 0.915 | 0.190 | 0.990 | 0.049 | 1.000 | 0.138 | 1.000 | -0.651 | 0.036 | -0.675 | 0.005 | -0.677 | 0.012 | -1.008 | 0.000 | Conserved hypothetical protein | conserved hypotheticals |
| Rv2323c | Rv2323c | -1.159 | 0.010 | 0.159 | 0.990 | -0.421 | 1.000 | 0.130 | 1.000 | 0.098 | 0.885 | -0.665 | 0.205 | -0.763 | 0.249 | -0.402 | 0.424 | Conserved protein | conserved hypotheticals |
| Rv3073c | Rv3073c | 0.020 | 0.992 | -0.219 | 0.990 | -0.028 | 1.000 | -0.108 | 1.000 | -0.604 | 0.180 | -0.655 | 0.147 | -0.092 | 0.910 | -1.042 | 0.001 | Conserved hypothetical protein | conserved hypotheticals |
| Rv2990c | Rv2990c | -0.412 | 0.426 | -0.100 | 0.990 | -0.072 | 1.000 | -0.296 | 1.000 | -1.011 | 0.001 | -0.655 | 0.055 | -0.898 | 0.008 | -1.140 | 0.000 | Hypothetical protein | conserved hypotheticals |
| Rv1883c | Rv1883c | 0.437 | 0.097 | -0.096 | 0.990 | 0.023 | 1.000 | -0.115 | 1.000 | -0.555 | 0.039 | -0.654 | 0.002 | -0.759 | 0.001 | -1.572 | 0.000 | Conserved hypothetical protein | conserved hypotheticals |
| Rv0679c | Rv0679c | -0.065 | 0.972 | 0.032 | 0.992 | 0.101 | 1.000 | 0.086 | 1.000 | -0.531 | 0.320 | -0.635 | 0.169 | -0.807 | 0.095 | -1.181 | 0.000 | Conserved threonine rich protein | conserved hypotheticals |
| Rv3717 | Rv3717 | 0.179 | 0.952 | 0.159 | 0.990 | 0.047 | 1.000 | 0.087 | 1.000 | 1.050 | 0.131 | -0.603 | 0.258 | -0.253 | 0.772 | -1.084 | 0.008 | Conserved hypothetical protein | conserved hypotheticals |
| Rv0530A | Rv0530A | 0.232 | NA | 0.086 | NA | 0.064 | 1.000 | 0.130 | 1.000 | -0.623 | 0.320 | -0.600 | 0.387 | -1.168 | 0.095 | -1.363 | 0.007 | Conserved protein | conserved hypotheticals |
| Rv3486 | Rv3486 | 1.390 | 0.056 | 0.252 | 0.990 | 0.213 | 1.000 | 0.249 | 1.000 | 0.068 | 0.936 | -0.578 | 0.160 | -0.079 | 0.927 | -0.829 | 0.014 | Conserved protein | conserved hypotheticals |
| Rv0470A | Rv0470A | 0.021 | NA | 0.634 | NA | 0.530 | 1.000 | 0.565 | 1.000 | -0.052 | 0.952 | -0.553 | 0.344 | -1.153 | 0.088 | -0.705 | 0.109 | Hypothetical protein | conserved hypotheticals |
| Rv3633 | Rv3633 | -0.689 | 0.401 | -0.514 | 0.968 | -0.193 | 1.000 | -0.304 | 1.000 | -0.986 | 0.079 | -0.537 | 0.402 | -0.628 | 0.368 | -1.422 | 0.002 | Conserved protein | conserved hypotheticals |
| Rv1810 | Rv1810 | -0.095 | 0.953 | 0.036 | 0.992 | 0.020 | 1.000 | 0.100 | 1.000 | -0.729 | 0.098 | -0.526 | 0.256 | -0.638 | 0.211 | -1.445 | 0.000 | Conserved protein | conserved hypotheticals |
| Rv0500A | Rv0500A | -0.608 | 0.373 | -0.319 | NA | 0.076 | 1.000 | -0.195 | 1.000 | 0.125 | 0.872 | -0.517 | 0.288 | -0.821 | 0.130 | -1.010 | 0.009 | Conserved protein | conserved hypotheticals |
| Rv1265 | Rv1265 | -0.445 | 0.417 | -0.406 | 0.913 | -0.474 | 1.000 | 0.006 | 1.000 | -1.106 | 0.001 | -0.497 | 0.205 | -0.531 | 0.228 | -0.798 | 0.011 | Unknown protein | conserved hypotheticals |
| Rv0580c | Rv0580c | -0.206 | 0.809 | 0.168 | 0.990 | -0.065 | 1.000 | 0.172 | 1.000 | -0.611 | 0.152 | -0.490 | 0.271 | -0.474 | 0.399 | -1.096 | 0.001 | Conserved protein | conserved hypotheticals |
| Rv3190A | Rv3190A | -0.116 | 0.922 | 0.258 | 0.990 | -0.228 | 1.000 | 0.150 | 1.000 | -1.100 | 0.008 | -0.474 | 0.389 | -0.774 | 0.190 | -0.285 | 0.436 | Conserved protein | conserved hypotheticals |
| Rv1501 | Rv1501 | 0.325 | 0.750 | -0.221 | 0.990 | -0.135 | 1.000 | -0.096 | 1.000 | 0.166 | 0.803 | -0.471 | 0.387 | -0.386 | 0.570 | -1.319 | 0.001 | Conserved hypothetical protein | conserved hypotheticals |
| Rv2517c | Rv2517c | -0.094 | 0.969 | 0.026 | 0.994 | 0.050 | 1.000 | 0.188 | 1.000 | -1.200 | 0.021 | -0.445 | 0.463 | 0.370 | 0.602 | 0.037 | 0.964 | Unknown protein | conserved hypotheticals |
| Rv1489A | Rv1489A | 0.251 | NA | 0.130 | NA | -0.193 | 1.000 | -0.005 | 1.000 | -0.879 | 0.060 | -0.400 | 0.530 | -0.857 | 0.151 | -1.299 | 0.002 | Conserved protein | conserved hypotheticals |
| Rv2225 | panB | -1.148 | 0.001 | -0.704 | 0.348 | -0.036 | 1.000 | -0.641 | 1.000 | -0.262 | 0.600 | -0.391 | 0.364 | -0.677 | 0.115 | 0.321 | 0.385 | Conserved protein | conserved hypotheticals |
| Rv3046c | Rv3046c | 0.294 | NA | 0.122 | NA | 0.732 | 1.000 | 0.068 | 1.000 | -0.458 | 0.520 | -0.385 | 0.589 | -0.098 | 0.930 | -1.259 | 0.001 | Conserved protein | conserved hypotheticals |
| Rv1754c | Rv1754c | 0.417 | 0.771 | -0.560 | 0.839 | -0.103 | 1.000 | -0.400 | 1.000 | 0.439 | 0.482 | -0.378 | 0.543 | -0.816 | 0.144 | -1.022 | 0.010 | Conserved protein | conserved hypotheticals |
| Rv2663 | Rv2663 | -0.139 | 0.911 | -0.070 | 0.992 | -0.389 | 1.000 | 0.161 | 1.000 | -1.019 | 0.009 | -0.352 | 0.456 | -0.308 | 0.583 | -1.131 | 0.001 | Hypothetical protein | conserved hypotheticals |
| Rv0424c | Rv0424c | -0.140 | 0.914 | -0.206 | 0.990 | 0.312 | 1.000 | -0.015 | 1.000 | -1.213 | 0.002 | -0.338 | 0.484 | -0.561 | 0.234 | -0.318 | 0.306 | Hypothetical protein | conserved hypotheticals |
| Rv1907c | Rv1907c | -0.336 | 0.391 | 0.130 | 0.990 | -0.147 | 1.000 | 0.050 | 1.000 | -1.102 | 0.000 | -0.316 | 0.286 | -0.770 | 0.003 | -0.160 | 0.529 | Hypothetical protein | conserved hypotheticals |
| Rv0057 | Rv0057 | 0.034 | 0.983 | 0.038 | 0.992 | -0.041 | 1.000 | -0.039 | 1.000 | 1.155 | 0.000 | -0.268 | 0.488 | -0.514 | 0.161 | 0.240 | 0.352 | Hypothetical protein | conserved hypotheticals |
| Rv1171 | Rv1171 | -0.076 | 0.970 | 0.102 | 0.990 | -0.149 | 1.000 | -0.085 | 1.000 | -1.020 | 0.043 | -0.252 | 0.625 | -0.157 | 0.805 | -0.644 | 0.073 | Conserved hypothetical protein | conserved hypotheticals |
| Rv1724c | Rv1724c | 0.536 | NA | 0.261 | NA | -0.220 | 1.000 | 0.440 | 1.000 | -0.259 | NA | -0.245 | 0.783 | -0.097 | NA | -1.202 | 0.019 | Hypothetical protein | conserved hypotheticals |
| Rv2478c | Rv2478c | 1.099 | NA | -0.218 | NA | -0.733 | 1.000 | 0.661 | 1.000 | 0.877 | 0.212 | -0.188 | 0.817 | 3.810 | 0.016 | 1.074 | 0.046 | Conserved hypothetical protein | conserved hypotheticals |
| Rv3612c | Rv3612c | 0.621 | 0.592 | -0.537 | 0.987 | -0.039 | 1.000 | -0.527 | 1.000 | 0.088 | 0.936 | -0.179 | 0.843 | -1.186 | 0.099 | -1.952 | 0.000 | Conserved hypothetical protein | conserved hypotheticals |
| Rv2360c | Rv2360c | 1.004 | 0.025 | 0.262 | 0.976 | -0.033 | 1.000 | 0.043 | 1.000 | 0.265 | 0.636 | -0.130 | 0.798 | -0.652 | 0.184 | -0.124 | 0.754 | Unknown protein | conserved hypotheticals |
| Rv1535 | Rv1535 | -2.068 | 0.000 | -0.791 | 0.641 | -0.308 | 1.000 | -0.131 | 1.000 | 0.060 | 0.939 | -0.116 | 0.863 | 0.002 | 1.000 | -0.671 | 0.180 | Unknown protein | conserved hypotheticals |
| Rv0963c | Rv0963c | 0.607 | NA | 0.596 | NA | -0.904 | 1.000 | 0.195 | 1.000 | 0.645 | 0.423 | -0.087 | 0.939 | 1.770 | 0.033 | 0.235 | 0.766 | Conserved hypothetical protein | conserved hypotheticals |
| Rv0460 | Rv0460 | -0.451 | NA | -0.516 | NA | -0.652 | 1.000 | -0.319 | 1.000 | -0.170 | NA | -0.082 | 0.942 | -0.319 | NA | -1.566 | 0.011 | Conserved hydrophobic protein | conserved hypotheticals |
| Rv2664 | Rv2664 | -0.832 | 0.390 | -0.406 | 0.990 | -0.053 | 1.000 | -0.028 | 1.000 | -0.772 | 0.305 | -0.074 | 0.914 | -0.644 | 0.258 | -1.166 | 0.004 | Hypothetical protein | conserved hypotheticals |
| Rv0739 | Rv0739 | -0.413 | NA | -0.073 | NA | -0.271 | 1.000 | 0.007 | 1.000 | 0.353 | 0.665 | -0.046 | 0.964 | 0.060 | 0.968 | -1.156 | 0.008 | Conserved hypothetical protein | conserved hypotheticals |
| Rv1638A | Rv1638A | -0.536 | NA | -0.706 | NA | -0.191 | 1.000 | 0.257 | 1.000 | -1.323 | 0.024 | 0.005 | 0.996 | 0.264 | 0.853 | -0.532 | 0.451 | Conserved hypothetical protein | conserved hypotheticals |
| Rv1190 | Rv1190 | -0.184 | NA | -0.293 | NA | 0.000 | 1.000 | 0.395 | 1.000 | -0.037 | 0.968 | 0.053 | 0.951 | 0.000 | 1.000 | -1.008 | 0.008 | Conserved hypothetical protein | conserved hypotheticals |
| Rv0140 | Rv0140 | 0.001 | 1.000 | -0.051 | 0.992 | 0.069 | 1.000 | 0.149 | 1.000 | -1.410 | 0.040 | 0.106 | 0.915 | 0.493 | 0.619 | -0.037 | 0.976 | Conserved protein | conserved hypotheticals |
| Rv3463 | Rv3463 | -0.331 | 0.858 | -0.590 | 0.990 | -0.452 | 1.000 | -0.321 | 1.000 | -1.211 | 0.080 | 0.107 | 0.910 | 0.505 | 0.585 | 0.427 | 0.532 | Conserved protein | conserved hypotheticals |
| Rv0059 | Rv0059 | 0.314 | 0.807 | 0.089 | 0.990 | -0.255 | 1.000 | -0.065 | 1.000 | 1.132 | 0.015 | 0.111 | 0.859 | -0.582 | 0.269 | 0.517 | 0.069 | Hypothetical protein | conserved hypotheticals |
| Rv2466c | Rv2466c | -0.572 | 0.685 | -0.214 | 0.990 | -0.192 | 1.000 | 0.072 | 1.000 | -1.375 | 0.045 | 0.121 | 0.904 | 0.424 | 0.677 | 0.449 | 0.530 | Conserved protein | conserved hypotheticals |

| | | | | | | | | | | | | | | | | | | | |
|---|---|---|---|---|---|---|---|---|---|---|---|---|---|---|---|---|---|---|---|
| Rv1268c | Rv1268c | 0.552 | NA | -0.372 | NA | -0.040 | 1.000 | -0.453 | 1.000 | 0.174 | 0.884 | 0.122 | 0.919 | 0.017 | 0.995 | -1.286 | 0.022 | Hypothetical protein | conserved hypotheticals |
| Rv2247 | accD6 | -0.412 | 0.509 | -0.118 | 0.990 | -0.052 | 1.000 | 0.114 | 1.000 | 0.378 | 0.399 | 0.159 | 0.767 | 0.228 | 0.692 | 1.012 | 0.002 | Conserved hypothetical protein | conserved hypotheticals |
| Rv2023c | Rv2023c | -0.315 | NA | -0.070 | NA | 0.000 | 1.000 | 0.207 | 1.000 | -0.606 | 0.214 | 0.226 | 0.792 | -0.591 | 0.227 | -1.006 | 0.001 | Hypothetical protein | conserved hypotheticals |
| Rv1078 | Rv1078 | -1.041 | 0.001 | -0.631 | 0.208 | 0.001 | 1.000 | -0.303 | 1.000 | 0.189 | 0.671 | 0.232 | 0.561 | 0.153 | 0.757 | 0.359 | 0.188 | Probable proline-rich antigen homolog Pra | conserved hypotheticals |
| Rv2407 | Rv2407 | 0.131 | NA | 0.051 | NA | -0.152 | 1.000 | 0.195 | 1.000 | 1.144 | 0.076 | 0.275 | 0.757 | 0.277 | 0.776 | 0.019 | 0.985 | Conserved hypothetical protein | conserved hypotheticals |
| Rv2288 | Rv2288 | 0.035 | NA | -0.229 | NA | -0.583 | 1.000 | 0.328 | 1.000 | -0.179 | 0.794 | 0.329 | 0.754 | 1.762 | 0.078 | -0.292 | 0.663 | Hypothetical protein | conserved hypotheticals |
| Rv2189c | Rv2189c | -0.454 | 0.780 | -0.267 | 0.990 | -0.237 | 1.000 | -0.125 | 1.000 | 1.709 | 0.001 | 0.387 | 0.510 | 0.189 | 0.815 | 0.744 | 0.068 | Conserved hypothetical protein | conserved hypotheticals |
| Rv1716 | Rv1716 | 0.297 | 0.810 | 0.083 | 0.992 | -0.111 | 1.000 | 0.271 | 1.000 | 0.307 | 0.612 | 0.485 | 0.333 | 0.283 | 0.692 | 1.320 | 0.000 | Conserved hypothetical protein | conserved hypotheticals |
| Rv0695 | Rv0695 | -0.065 | 0.971 | 0.161 | 0.990 | 0.370 | 1.000 | 0.244 | 1.000 | -1.201 | 0.001 | 0.493 | 0.252 | -0.524 | 0.278 | 0.120 | 0.800 | Conserved hypothetical protein | conserved hypotheticals |
| Rv2917 | Rv2917 | -0.260 | NA | -1.278 | NA | -0.390 | 1.000 | -0.211 | 1.000 | 0.815 | 0.383 | 0.517 | 0.648 | 1.666 | 0.093 | 0.623 | 0.434 | Conserved hypothetical alanine and arginine rich protein | conserved hypotheticals |
| Rv3031 | Rv3031 | 0.199 | 0.866 | -0.095 | 0.990 | -0.085 | 1.000 | -0.214 | 1.000 | 0.756 | 0.062 | 0.605 | 0.113 | 0.293 | 0.572 | 1.015 | 0.000 | Conserved protein | conserved hypotheticals |
| Rv2522c | Rv2522c | 0.127 | 0.903 | -0.104 | 0.990 | -0.041 | 1.000 | -0.044 | 1.000 | -0.041 | 0.942 | 0.612 | 0.058 | 0.703 | 0.049 | 1.037 | 0.000 | Conserved hypothetical protein | conserved hypotheticals |
| Rv3433c | Rv3433c | 0.490 | 0.694 | 0.058 | 0.992 | -0.169 | 1.000 | -0.199 | 1.000 | 0.584 | 0.299 | 0.615 | 0.195 | 0.031 | 0.978 | 1.266 | 0.000 | Conserved protein | conserved hypotheticals |
| Rv1571 | Rv1571 | -0.215 | NA | 0.103 | NA | 0.300 | 1.000 | -0.023 | 1.000 | 0.088 | 0.946 | 0.638 | 0.411 | 0.216 | 0.838 | 1.017 | 0.033 | Conserved protein | conserved hypotheticals |
| Rv1718 | Rv1718 | -0.473 | 0.661 | 0.182 | 0.990 | 0.026 | 1.000 | 0.361 | 1.000 | 0.377 | 0.551 | 0.662 | 0.205 | 0.032 | 0.981 | 1.076 | 0.008 | Conserved hypothetical protein | conserved hypotheticals |
| Rv2047c | Rv2047c | 0.363 | 0.666 | 0.246 | 0.990 | 0.008 | 1.000 | 0.253 | 1.000 | 0.161 | 0.794 | 0.683 | 0.118 | 0.063 | 0.944 | 1.051 | 0.003 | Conserved hypothetical protein | conserved hypotheticals |
| Rv3510c | Rv3510c | -0.247 | NA | -0.356 | NA | 0.283 | 1.000 | -0.210 | 1.000 | 1.107 | 0.026 | 0.697 | 0.193 | 1.100 | 0.038 | 0.559 | 0.108 | Conserved protein | conserved hypotheticals |
| Rv2974c | Rv2974c | -0.278 | 0.809 | 0.131 | 0.990 | -0.150 | 1.000 | 0.037 | 1.000 | 0.365 | 0.483 | 0.711 | 0.091 | 0.243 | 0.677 | 1.092 | 0.001 | Conserved hypothetical alanine rich protein | conserved hypotheticals |
| Rv2897c | Rv2897c | 0.221 | 0.894 | 0.126 | 0.990 | 0.177 | 1.000 | 0.200 | 1.000 | 0.646 | 0.221 | 0.743 | 0.117 | 0.352 | 0.604 | 1.217 | 0.001 | Conserved hypothetical protein | conserved hypotheticals |
| Rv3422c | Rv3422c | 0.337 | 0.524 | 0.148 | 0.990 | 0.196 | 1.000 | 0.200 | 1.000 | 0.241 | 0.560 | 0.754 | 0.017 | 0.615 | 0.125 | 1.127 | 0.000 | Conserved hypothetical protein | conserved hypotheticals |
| Rv3555c | Rv3555c | 1.088 | 0.014 | 0.572 | 0.672 | 0.583 | 1.000 | 0.118 | 1.000 | 0.364 | 0.477 | 0.772 | 0.044 | 0.710 | 0.106 | 1.146 | 0.000 | Conserved protein | conserved hypotheticals |
| Rv0358 | Rv0358 | 0.118 | NA | -0.234 | NA | -0.388 | 1.000 | 0.221 | 1.000 | 0.387 | 0.612 | 0.826 | 0.174 | 1.046 | 0.090 | 0.836 | 0.062 | Conserved protein | conserved hypotheticals |
| Rv3421c | Rv3421c | 0.223 | 0.871 | 0.032 | 0.992 | -0.021 | 1.000 | -0.255 | 1.000 | 0.198 | 0.788 | 0.872 | 0.066 | -0.087 | 0.929 | 1.067 | 0.006 | Conserved hypothetical protein | conserved hypotheticals |
| Rv2670c | Rv2670c | 0.251 | NA | -0.278 | NA | 0.029 | 1.000 | -0.232 | 1.000 | 0.376 | 0.510 | 0.974 | 0.114 | 1.003 | 0.176 | 1.023 | 0.011 | Conserved hypothetical protein | conserved hypotheticals |
| Rv1685c | Rv1685c | 0.172 | NA | 0.149 | NA | -0.170 | 1.000 | -0.128 | 1.000 | 0.424 | 0.788 | 1.002 | 0.425 | 1.985 | 0.083 | 1.724 | 0.055 | Conserved hypothetical protein | conserved hypotheticals |
| Rv1375 | Rv1375 | -0.668 | 0.483 | -0.667 | 0.898 | -0.243 | 1.000 | -0.854 | 1.000 | -1.762 | 0.001 | 1.119 | 0.059 | 1.443 | 0.016 | 0.662 | 0.234 | Conserved hypothetical protein | conserved hypotheticals |
| Rv3026c | Rv3026c | 0.349 | NA | 0.061 | NA | -0.237 | 1.000 | -0.516 | 1.000 | 0.696 | 0.252 | 1.210 | 0.049 | 0.745 | 0.311 | 0.481 | 0.307 | Conserved hypothetical protein | conserved hypotheticals |
| Rv1913 | Rv1913 | 0.178 | NA | 0.448 | NA | 0.407 | 1.000 | -0.339 | 1.000 | 0.437 | NA | 1.228 | 0.094 | 0.925 | NA | -0.035 | 0.975 | Conserved hypothetical protein | conserved hypotheticals |
| Rv1376 | Rv1376 | 0.029 | 0.992 | -0.389 | 0.990 | -0.122 | 1.000 | -0.598 | 1.000 | -1.147 | 0.034 | 1.269 | 0.018 | 1.240 | 0.034 | 0.645 | 0.215 | Conserved hypothetical protein | conserved hypotheticals |
| Rv2626c | hrp1 | -0.705 | 0.308 | -0.078 | 0.992 | 0.283 | 1.000 | 0.462 | 1.000 | 0.810 | 0.116 | 1.404 | 0.002 | 0.836 | 0.140 | 0.463 | 0.341 | Hypoxic response protein 1 Hrp1 | conserved hypotheticals |
| Rv2628 | Rv2628 | -0.403 | 0.600 | 0.078 | 0.992 | 0.338 | 1.000 | 0.603 | 1.000 | 0.414 | 0.423 | 1.626 | 0.000 | 1.286 | 0.001 | 0.846 | 0.020 | Hypothetical protein | conserved hypotheticals |
| Rv0516c | Rv0516c | -0.632 | 0.345 | -0.299 | 0.990 | -0.094 | 1.000 | -0.446 | 1.000 | -1.657 | 0.000 | -1.252 | 0.003 | -1.192 | 0.009 | -1.716 | 0.000 | Possible anti-anti-sigma factor | information pathways |
| Rv2710 | sigB | -0.094 | 0.954 | 0.177 | 0.990 | 0.047 | 1.000 | 0.113 | 1.000 | -0.122 | 0.848 | -1.107 | 0.005 | -0.656 | 0.192 | -0.654 | 0.088 | RNA polymerase sigma factor SigB | information pathways |
| Rv0717 | rpsN1 | -0.144 | NA | -0.493 | NA | 0.014 | 1.000 | -0.343 | 1.000 | -0.056 | 0.948 | -1.077 | 0.027 | -0.210 | 0.780 | -0.513 | 0.212 | 30S ribosomal protein S14 RpsN1 | information pathways |
| Rv3414c | sigD | -0.420 | 0.244 | -0.139 | 0.990 | 0.011 | 1.000 | -0.050 | 1.000 | -0.258 | 0.478 | -1.004 | 0.000 | -0.714 | 0.005 | -0.084 | 0.773 | Probable alternative RNA polymerase sigma-D factor SigD | information pathways |
| Rv3287c | rsbW | 0.039 | 0.983 | 0.320 | 0.976 | -0.203 | 1.000 | 0.169 | 1.000 | -0.033 | 0.961 | -0.986 | 0.002 | -0.725 | 0.052 | -1.243 | 0.000 | Anti-sigma factor RsbW (sigma negative effector) | information pathways |
| Rv3286c | sigF | 0.207 | 0.807 | 0.484 | 0.735 | 0.424 | 1.000 | 0.041 | 1.000 | -0.156 | 0.762 | -0.955 | 0.001 | -0.786 | 0.019 | -1.309 | 0.000 | Alternative RNA polymerase sigma factor SigF | information pathways |
| Rv3241c | Rv3241c | -0.351 | 0.490 | -0.138 | 0.990 | 0.013 | 1.000 | 0.084 | 1.000 | -0.739 | 0.017 | -0.858 | 0.004 | -0.665 | 0.059 | -1.018 | 0.000 | Conserved protein | information pathways |
| Rv1189 | sigI | 0.260 | 0.778 | 0.022 | NA | 0.857 | 1.000 | 0.068 | 1.000 | -0.220 | 0.711 | -0.746 | 0.123 | 0.000 | 1.000 | -1.550 | 0.000 | Possible alternative RNA polymerase sigma factor SigI | information pathways |
| Rv0001 | dnaA | -0.587 | 0.179 | -0.358 | 0.964 | -0.030 | 1.000 | -0.331 | 1.000 | -0.689 | 0.048 | -0.538 | 0.140 | -0.503 | 0.225 | -1.118 | 0.000 | Chromosomal replication initiator protein DnaA | information pathways |
| Rv2985 | mutT1 | 0.033 | 0.986 | -0.156 | 0.990 | 0.048 | 1.000 | 0.083 | 1.000 | -0.082 | 0.873 | -0.489 | 0.239 | 0.042 | 0.959 | -1.007 | 0.000 | Possible hydrolase MutT1 | information pathways |
| Rv1221 | sigE | 0.105 | 0.968 | 0.277 | 0.990 | -0.016 | 1.000 | 0.206 | 1.000 | -1.019 | 0.066 | -0.462 | 0.488 | 0.125 | 0.904 | -0.217 | 0.735 | Alternative RNA polymerase sigma factor SigE | information pathways |
| Rv2441c | rpmA | -1.029 | 0.002 | -0.477 | 0.735 | 0.218 | 1.000 | -0.122 | 1.000 | -0.836 | 0.012 | -0.187 | 0.673 | 0.027 | 0.973 | -0.575 | 0.051 | 50S ribosomal protein L27 RpmA | information pathways |
| Rv3442c | rpsI | -1.027 | 0.001 | -0.413 | 0.805 | -0.023 | 1.000 | -0.270 | 1.000 | -0.018 | 0.975 | 0.193 | 0.648 | -0.070 | 0.907 | 0.201 | 0.536 | 30S ribosomal protein S9 RpsI | information pathways |
| Rv3585 | radA | 0.133 | 0.910 | 0.136 | 0.990 | 0.043 | 1.000 | 0.010 | 1.000 | -0.164 | 0.764 | 0.534 | 0.176 | 0.005 | 0.998 | 1.147 | 0.000 | DNA repair protein RadA (DNA repair protein SMS) | information pathways |
| Rv2592c | ruvB | -0.017 | 0.999 | 0.009 | 0.998 | -0.064 | 1.000 | 0.048 | 1.000 | 0.867 | 0.133 | 0.632 | 0.188 | 0.287 | 0.677 | 1.103 | 0.004 | Probable holliday junction DNA helicase RuvB | information pathways |
| Rv2191 | Rv2191 | 0.666 | 0.239 | 0.061 | 0.992 | 0.149 | 1.000 | 0.057 | 1.000 | 0.335 | 0.522 | 0.822 | 0.036 | 0.155 | 0.821 | 1.242 | 0.000 | Conserved hypothetical protein | information pathways |
| Rv3201c | Rv3201c | -0.198 | 0.859 | -0.033 | 0.992 | 0.117 | 1.000 | -0.008 | 1.000 | 0.155 | 0.798 | 0.862 | 0.036 | 0.305 | 0.605 | 1.146 | 0.001 | Probable ATP-dependent DNA helicase | information pathways |
| Rv3202c | Rv3202c | -0.008 | 1.000 | 0.062 | 0.992 | 0.118 | 1.000 | 0.015 | 1.000 | 0.416 | 0.359 | 1.055 | 0.003 | 0.365 | 0.482 | 1.345 | 0.000 | Possible ATP-dependent DNA helicase | information pathways |
| Rv3750c | Rv3750c | 0.029 | 0.989 | -0.125 | 0.990 | -0.705 | 1.000 | -0.131 | 1.000 | 0.346 | 0.534 | -1.093 | 0.000 | 0.233 | 0.805 | -0.401 | 0.449 | Possible excisionase | insertion seqs and phages |
| Rv0605 | Rv0605 | 0.502 | 0.263 | 0.318 | 0.965 | 0.123 | 1.000 | 0.217 | 1.000 | -1.055 | 0.001 | -0.669 | 0.054 | -0.818 | 0.021 | 0.185 | 0.614 | Possible resolvase | insertion seqs and phages |
| Rv1042c | Rv1042c | 0.519 | 0.393 | -0.014 | 0.998 | -0.002 | 1.000 | -0.224 | 1.000 | -1.913 | 0.000 | -0.468 | 0.324 | 0.134 | 0.865 | -0.895 | 0.018 | Probable is like-2 transposase | insertion seqs and phages |
| Rv2657c | Rv2657c | -0.797 | NA | -0.189 | NA | 0.029 | 1.000 | -0.325 | 1.000 | -1.447 | 0.066 | -0.392 | 0.542 | 0.185 | 0.849 | -1.022 | 0.027 | Probable PhiRv2 prophage protein | insertion seqs and phages |
| Rv2659c | Rv2659c | -0.272 | 0.886 | 0.012 | 0.998 | -0.851 | 1.000 | -0.401 | 1.000 | 0.233 | 0.784 | -0.336 | 0.646 | 1.685 | 0.068 | 0.078 | 0.935 | Probable PhiRv2 prophage integrase | insertion seqs and phages |
| Rv3467 | Rv3467 | -1.012 | NA | -1.047 | NA | 0.221 | 1.000 | -0.089 | 1.000 | 1.430 | 0.090 | 0.643 | 0.626 | 0.660 | 0.645 | 0.099 | 0.926 | Conserved hypothetical protein | insertion seqs and phages |

| | | | | | | | | | | | | | | | | | | | |
|---|---|---|---|---|---|---|---|---|---|---|---|---|---|---|---|---|---|---|---|
| Rv2812 | *Rv2812* | 1.336 | NA | -0.368 | NA | -0.700 | 1.000 | -0.210 | 1.000 | 0.680 | 0.268 | 1.387 | 0.021 | 1.233 | 0.095 | 0.392 | 0.462 | Probable transposase | insertion seqs and phages |
| Rv0208c | *Rv0208c* | -0.378 | 0.752 | 0.098 | 0.992 | 0.110 | 1.000 | 0.001 | 1.000 | -1.439 | 0.005 | -1.968 | 0.000 | -1.297 | 0.019 | -1.613 | 0.000 | Hypothetical methlytransferase (methylase) | int. metabolism & respiration |
| Rv3083 | *Rv3083* | 1.452 | 0.013 | 0.816 | 0.735 | 1.106 | 1.000 | 1.814 | 0.008 | -1.085 | 0.054 | -1.391 | 0.009 | -0.646 | 0.372 | -1.428 | 0.003 | Probable monooxygenase (hydroxylase) | int. metabolism & respiration |
| Rv1497 | *lipL* | -0.234 | 0.582 | 0.109 | 0.990 | 0.011 | 1.000 | 0.047 | 1.000 | -0.413 | 0.099 | -1.346 | 0.000 | -1.093 | 0.000 | -1.340 | 0.000 | Probable esterase LipL | int. metabolism & respiration |
| Rv3085 | *Rv3085* | 1.006 | 0.085 | 0.807 | 0.672 | 1.233 | 0.257 | 1.889 | 0.001 | -1.606 | 0.001 | -1.286 | 0.007 | -0.310 | 0.692 | -1.445 | 0.001 | Probable short-chain type dehydrogenase/reductase | int. metabolism & respiration |
| Rv3290c | *lat* | -0.307 | 0.599 | 0.152 | 0.990 | -0.042 | 1.000 | 0.193 | 1.000 | -1.377 | 0.000 | -1.255 | 0.000 | -0.640 | 0.073 | -1.693 | 0.000 | Probable L-lysine-epsilon aminotransferase Lat (L-lysine aminotransferase) (lysine 6-aminotransferase) | int. metabolism & respiration |
| Rv2111c | *pup* | -0.097 | 0.951 | 0.215 | 0.990 | -0.252 | 1.000 | 0.335 | 1.000 | -0.837 | 0.092 | -1.233 | 0.005 | -0.711 | 0.224 | -0.754 | 0.086 | Prokaryotic ubiquitin-like protein Pup | int. metabolism & respiration |
| Rv0467 | *icl1* | -1.150 | 0.126 | -0.198 | 0.990 | 0.165 | 1.000 | -0.192 | 1.000 | -1.849 | 0.001 | -1.203 | 0.057 | -1.588 | 0.012 | -1.594 | 0.003 | Isocitrate lyase Icl (isocitrase) (isocitratase) | int. metabolism & respiration |
| Rv3084 | *lipR* | 1.197 | 0.017 | 0.904 | 0.386 | 1.203 | 0.200 | 2.109 | 0.000 | -1.590 | 0.000 | -1.196 | 0.007 | -0.451 | 0.497 | -1.439 | 0.000 | Probable acetyl-hydrolase/esterase LipR | int. metabolism & respiration |
| Rv3049c | *Rv3049c* | -0.258 | 0.771 | -0.008 | 0.998 | -0.065 | 1.000 | -0.038 | 1.000 | -0.551 | 0.193 | -1.187 | 0.001 | -1.041 | 0.007 | -1.229 | 0.000 | Probable monooxygenase | int. metabolism & respiration |
| Rv0770 | *Rv0770* | 0.264 | 0.894 | -0.009 | 0.999 | -0.165 | 1.000 | 0.063 | 1.000 | -0.695 | 0.338 | -1.112 | 0.088 | 0.021 | 0.995 | -0.166 | 0.829 | Probable dehydrogenase/reductase | int. metabolism & respiration |
| Rv3293 | *pcd* | 0.028 | 0.989 | 0.091 | 0.990 | 0.014 | 1.000 | -0.091 | 1.000 | -0.227 | 0.616 | -1.104 | 0.000 | -0.533 | 0.171 | -0.680 | 0.016 | Probable piperideine-6-carboxilic acid dehydrogenase Pcd (piperideine-6-carboxylate dehydrogenase) | int. metabolism & respiration |
| Rv0211 | *pckA* | 0.128 | 0.927 | 0.233 | 0.990 | 0.121 | 1.000 | 0.167 | 1.000 | -0.926 | 0.025 | -0.992 | 0.016 | -0.954 | 0.033 | -1.157 | 0.001 | Probable iron-regulated phosphoenolpyruvate carboxykinase [GTP] PckA (phosphoenolpyruvate carboxylase) (PEPCK)(pep carboxykinase) | int. metabolism & respiration |
| Rv0249c | *Rv0249c* | -0.120 | 0.909 | -0.112 | 0.990 | -0.040 | 1.000 | -0.059 | 1.000 | -1.147 | 0.000 | -0.973 | 0.002 | -0.871 | 0.012 | -1.182 | 0.000 | Probable succinate dehydrogenase [membrane anchor subunit] (succinic dehydrogenase) | int. metabolism & respiration |
| Rv2457c | *clpX* | 0.116 | 0.858 | 0.082 | 0.990 | 0.025 | 1.000 | 0.100 | 1.000 | -0.746 | 0.001 | -0.966 | 0.000 | -0.678 | 0.005 | -1.007 | 0.000 | subunit ClpX | int. metabolism & respiration |
| Rv0693 | *Rv0693* | 0.267 | 0.858 | 0.541 | 0.965 | 0.339 | 1.000 | 0.376 | 1.000 | -2.320 | 0.000 | -0.931 | 0.107 | -0.724 | 0.291 | -1.182 | 0.014 | Probable coenzyme PQQ synthesis protein E PqqE (coenzyme PQQ synthesis protein III) | int. metabolism & respiration |
| Rv3086 | *adhD* | 0.552 | 0.294 | 0.586 | 0.699 | 0.688 | 1.000 | 1.088 | 0.046 | -1.353 | 0.000 | -0.892 | 0.015 | -0.074 | 0.922 | -1.006 | 0.002 | Probable zinc-type alcohol dehydrogenase AdhD (aldehyde reductase) | int. metabolism & respiration |
| Rv1131 | *prpC* | -0.439 | 0.858 | -0.419 | 0.990 | 0.132 | 1.000 | -0.173 | 1.000 | -1.701 | 0.065 | -0.877 | 0.414 | -0.752 | 0.558 | -1.439 | 0.080 | Probable methylcitrate synthase PrpC | int. metabolism & respiration |
| Rv3727 | *Rv3727* | -0.330 | 0.724 | -0.015 | 0.998 | 0.024 | 1.000 | -0.272 | 1.000 | -0.138 | 0.838 | -0.857 | 0.029 | -1.129 | 0.004 | -1.642 | 0.000 | Possible oxidoreductase | int. metabolism & respiration |
| Rv3742c | *Rv3742c* | 0.565 | 0.807 | 0.109 | 0.992 | -0.012 | 1.000 | -0.019 | 1.000 | -1.180 | 0.267 | -0.843 | 0.455 | -0.977 | 0.425 | -1.428 | 0.094 | Possible oxidoreductase | int. metabolism & respiration |
| Rv0560c | *Rv0560c* | -0.705 | 0.525 | -0.044 | 0.992 | 0.122 | 1.000 | 0.100 | 1.000 | -1.750 | 0.004 | -0.822 | 0.239 | -1.236 | 0.090 | -0.981 | 0.102 | Possible benzoquinone methyltransferase (methylase) | int. metabolism & respiration |
| Rv1076 | *lipU* | 0.226 | 0.846 | 0.191 | 0.990 | -0.100 | 1.000 | 0.148 | 1.000 | 0.345 | 0.525 | -0.820 | 0.027 | -0.547 | 0.229 | -1.153 | 0.000 | Possible lipase LipU | int. metabolism & respiration |
| Rv0694 | *Rv0694* | 0.173 | 0.915 | 0.431 | 0.990 | 0.427 | 1.000 | 0.363 | 1.000 | -2.172 | 0.000 | -0.716 | 0.206 | -0.594 | 0.376 | -0.942 | 0.043 | Possible L-lactate dehydrogenase (cytochrome) LldD1 | int. metabolism & respiration |
| Rv0148 | *Rv0148* | -0.581 | 0.344 | -0.008 | 0.998 | 0.072 | 1.000 | -0.041 | 1.000 | -0.915 | 0.029 | -0.643 | 0.157 | -0.781 | 0.102 | -1.005 | 0.006 | Probable short-chain type dehydrogenase/reductase | int. metabolism & respiration |
| Rv2674 | *msrB* | -0.126 | 0.870 | 0.016 | 0.993 | 0.045 | 1.000 | 0.059 | 1.000 | -1.105 | 0.000 | -0.604 | 0.075 | -0.273 | 0.562 | -0.530 | 0.046 | (protein-methionine-R-oxide reductase) (peptide met(O) reductase) | int. metabolism & respiration |
| Rv0886 | *fprB* | -0.766 | 0.136 | -0.284 | 0.990 | 0.210 | 1.000 | -0.427 | 1.000 | -0.773 | 0.080 | -0.543 | 0.239 | -0.855 | 0.083 | -1.128 | 0.002 | (adrenodoxin reductase) (AR) (ferredoxin-NADP(+) reductase) | int. metabolism & respiration |
| Rv0149 | *Rv0149* | -0.016 | 0.992 | 0.083 | 0.992 | 0.046 | 1.000 | -0.026 | 1.000 | -0.892 | 0.004 | -0.530 | 0.123 | -0.917 | 0.005 | -1.150 | 0.000 | Possible quinone oxidoreductase (NADPH:quinone oxidoreductase) (zeta-crystallin) | int. metabolism & respiration |
| Rv1188 | *Rv1188* | 0.845 | 0.179 | 0.510 | 0.964 | 0.192 | 1.000 | 0.113 | 1.000 | -0.385 | 0.543 | -0.528 | 0.392 | -0.582 | 0.359 | -1.684 | 0.000 | Probable proline dehydrogenase | int. metabolism & respiration |
| Rv3854c | *ethA* | 0.746 | 0.041 | 0.409 | 0.839 | 0.121 | 1.000 | 0.195 | 1.000 | -0.395 | 0.305 | -0.494 | 0.166 | -0.462 | 0.252 | -1.191 | 0.000 | Monooxygenase EthA | int. metabolism & respiration |
| Rv2368c | *phoH1* | 1.066 | 0.000 | 0.543 | 0.042 | 0.093 | 1.000 | 0.326 | 1.000 | 0.085 | 0.793 | -0.375 | 0.099 | -0.393 | 0.121 | -0.732 | 0.000 | Probable PHOH-like protein PhoH1 (phosphate starvation-inducible protein PSIH) | int. metabolism & respiration |
| Rv1882c | *Rv1882c* | 1.008 | 0.000 | 0.033 | 0.992 | 0.079 | 1.000 | -0.184 | 1.000 | -0.079 | 0.864 | -0.241 | 0.387 | -0.751 | 0.002 | -1.371 | 0.000 | Probable short-chain type dehydrogenase/reductase | int. metabolism & respiration |
| Rv3324c | *moaC3* | -0.063 | 0.961 | -0.214 | 0.990 | -0.044 | 1.000 | 0.031 | 1.000 | -0.575 | 0.096 | -0.117 | 0.882 | -0.352 | 0.319 | -1.211 | 0.000 | MoaC3 | int. metabolism & respiration |
| Rv1652 | *argC* | -0.039 | 0.983 | -0.502 | 0.735 | 0.117 | 1.000 | -0.122 | 1.000 | 1.070 | 0.001 | -0.073 | 0.888 | -0.210 | 0.698 | 0.577 | 0.074 | ArgC | int. metabolism & respiration |
| Rv3323c | *moaX* | -0.110 | 0.894 | -0.163 | 0.990 | -0.081 | 1.000 | 0.022 | 1.000 | -0.384 | 0.245 | -0.030 | 0.954 | -0.219 | 0.545 | -1.145 | 0.000 | Probable MoaD-MoaE fusion protein MoaX | int. metabolism & respiration |
| Rv0156 | *pntAb* | -0.681 | NA | -0.224 | NA | 0.038 | 1.000 | -0.338 | 1.000 | -0.094 | 0.902 | -0.028 | 0.967 | -0.053 | 0.961 | 1.238 | 0.001 | Probable NAD(P) transhydrogenase (subunit alpha) PntAb [second part; integral membrane protein] (pyridine nucleotide transhydrogenase subunit alpha) (nicotinamide nucleotide transhydrogenase subunit alpha) | int. metabolism & respiration |
| Rv3379c | *dxs2* | 0.412 | NA | 0.747 | NA | 0.756 | 1.000 | 0.057 | 1.000 | 0.478 | 0.351 | 0.047 | 0.955 | 1.621 | 0.008 | 0.414 | 0.344 | deoxyxylulose-5-phosphate synthase) (DXP synthase) (DXPS) | int. metabolism & respiration |
| Rv0327c | *cyp135A1* | 0.613 | NA | -0.064 | NA | 0.363 | 1.000 | 0.561 | 1.000 | 0.473 | 0.528 | 0.093 | 0.919 | 1.487 | 0.059 | 0.344 | 0.592 | Possible cytochrome P450 135A1 Cyp135A1 | int. metabolism & respiration |
| Rv2220 | *glnA1* | -1.046 | 0.000 | -0.440 | 0.641 | -0.065 | 1.000 | -0.363 | 1.000 | 0.002 | 0.995 | 0.149 | 0.680 | 0.160 | 0.698 | -0.325 | 0.200 | Glutamine synthetase GlnA1 (glutamine synthase) (GS-I) | int. metabolism & respiration |
| Rv1655 | *argD* | -0.182 | 0.931 | -0.086 | 0.992 | -0.251 | 1.000 | 0.414 | 1.000 | 0.391 | 0.593 | 0.170 | 0.840 | 0.233 | 0.813 | 1.064 | 0.036 | Probable acetylornithine aminotransferase ArgD | int. metabolism & respiration |

| | | | | | | | | | | | | | | | | | | | |
|---|---|---|---|---|---|---|---|---|---|---|---|---|---|---|---|---|---|---|---|
| **Rv2988c** | *leuC* | 0.013 | 0.999 | 0.198 | 0.990 | 0.034 | 1.000 | 0.283 | 1.000 | 0.265 | 0.699 | 0.183 | 0.792 | 0.316 | 0.670 | **1.091** | **0.011** | LeuC (isopropylmalate isomerase) (alpha-IPM isomerase) (IPMI) | int. metabolism & respiration |
| **Rv3322c** | *Rv3322c* | 0.469 | 0.196 | 0.179 | 0.990 | -0.060 | 1.000 | 0.070 | 1.000 | -0.258 | 0.500 | 0.323 | 0.482 | -0.264 | 0.417 | **-1.059** | **0.000** | Possible methyltransferase | int. metabolism & respiration |
| **Rv0331** | *Rv0331* | -0.260 | 0.877 | -0.048 | 0.992 | 0.303 | 1.000 | 0.221 | 1.000 | 0.075 | 0.938 | 0.347 | 0.622 | **1.349** | **0.020** | **1.123** | **0.014** | Possible dehydrogenase/reductase | int. metabolism & respiration |
| **Rv2987c** | *leuD* | 0.027 | 0.991 | 0.242 | 0.990 | -0.134 | 1.000 | 0.310 | 1.000 | 0.574 | 0.327 | 0.404 | 0.482 | 0.561 | 0.356 | **1.141** | **0.006** | LeuD (isopropylmalate isomerase) (alpha-IPM isomerase) (IPMI) | int. metabolism & respiration |
| **Rv0696** | *Rv0696* | 0.141 | 0.906 | 0.144 | 0.990 | 0.321 | 1.000 | 0.371 | 1.000 | **-1.110** | **0.002** | 0.405 | 0.355 | -0.668 | 0.121 | -0.125 | 0.783 | Probable membrane sugar transferase | int. metabolism & respiration |
| **Rv1714** | *Rv1714* | -0.358 | 0.648 | 0.183 | 0.990 | -0.083 | 1.000 | -0.093 | 1.000 | 0.540 | 0.186 | 0.456 | 0.205 | 0.239 | 0.653 | **1.154** | **0.000** | Probable oxidoreductase | int. metabolism & respiration |
| **Rv0524** | *hemL* | 0.131 | 0.910 | 0.252 | 0.990 | 0.050 | 1.000 | 0.144 | 1.000 | -0.147 | 0.788 | 0.460 | 0.256 | 0.034 | 0.968 | **1.036** | **0.001** | HemL (GSA) (glutamate-1-semialdehyde aminotransferase) (GSA-at) | int. metabolism & respiration |
| **Rv2363** | *amiA2* | -0.129 | NA | -0.896 | NA | 0.021 | 1.000 | -0.158 | 1.000 | 0.481 | 0.417 | 0.460 | 0.510 | **1.286** | **0.046** | 0.473 | 0.337 | Probable amidase AmiA2 (aminohydrolase) | int. metabolism & respiration |
| **Rv3145** | *nuoA* | 0.052 | 0.977 | 0.029 | 0.992 | 0.011 | 1.000 | 0.025 | 1.000 | **-1.205** | **0.000** | 0.463 | 0.242 | 0.206 | 0.707 | 0.766 | 0.008 | Probable NADH dehydrogenase I (chain A) NuoA (NADH-ubiquinone oxidoreductase chain A) | int. metabolism & respiration |
| **Rv0669c** | *Rv0669c* | -0.062 | 0.971 | -0.668 | 0.247 | -0.367 | 1.000 | -0.442 | 1.000 | 0.251 | 0.567 | 0.477 | 0.248 | **1.062** | **0.005** | 0.018 | 0.975 | Possible hydrolase | int. metabolism & respiration |
| **Rv1432** | *Rv1432* | 0.420 | NA | -0.450 | NA | -0.517 | 1.000 | 0.030 | 1.000 | 0.736 | 0.198 | 0.583 | 0.353 | **1.303** | **0.043** | 0.101 | 0.878 | Probable dehydrogenase | int. metabolism & respiration |
| **Rv3419c** | *gcp* | 0.018 | 0.992 | 0.073 | 0.992 | 0.028 | 1.000 | -0.106 | 1.000 | -0.142 | 0.782 | 0.594 | 0.092 | 0.029 | 0.973 | **1.024** | **0.000** | Probable O-sialoglycoprotein endopeptidase Gcp (glycoprotease) | int. metabolism & respiration |
| **Rv1613** | *trpA* | 0.145 | 0.894 | 0.027 | 0.992 | 0.005 | 1.000 | 0.118 | 1.000 | 0.455 | 0.257 | 0.615 | 0.098 | 0.176 | 0.756 | **1.063** | **0.000** | Probable tryptophan synthase, alpha subunit TrpA | int. metabolism & respiration |
| **Rv2958c** | *Rv2958c* | -0.728 | 0.207 | -0.861 | 0.348 | -0.065 | 1.000 | -0.671 | 1.000 | **1.259** | **0.003** | 0.623 | 0.277 | 0.651 | 0.312 | 0.165 | 0.766 | Possible glycosyl transferase | int. metabolism & respiration |
| **Rv0526** | *Rv0526* | 0.289 | 0.778 | 0.128 | 0.990 | -0.007 | 1.000 | 0.055 | 1.000 | 0.112 | 0.868 | 0.639 | 0.169 | 0.104 | 0.900 | **1.035** | **0.005** | Possible thioredoxin protein (thiol-disulfide interchange protein) | int. metabolism & respiration |
| **Rv0032** | *bioF2* | 0.234 | 0.859 | -0.371 | 0.990 | -0.093 | 1.000 | -0.257 | 1.000 | 0.646 | 0.215 | 0.645 | 0.249 | **1.023** | **0.084** | 0.066 | 0.922 | Possible 8-amino-7-oxononanoate synthase BioF2 (AONS) (8-amino-7-ketopelargonate synthase) (7-keto-8-amino-pelargonic acid synthetase) (7-KAP synthetase) (L-alanine--pimelyl CoA ligase) | int. metabolism & respiration |
| **Rv0865** | *mog* | 0.137 | 0.927 | 0.207 | 0.990 | 0.164 | 1.000 | 0.001 | 1.000 | **1.089** | **0.021** | 0.657 | 0.106 | -0.353 | 0.534 | 0.522 | 0.106 | Probable molybdopterin biosynthesis Mog protein | int. metabolism & respiration |
| **Rv3137** | *Rv3137* | -0.636 | 0.073 | -0.459 | 0.735 | -0.184 | 1.000 | -0.340 | 1.000 | **1.253** | **0.000** | 0.670 | 0.082 | **1.491** | **0.000** | 0.946 | 0.001 | Probable monophosphatase | int. metabolism & respiration |
| **Rv2739c** | *Rv2739c* | 0.027 | 0.989 | 0.034 | 0.992 | 0.021 | 1.000 | -0.065 | 1.000 | -0.296 | 0.532 | 0.682 | 0.049 | 0.177 | 0.740 | **1.045** | **0.000** | Possible alanine rich transferase | int. metabolism & respiration |
| **Rv2959c** | *Rv2959c* | -0.774 | 0.161 | **-1.193** | **0.034** | -0.042 | 1.000 | -0.718 | 1.000 | **1.138** | **0.009** | 0.690 | 0.189 | 0.568 | 0.363 | 0.255 | 0.615 | Possible methyltransferase (methylase) | int. metabolism & respiration |
| **Rv3151** | *nuoG* | 0.026 | 0.991 | 0.156 | 0.990 | 0.019 | 1.000 | 0.172 | 1.000 | -0.308 | 0.612 | 0.690 | 0.166 | 0.062 | 0.952 | **1.126** | **0.005** | Probable NADH dehydrogenase I (chain G) NuoG (NADH-ubiquinone oxidoreductase chain G) | int. metabolism & respiration |
| **Rv0186** | *bglS* | 0.053 | 0.972 | 0.067 | 0.992 | 0.009 | 1.000 | -0.043 | 1.000 | 0.174 | 0.709 | 0.691 | 0.037 | 0.214 | 0.668 | **1.015** | **0.000** | Probable beta-glucosidase BglS (gentiobiase) (cellobiase) (beta-D-glucoside glucohydrolase) | int. metabolism & respiration |
| **Rv2678c** | *hemE* | 0.364 | 0.491 | 0.098 | 0.990 | -0.043 | 1.000 | 0.082 | 1.000 | 0.297 | 0.467 | 0.723 | 0.019 | 0.169 | 0.730 | **1.105** | **0.000** | Probable uroporphyrinogen decarboxylase HemE (uroporphyrinogen III decarboxylase) (URO-D) (UPD) | int. metabolism & respiration |
| **Rv0291** | *mycP3* | -0.161 | 0.916 | 0.120 | 0.992 | 0.012 | 1.000 | 0.095 | 1.000 | 0.376 | 0.541 | 0.726 | 0.166 | 0.156 | 0.857 | **1.100** | **0.010** | protease) (subtilisin-like protease) (subtilase-like) (mycosin-3) | int. metabolism & respiration |
| **Rv0161** | *Rv0161* | -0.124 | 0.954 | -0.349 | NA | -0.268 | 1.000 | -0.032 | 1.000 | 0.138 | 0.851 | 0.730 | 0.213 | **1.220** | **0.049** | 0.429 | 0.373 | Possible oxidoreductase | int. metabolism & respiration |
| **Rv0889c** | *citA* | 0.235 | 0.544 | 0.372 | 0.641 | 0.081 | 1.000 | 0.248 | 1.000 | -0.001 | 0.997 | 0.778 | 0.000 | 0.587 | 0.017 | **1.000** | **0.000** | Probable citrate synthase II CitA | int. metabolism & respiration |
| **Rv1569** | *bioF1* | -0.117 | 0.910 | -0.069 | 0.992 | 0.028 | 1.000 | -0.032 | 1.000 | 0.450 | 0.215 | 0.846 | 0.004 | 0.589 | 0.095 | **1.081** | **0.000** | Probable 8-amino-7-oxononanoate synthase BioF1 (AONS) (8-amino-7-ketopelargonate synthase) (7-keto-8-amino-pelargonic acid synthetase) (7-KAP synthetase) (L-alanine--pimelyl CoA ligase) | int. metabolism & respiration |
| **Rv0157** | *pntB* | -0.139 | 0.900 | -0.024 | 0.992 | -0.019 | 1.000 | -0.034 | 1.000 | 0.221 | 0.652 | 0.876 | 0.012 | 0.338 | 0.497 | **1.191** | **0.000** | Probable NAD(P) transhydrogenase (subunit beta) PntB [integral membrane protein] (pyridine nucleotide transhydrogenase subunit beta) (nicotinamide nucleotide transhydrogenase subunit beta) | int. metabolism & respiration |
| **Rv0527** | *ccdA* | 0.165 | 0.894 | 0.142 | 0.990 | -0.001 | 1.000 | 0.169 | 1.000 | -0.187 | 0.760 | 0.978 | 0.017 | 0.143 | 0.849 | **1.203** | **0.001** | Possible cytochrome C-type biogenesis protein CcdA | int. metabolism & respiration |
| **Rv1885c** | *Rv1885c* | 0.048 | 0.983 | -0.824 | NA | -0.493 | 1.000 | -0.716 | 1.000 | **1.462** | **0.002** | **1.001** | **0.043** | **1.103** | **0.031** | 0.925 | 0.024 | Chorismate mutase | int. metabolism & respiration |
| **Rv3303c** | *lpdA* | -0.155 | 0.846 | -0.060 | 0.992 | 0.056 | 1.000 | -0.125 | 1.000 | 0.093 | 0.830 | **1.012** | **0.000** | 0.684 | 0.045 | 0.970 | 0.000 | NAD(P)H quinone reductase LpdA | int. metabolism & respiration |
| **Rv0573c** | *pncB2* | -0.441 | 0.594 | -0.181 | 0.990 | 0.217 | 1.000 | 0.123 | 1.000 | 0.486 | 0.302 | **1.020** | **0.005** | 0.885 | 0.028 | 0.301 | 0.414 | Nicotinic acid phosphoribosyltransferase PncB2 | int. metabolism & respiration |
| **Rv1542c** | *glbN* | -0.171 | 0.915 | -0.319 | 0.990 | 0.308 | 1.000 | -0.797 | 1.000 | -0.017 | 0.985 | 1.057 | 0.139 | **1.338** | **0.038** | 0.172 | 0.875 | Hemoglobin GlbN | int. metabolism & respiration |
| **Rv3548c** | *Rv3548c* | 0.757 | 0.132 | **1.516** | **0.000** | **1.548** | **0.000** | 0.653 | 1.000 | **1.023** | **0.008** | **1.064** | **0.005** | 0.852 | 0.052 | **1.059** | **0.002** | Probable short-chain type dehydrogenase/reductase | int. metabolism & respiration |
| **Rv3559c** | *Rv3559c* | 0.103 | 0.977 | **1.300** | **0.100** | 0.766 | 1.000 | 0.516 | 1.000 | 0.844 | 0.187 | **1.241** | **0.031** | 0.939 | 0.145 | **1.127** | **0.015** | Probable oxidoreductase | int. metabolism & respiration |
| **Rv3109** | *moaA1* | 0.969 | NA | -0.586 | NA | -0.554 | 1.000 | 0.021 | 1.000 | 0.642 | 0.377 | 1.241 | 0.128 | **1.871** | **0.026** | 0.292 | 0.676 | MoaA1 | int. metabolism & respiration |

| | | | | | | | | | | | | | | | | | | | |
|---|---|---|---|---|---|---|---|---|---|---|---|---|---|---|---|---|---|---|---|
| **Rv3553** | *Rv3553* | 0.609 | 0.505 | **1.249** | **0.034** | **1.639** | **0.000** | 0.521 | 1.000 | 0.846 | 0.109 | **1.324** | **0.003** | 0.953 | 0.079 | **1.399** | **0.000** | Possible oxidoreductase | int. metabolism & respiration |
| **Rv0428c** | *Rv0428c* | 0.076 | NA | -0.343 | NA | -0.145 | 1.000 | -0.212 | 1.000 | 0.131 | 0.880 | **1.330** | **0.025** | 0.110 | 0.924 | 0.838 | 0.042 | GCN5-related N-acetyltransferase | int. metabolism & respiration |
| **Rv3549c** | *Rv3549c* | **1.542** | **0.000** | **2.089** | **0.000** | **1.913** | **0.000** | 0.664 | 1.000 | **1.765** | **0.000** | **1.342** | **0.000** | **1.121** | **0.007** | **1.358** | **0.000** | Probable short-chain type dehydrogenase/reductase | int. metabolism & respiration |
| **Rv3552** | *Rv3552* | 0.230 | 0.911 | **1.673** | **0.006** | **2.145** | **0.000** | 0.447 | 1.000 | 0.981 | 0.122 | **1.345** | **0.013** | **1.533** | **0.007** | **1.484** | **0.002** | Possible CoA-transferase (beta subunit) | int. metabolism & respiration |
| **Rv2121c** | *hisG* | 0.553 | NA | -0.578 | NA | 0.039 | 1.000 | -0.374 | 1.000 | 0.938 | NA | **1.360** | **0.027** | 0.360 | NA | 0.239 | 0.693 | ATP phosphoribosyltransferase HisG | int. metabolism & respiration |
| **Rv3551** | *Rv3551* | **1.150** | **0.044** | **2.148** | **0.000** | **2.054** | **0.000** | 0.646 | 1.000 | **1.444** | **0.002** | **1.453** | **0.001** | **1.350** | **0.005** | **1.720** | **0.000** | Possible CoA-transferase (alpha subunit) | int. metabolism & respiration |
| **Rv1050** | *Rv1050* | 0.712 | NA | -0.426 | NA | -0.278 | 1.000 | -0.497 | 1.000 | 1.068 | NA | **1.868** | **0.053** | 1.496 | NA | -0.249 | NA | Probable oxidoreductase | int. metabolism & respiration |
| **Rv2277c** | *Rv2277c* | -0.728 | NA | -1.325 | NA | -0.252 | 1.000 | -0.374 | 1.000 | 0.484 | 0.484 | **2.530** | **0.041** | 0.233 | 0.832 | 0.200 | 0.775 | Possible glycerolphosphodiesterase | int. metabolism & respiration |
| **Rv0468** | *fadB2* | **-1.004** | **0.091** | -0.237 | 0.990 | -0.013 | 1.000 | -0.369 | 1.000 | **-1.346** | **0.006** | **-1.118** | **0.021** | **-1.406** | **0.005** | **-1.430** | **0.001** | 3-hydroxybutyryl-CoA dehydrogenase FadB2 (beta-hydroxybutyryl-CoA dehydrogenase) (BHBD) | lipid metabolism |
| **Rv2482c** | *plsB2* | 0.037 | 0.978 | 0.114 | 0.990 | -0.022 | 1.000 | 0.144 | 1.000 | -0.483 | 0.045 | -0.853 | 0.000 | -0.598 | 0.016 | **-1.076** | **0.000** | (GPAT) | lipid metabolism |
| **Rv3089** | *fadD13* | -0.045 | 0.983 | 0.279 | 0.990 | 0.701 | 1.000 | 0.669 | 1.000 | **-1.571** | **0.000** | -0.842 | 0.029 | -0.610 | 0.193 | **-1.381** | **0.000** | Probable chain -fatty-acid-CoA ligase FadD13 (fatty-acyl-CoA synthetase) | lipid metabolism |
| **Rv3088** | *tgs4* | -0.006 | 1.000 | 0.528 | 0.823 | 0.808 | 1.000 | **1.155** | **0.046** | **-1.287** | **0.000** | -0.810 | 0.042 | -0.318 | 0.577 | **-1.336** | **0.000** | Putative triacylglycerol synthase (diacylglycerol acyltransferase) Tgs4 | lipid metabolism |
| **Rv3523** | *ltp3* | -0.137 | 0.915 | 0.174 | 0.990 | -0.135 | 1.000 | 0.026 | 1.000 | **-1.026** | **0.010** | -0.727 | 0.091 | -0.677 | 0.169 | -0.928 | 0.010 | Ltp3 | lipid metabolism |
| **Rv0129c** | *fbpC* | -0.213 | 0.858 | -0.248 | 0.990 | -0.125 | 1.000 | -0.054 | 1.000 | 0.017 | 0.983 | -0.663 | 0.124 | -0.781 | 0.097 | **-1.043** | **0.003** | Secreted antigen 85-C FbpC (85C) (antigen 85 complex C) (AG58C) (mycolyl transferase 85C) (fibronectin-binding protein C) | lipid metabolism |
| **Rv3087** | *Rv3087* | 0.217 | 0.842 | 0.609 | 0.716 | 0.902 | 0.642 | **1.166** | **0.053** | -0.998 | 0.013 | -0.636 | 0.137 | -0.209 | 0.748 | -0.897 | 0.011 | acyltransferase) | lipid metabolism |
| **Rv0672** | *fadE8* | -0.284 | 0.622 | 0.076 | 0.992 | 0.106 | 1.000 | 0.043 | 1.000 | -0.501 | 0.112 | -0.594 | 0.038 | **-1.049** | **0.000** | -0.333 | 0.209 | Probable acyl-CoA dehydrogenase FadE8 | lipid metabolism |
| **Rv3734c** | *tgs2* | 0.275 | 0.754 | 0.132 | 0.990 | -0.118 | 1.000 | 0.239 | 1.000 | -0.533 | 0.239 | -0.390 | 0.397 | -0.435 | 0.407 | **-1.447** | **0.000** | Putative triacylglycerol synthase (diacylglycerol acyltransferase) Tgs2 | lipid metabolism |
| **Rv2590** | *fadD9* | 0.207 | 0.829 | 0.287 | 0.990 | 0.165 | 1.000 | 0.321 | 1.000 | **-1.221** | **0.000** | -0.341 | 0.462 | -0.631 | 0.161 | -0.902 | 0.007 | Probable fatty-acid-CoA ligase FadD9 (fatty-acid-CoA synthetase) (fatty-acid-CoA synthase) | lipid metabolism |
| **Rv2243** | *fabD* | -0.308 | 0.636 | -0.526 | 0.641 | -0.002 | 1.000 | -0.259 | 1.000 | **1.184** | **0.000** | -0.029 | 0.954 | 0.238 | 0.633 | 0.752 | 0.012 | Meromycolate extension acyl carrier protein AcpM | lipid metabolism |
| **Rv2246** | *kasB* | **-1.200** | **0.000** | -0.455 | 0.823 | -0.088 | 1.000 | -0.046 | 1.000 | 0.538 | 0.162 | -0.015 | 0.977 | 0.094 | 0.883 | **1.030** | **0.001** | Acetyl/propionyl-CoA carboxylase (beta subunit) AccD6 | lipid metabolism |
| **Rv1013** | *pks16* | **-1.039** | **0.000** | -0.471 | 0.166 | -0.006 | 1.000 | -0.166 | 1.000 | **1.336** | **0.000** | 0.044 | 0.888 | 0.178 | 0.604 | 0.033 | 0.912 | Putative polyketide synthase Pks16 | lipid metabolism |
| **Rv1153c** | *omt* | **1.008** | **0.024** | 0.102 | NA | 0.429 | 1.000 | -0.012 | 1.000 | -0.142 | 0.825 | 0.046 | 0.944 | 0.282 | 0.682 | -0.581 | 0.093 | Probable O-methyltransferase Omt | lipid metabolism |
| **Rv2245** | *kasA* | **-1.476** | **0.000** | -0.566 | 0.641 | -0.026 | 1.000 | -0.060 | 1.000 | 0.801 | 0.017 | 0.057 | 0.910 | 0.157 | 0.779 | 0.950 | 0.001 | 3-oxoacyl-[acyl-carrier protein] synthase 2 KasB (beta-ketoacyl-ACP synthase) (KAS I) | lipid metabolism |
| **Rv3824c** | *papA1* | -0.176 | 0.949 | -0.441 | 0.990 | -0.135 | 1.000 | 0.058 | 1.000 | -0.821 | 0.330 | 0.062 | 0.957 | **2.192** | **0.003** | **2.210** | **0.000** | Conserved polyketide synthase associated protein PapA1 | lipid metabolism |
| **Rv2244** | *acpM* | **-1.258** | **0.000** | -0.755 | 0.118 | -0.135 | 1.000 | -0.271 | 1.000 | **1.038** | **0.000** | 0.135 | 0.770 | 0.204 | 0.685 | 0.701 | 0.014 | 3-oxoacyl-[acyl-carrier protein] synthase 1 KasA (beta-ketoacyl-ACP synthase) (KAS I) | lipid metabolism |
| **Rv3820c** | *papA2* | 0.013 | 0.997 | -0.991 | NA | -0.034 | 1.000 | -0.205 | 1.000 | 0.630 | 0.129 | 0.223 | 0.743 | **1.209** | **0.023** | 0.441 | 0.276 | Possible conserved polyketide synthase associated protein PapA2 | lipid metabolism |
| **Rv2523c** | *acpS* | -0.134 | 0.945 | -0.171 | 0.990 | -0.121 | 1.000 | 0.233 | 1.000 | -0.242 | 0.746 | 0.516 | 0.392 | 0.506 | 0.488 | **1.238** | **0.004** | synthase) (CoA:APO-[ACP]pantetheinephosphotransferase) (CoA:APO-[acyl-carrier protein]pantetheinephosphotransferase) | lipid metabolism |
| **Rv2524c** | *fas* | -0.177 | 0.875 | 0.023 | 0.993 | -0.036 | 1.000 | 0.138 | 1.000 | 0.915 | 0.017 | 0.519 | 0.233 | 0.379 | 0.483 | **1.310** | **0.000** | Probable fatty acid synthase Fas (fatty acid synthetase) | lipid metabolism |
| **Rv1181** | *pks4* | -0.210 | 0.906 | -0.091 | 0.992 | 0.052 | 1.000 | -0.133 | 1.000 | -0.226 | 0.788 | 0.526 | 0.459 | **1.693** | **0.003** | **1.390** | **0.006** | Probable polyketide beta-ketoacyl synthase Pks4 | lipid metabolism |
| **Rv2947c** | *pks15* | 0.048 | 0.978 | 0.063 | 0.992 | -0.086 | 1.000 | 0.183 | 1.000 | 0.945 | 0.004 | 0.577 | 0.120 | 0.331 | 0.496 | **1.035** | **0.001** | Probable polyketide synthase Pks15 | lipid metabolism |
| **Rv2934** | *ppsD* | 0.044 | 0.983 | 0.213 | 0.990 | 0.020 | 1.000 | 0.147 | 1.000 | 0.560 | 0.245 | 0.710 | 0.106 | 0.288 | 0.643 | **1.107** | **0.002** | Phenolpthiocerol synthesis type-I polyketide synthase PpsD | lipid metabolism |
| **Rv3556c** | *fadA6* | 0.512 | 0.314 | **1.252** | **0.000** | **1.396** | **0.000** | 0.510 | 1.000 | 0.874 | 0.010 | 0.722 | 0.040 | 0.965 | 0.006 | **1.228** | **0.000** | Probable acetyl-CoA acetyltransferase FadA6 (acetoacetyl-CoA thiolase) | lipid metabolism |
| **Rv1182** | *papA3* | 0.041 | 0.989 | -0.234 | 0.990 | -0.077 | 1.000 | -0.431 | 1.000 | -0.852 | 0.193 | 0.730 | 0.355 | **2.150** | **0.000** | **1.625** | **0.002** | Probable conserved polyketide synthase associated protein PapA3 | lipid metabolism |
| **Rv2933** | *ppsC* | 0.212 | 0.859 | 0.195 | 0.990 | 0.047 | 1.000 | 0.144 | 1.000 | 0.766 | 0.096 | 0.740 | 0.106 | 0.335 | 0.593 | **1.210** | **0.001** | Phenolpthiocerol synthesis type-I polyketide synthase PpsC | lipid metabolism |
| **Rv3561** | *fadD3* | **1.171** | **0.028** | **1.088** | **0.099** | 0.583 | 1.000 | 0.403 | 1.000 | 0.719 | 0.150 | 0.795 | 0.098 | 0.323 | 0.628 | 0.790 | 0.052 | Probable fatty-acid-CoA ligase FadD3 (fatty-acid-CoA synthetase) (fatty-acid-CoA synthase) | lipid metabolism |
| **Rv3563** | *fadE32* | 0.355 | 0.803 | 0.616 | 0.735 | 0.635 | 1.000 | 0.255 | 1.000 | 0.467 | 0.472 | 0.818 | 0.096 | 0.580 | 0.343 | **1.206** | **0.003** | Probable acyl-CoA dehydrogenase FadE32 | lipid metabolism |
| **Rv2946c** | *pks1* | 0.119 | 0.940 | 0.116 | 0.992 | -0.032 | 1.000 | 0.098 | 1.000 | **1.010** | **0.023** | 0.924 | 0.045 | 0.424 | 0.497 | **1.378** | **0.000** | Probable polyketide synthase Pks1 | lipid metabolism |
| **Rv2383c** | *mbtB* | -0.386 | 0.636 | -0.196 | 0.990 | 0.097 | 1.000 | -0.183 | 1.000 | 0.752 | 0.081 | **1.009** | **0.013** | **1.169** | **0.009** | 0.834 | 0.030 | synthetase) | lipid metabolism |
| **Rv2378c** | *mbtG* | -0.007 | 1.000 | 0.315 | 0.964 | 0.013 | 1.000 | 0.009 | 1.000 | 0.575 | 0.146 | **1.073** | **0.000** | 0.500 | 0.302 | 0.764 | 0.008 | Lysine-N-oxygenase MbtG (L-lysine 6-monooxygenase) (lysine N6-hydroxylase) | lipid metabolism |

| | | | | | | | | | | | | | | | | | | | |
|---|---|---|---|---|---|---|---|---|---|---|---|---|---|---|---|---|---|---|---|
| Rv0119 | *fadD7* | -0.472 | NA | -0.111 | NA | -0.614 | 1.000 | -0.095 | 1.000 | 0.467 | 0.509 | 1.090 | 0.081 | 0.844 | 0.297 | 0.421 | 0.472 | Probable fatty-acid-CoA ligase FadD7 (fatty-acid-CoA synthetase) (fatty-acid-CoA synthase) | lipid metabolism |
| Rv2382c | *mbtC* | 0.179 | 0.915 | 0.122 | 0.992 | -0.185 | 1.000 | 0.264 | 1.000 | 0.960 | 0.066 | 1.217 | 0.017 | 1.166 | 0.077 | 0.910 | 0.061 | Polyketide synthetase MbtC (polyketide synthase) | lipid metabolism |
| Rv3560c | *fadE30* | 0.487 | 0.600 | 1.530 | 0.000 | 0.992 | 0.382 | 0.429 | 1.000 | 1.090 | 0.020 | 1.248 | 0.003 | 0.529 | 0.350 | 1.042 | 0.006 | Probable acyl-CoA dehydrogenase FadE30 | lipid metabolism |
| Rv3550 | *echA20* | 0.784 | 0.312 | 2.227 | 0.000 | 2.093 | 0.000 | 0.771 | 1.000 | 1.154 | 0.020 | 1.885 | 0.000 | 1.192 | 0.014 | 1.566 | 0.000 | Probable enoyl-CoA hydratase EchA20 (enoyl hydrase) (unsaturated acyl-CoA hydratase) (crotonase) | lipid metabolism |
| Rv0280 | *PPE3* | 0.043 | 0.978 | 0.102 | 0.990 | 0.002 | 1.000 | 0.098 | 1.000 | -0.861 | 0.003 | -1.161 | 0.000 | -1.187 | 0.000 | -1.098 | 0.000 | PPE family protein PPE3 | PE/PPE |
| Rv2352c | *PPE38* | -0.548 | 0.305 | -0.346 | 0.976 | 0.512 | 1.000 | -1.391 | 1.000 | -0.729 | 0.057 | -1.131 | 0.001 | 0.901 | 0.057 | 0.000 | 1.000 | PPE family protein PPE38 | PE/PPE |
| Rv0442c | *PPE10* | 0.420 | 0.511 | 0.276 | 0.990 | -0.015 | 1.000 | 0.277 | 1.000 | -0.032 | 0.965 | -1.050 | 0.003 | -1.173 | 0.001 | -1.516 | 0.000 | PPE family protein PPE10 | PE/PPE |
| Rv3558 | *PPE64* | -0.474 | 0.435 | 0.071 | 0.992 | -0.039 | 1.000 | 0.022 | 1.000 | 0.004 | 0.994 | -1.037 | 0.003 | -1.013 | 0.009 | -0.911 | 0.006 | PPE family protein PPE64 | PE/PPE |
| Rv1646 | *PE17* | -0.263 | 0.747 | 0.112 | 0.990 | 0.041 | 1.000 | -0.026 | 1.000 | -0.776 | 0.036 | -0.669 | 0.075 | -0.694 | 0.093 | -1.377 | 0.000 | PE family protein PE17 | PE/PPE |
| Rv1809 | *PPE33* | -0.620 | 0.421 | -0.275 | 0.990 | -0.262 | 1.000 | 0.042 | 1.000 | -0.864 | 0.091 | -0.623 | 0.252 | -0.747 | 0.216 | -1.634 | 0.000 | PPE family protein PPE33 | PE/PPE |
| Rv0151c | *PE1* | 0.261 | 0.807 | 0.239 | 0.990 | 0.238 | 1.000 | 0.029 | 1.000 | 0.355 | 0.536 | -0.577 | 0.251 | -0.291 | 0.687 | -1.040 | 0.008 | PE family protein PE1 | PE/PPE |
| Rv0304c | *PPE5* | 0.252 | 0.807 | 0.123 | 0.990 | 0.031 | 1.000 | 0.149 | 1.000 | -0.893 | 0.026 | -0.505 | 0.286 | -0.323 | 0.585 | -1.078 | 0.002 | PPE family protein PPE5 | PE/PPE |
| Rv0160c | *PE4* | 0.551 | NA | -0.409 | NA | 0.291 | 1.000 | -0.204 | 1.000 | 0.438 | 0.557 | -0.305 | 0.709 | -0.288 | 0.774 | -1.291 | 0.013 | PE family protein PE4 | PE/PPE |
| Rv1808 | *PPE32* | -0.197 | 0.894 | -0.617 | 0.870 | -0.439 | 1.000 | -0.267 | 1.000 | -0.919 | 0.075 | -0.157 | 0.827 | -0.358 | 0.625 | -1.010 | 0.024 | PPE family protein PPE32 | PE/PPE |
| Rv3746c | *PE34* | 0.015 | 0.999 | 0.474 | 0.990 | -0.039 | 1.000 | 0.443 | 1.000 | -1.567 | 0.005 | 0.001 | 1.000 | 0.573 | 0.483 | -0.253 | 0.713 | protein) | PE/PPE |
| Rv1917c | *PPE34* | 0.363 | 0.724 | -0.293 | 0.990 | 0.030 | 1.000 | 0.059 | 1.000 | 0.560 | 0.280 | 0.146 | 0.836 | 1.239 | 0.020 | 0.086 | 0.888 | PPE family protein PPE34 | PE/PPE |
| Rv1168c | *PPE17* | -0.400 | 0.750 | -0.132 | 0.992 | 0.174 | 1.000 | -0.028 | 1.000 | -1.448 | 0.006 | 0.180 | 0.816 | 0.284 | 0.731 | 0.075 | 0.922 | PPE family protein PPE17 | PE/PPE |
| Rv1169c | *lipX* | -0.502 | 0.750 | -0.100 | 0.992 | 0.016 | 1.000 | 0.570 | 1.000 | -1.438 | 0.034 | 0.200 | 0.834 | 0.253 | 0.819 | 0.218 | 0.780 | PE family protein. Possible lipase LipX. | PE/PPE |
| Rv0285 | *PE5* | -0.243 | 0.843 | 0.209 | 0.990 | -0.053 | 1.000 | 0.229 | 1.000 | 0.197 | 0.769 | 0.432 | 0.442 | 0.251 | 0.731 | 1.006 | 0.014 | PE family protein PE5 | PE/PPE |
| Rv2353c | *PPE39* | 0.286 | 0.774 | 0.045 | 0.992 | 0.158 | 1.000 | 0.233 | 1.000 | 0.157 | 0.803 | 0.456 | 0.506 | -0.381 | 0.523 | -1.093 | 0.003 | PPE family protein PPE39 | PE/PPE |
| Rv1430 | *PE16* | 0.122 | 0.914 | -0.747 | 0.641 | -0.182 | 1.000 | -0.449 | 1.000 | 0.051 | 0.929 | 0.521 | 0.303 | 1.098 | 0.010 | -0.065 | 0.910 | PE family protein PE16 | PE/PPE |
| Rv3135 | *PPE50* | 0.000 | 1.000 | -0.499 | 0.965 | 0.306 | 1.000 | -0.193 | 1.000 | 0.000 | 1.000 | 0.534 | 0.407 | 0.821 | 0.250 | 1.314 | 0.008 | PPE family protein PPE50 | PE/PPE |
| Rv1325c | *PE_PGRS24* | -0.476 | 0.179 | -0.031 | 0.992 | -0.027 | 1.000 | 0.210 | 1.000 | -0.076 | 0.853 | 0.608 | 0.020 | 0.461 | 0.140 | 1.091 | 0.000 | PE-PGRS family protein PE_PGRS24 | PE/PPE |
| Rv1068c | *PE_PGRS20* | 0.107 | 0.975 | -0.135 | 0.990 | -0.010 | 1.000 | 0.092 | 1.000 | 0.379 | 0.601 | 0.659 | 0.198 | 0.708 | 0.233 | 1.021 | 0.008 | PE-PGRS family protein PE_PGRS20 | PE/PPE |
| Rv1067c | *PE_PGRS19* | -0.703 | 0.180 | 0.024 | 0.992 | -0.028 | 1.000 | 0.083 | 1.000 | 0.245 | 0.663 | 0.720 | 0.061 | 0.374 | 0.487 | 1.048 | 0.001 | PE-PGRS family protein PE_PGRS19 | PE/PPE |
| Rv3136 | *PPE51* | -1.320 | 0.000 | -0.475 | 0.437 | -0.068 | 1.000 | -0.535 | 1.000 | 1.071 | 0.000 | 1.164 | 0.000 | 1.771 | 0.000 | 1.658 | 0.000 | PPE family protein PPE51 | PE/PPE |
| Rv0981 | *mprA* | 0.220 | 0.805 | 0.278 | 0.990 | 0.061 | 1.000 | -0.033 | 1.000 | -1.364 | 0.000 | -1.455 | 0.000 | -1.115 | 0.002 | -1.779 | 0.000 | Mycobacterial persistence regulator MRPA (two component response transcriptional regulatory protein) | regulatory proteins |
| Rv3095 | *Rv3095* | 0.439 | 0.320 | 0.045 | 0.992 | 0.149 | 1.000 | 0.104 | 1.000 | -0.790 | 0.010 | -1.190 | 0.000 | -0.498 | 0.176 | -1.093 | 0.000 | Hypothetical transcriptional regulatory protein | regulatory proteins |
| Rv2034 | *Rv2034* | -0.916 | NA | -0.374 | NA | -0.050 | 1.000 | 0.368 | 1.000 | -0.895 | 0.339 | -1.173 | 0.097 | -0.298 | 0.793 | -0.112 | 0.903 | ArsR repressor protein | regulatory proteins |
| Rv1129c | *Rv1129c* | 0.217 | 0.946 | 0.087 | 0.992 | 0.180 | 1.000 | 0.260 | 1.000 | -1.494 | 0.100 | -1.102 | 0.252 | -1.530 | 0.125 | -0.708 | 0.424 | Probable transcriptional regulator protein | regulatory proteins |
| Rv0144 | *Rv0144* | 0.075 | 0.971 | 0.086 | 0.992 | 0.075 | 1.000 | 0.175 | 1.000 | -1.630 | 0.000 | -0.912 | 0.054 | -0.680 | 0.224 | -1.089 | 0.007 | Probable transcriptional regulatory protein (possibly TetR-family) | regulatory proteins |
| Rv1534 | *Rv1534* | 0.302 | 0.738 | 0.121 | 0.990 | 0.014 | 1.000 | 0.051 | 1.000 | -1.312 | 0.001 | -0.864 | 0.039 | -0.594 | 0.234 | -0.881 | 0.015 | Probable transcriptional regulator | regulatory proteins |
| Rv1556 | *Rv1556* | -0.037 | 0.986 | -0.364 | 0.990 | -0.114 | 1.000 | -0.101 | 1.000 | 0.052 | 0.947 | -0.824 | 0.069 | -0.729 | 0.164 | -1.157 | 0.002 | Possible regulatory protein | regulatory proteins |
| Rv0165c | *mce1R* | 0.187 | 0.830 | -0.016 | 0.994 | 0.225 | 1.000 | -0.117 | 1.000 | -0.761 | 0.048 | -0.617 | 0.091 | -0.082 | 0.931 | -1.068 | 0.003 | Probable transcriptional regulatory protein Mce1R (probably GntR-family) | regulatory proteins |
| Rv3219 | *whiB1* | -0.033 | 0.983 | -0.030 | 0.992 | -0.016 | 1.000 | -0.137 | 1.000 | -0.973 | 0.003 | -0.555 | 0.133 | -0.667 | 0.092 | -1.192 | 0.000 | Transcriptional regulatory protein WhiB-like WhiB1. Contains [4FE-4S]2+ cluster. | regulatory proteins |
| Rv3855 | *ethR* | 0.886 | 0.013 | 0.338 | 0.965 | 0.079 | 1.000 | 0.316 | 1.000 | -0.131 | 0.803 | -0.434 | 0.243 | -0.507 | 0.218 | -1.055 | 0.000 | EthR | regulatory proteins |
| Rv3833 | *Rv3833* | -0.224 | 0.909 | 0.111 | 0.992 | -0.264 | 1.000 | 0.018 | 1.000 | -1.276 | 0.020 | -0.079 | 0.914 | -0.120 | 0.896 | -0.690 | 0.129 | Transcriptional regulatory protein (probably AraC-family) | regulatory proteins |
| Rv3143 | *Rv3143* | -0.286 | 0.937 | -0.077 | 0.992 | -0.141 | 1.000 | 0.228 | 1.000 | 0.197 | 0.871 | -0.047 | 0.972 | -1.072 | 0.033 | -1.645 | 0.000 | Probable response regulator | regulatory proteins |
| Rv1776c | *Rv1776c* | 0.288 | NA | 0.247 | NA | 0.089 | 1.000 | 0.262 | 1.000 | 1.116 | 0.049 | -0.036 | 0.957 | 0.409 | 0.598 | 0.022 | 0.981 | Possible transcriptional regulatory protein | regulatory proteins |
| Rv2989 | *Rv2989* | -0.005 | 1.000 | 0.227 | 0.990 | -0.040 | 1.000 | 0.301 | 1.000 | 0.283 | 0.735 | 0.056 | 0.945 | 0.331 | 0.698 | 1.019 | 0.035 | Probable transcriptional regulatory protein | regulatory proteins |
| Rv2359 | *zur* | 1.160 | 0.000 | 0.387 | 0.856 | 0.142 | 1.000 | 0.263 | 1.000 | 0.997 | 0.004 | 0.083 | 0.876 | 0.050 | 0.951 | 0.456 | 0.164 | Probable zinc uptake regulation protein Zur | regulatory proteins |
| Rv3557c | *Rv3557c* | 1.361 | 0.000 | 1.606 | 0.000 | 1.585 | 0.000 | 0.555 | 1.000 | 1.204 | 0.000 | 0.641 | 0.081 | 0.851 | 0.024 | 0.853 | 0.005 | Transcriptional regulatory protein (probably TetR-family) | regulatory proteins |
| Rv2231c | *cobC* | -0.109 | 0.929 | -0.194 | 0.990 | 0.007 | 1.000 | -0.144 | 1.000 | 0.476 | 0.175 | 0.831 | 0.007 | 0.391 | 0.350 | 1.107 | 0.000 | PtkA | regulatory proteins |
| RVnc0015 | *mcr15* | -0.285 | 0.778 | -0.002 | 0.999 | -0.057 | 1.000 | -0.078 | 1.000 | -0.076 | 0.914 | -0.356 | 0.614 | 0.025 | 0.988 | -1.085 | 0.009 | | Stable rnas |
| Rvnr01 | *rrs* | -0.637 | 0.426 | -0.014 | 0.998 | -0.142 | 1.000 | -0.357 | 1.000 | -0.207 | 0.787 | 0.190 | 0.798 | -0.258 | 0.756 | 1.203 | 0.008 | | Stable rnas |
| Rvnr02 | *rrl* | -0.697 | 0.373 | -0.027 | 0.994 | -0.146 | 1.000 | -0.419 | 1.000 | -0.215 | 0.779 | 0.196 | 0.793 | -0.315 | 0.697 | 1.152 | 0.012 | | Stable rnas |
| RVnc0013 | *mcr11* | -0.126 | 0.914 | -0.007 | 0.998 | -0.246 | 1.000 | -0.174 | 1.000 | 0.734 | 0.046 | 0.225 | 0.648 | 0.939 | 0.014 | 1.206 | 0.000 | | Stable rnas |
| RVnc0036a | *MTS2823* | -0.415 | 0.824 | 0.130 | 0.992 | -0.299 | 1.000 | -0.011 | 1.000 | 0.609 | 0.509 | 1.739 | 0.017 | 0.753 | 0.445 | 1.465 | 0.025 | | Stable rnas |
| Rv2348c | *Rv2348c* | -0.527 | 0.373 | -0.062 | 0.992 | -0.238 | 1.000 | 0.121 | 1.000 | -0.311 | 0.531 | -1.126 | 0.002 | -0.829 | 0.057 | -1.378 | 0.000 | Hypothetical protein | unknown |

| | | | | | | | | | | | | | | | | | | | |
|---|---|---|---|---|---|---|---|---|---|---|---|---|---|---|---|---|---|---|---|
| Rv0251c | *hsp* | -0.533 | 0.551 | -0.279 | 0.990 | 0.071 | 1.000 | 0.031 | 1.000 | -0.941 | 0.072 | -1.984 | 0.000 | -1.897 | 0.000 | -1.785 | 0.000 | Heat shock protein Hsp (heat-stress-induced ribosome-binding protein A) | virulence, detoxification, adaptation |
| Rv0960 | *vapC9* | -0.068 | 0.983 | 0.296 | 0.990 | 0.160 | 1.000 | 0.500 | 1.000 | -0.577 | 0.435 | -1.677 | 0.003 | -1.073 | 0.124 | -1.449 | 0.006 | Possible toxin VapC9 | virulence, detoxification, adaptation |
| Rv1720c | *vapC12* | 0.333 | NA | 0.127 | NA | -0.176 | 1.000 | 0.100 | 1.000 | -0.525 | 0.459 | -1.474 | 0.021 | -0.648 | 0.474 | -0.637 | 0.278 | Possible toxin VapC12 | virulence, detoxification, adaptation |
| Rv2595 | *vapB40* | 0.344 | NA | 0.450 | NA | 0.256 | 1.000 | -0.207 | 1.000 | -0.958 | 0.045 | -1.428 | 0.001 | -0.485 | 0.482 | -0.946 | 0.028 | Possible antitoxin VapB40 | virulence, detoxification, adaptation |
| Rv0959A | *vapB9* | -0.277 | NA | -0.517 | NA | 1.000 | 1.000 | -0.397 | 1.000 | -0.292 | NA | -1.402 | 0.093 | -1.749 | NA | -1.845 | NA | Possible antitoxin VapB9 | virulence, detoxification, adaptation |
| Rv0262c | *aac* | -0.153 | 0.900 | 0.020 | 0.994 | 0.110 | 1.000 | 0.054 | 1.000 | -1.239 | 0.003 | -1.387 | 0.000 | -1.835 | 0.000 | -2.526 | 0.000 | Aminoglycoside 2'-N-acetyltransferase Aac (Aac(2')-IC) | virulence, detoxification, adaptation |
| Rv2319c | *Rv2319c* | 0.063 | 0.978 | -0.113 | 0.992 | -0.275 | 1.000 | 0.052 | 1.000 | 0.019 | 0.981 | -1.337 | 0.006 | -0.518 | 0.437 | -0.916 | 0.033 | Universal stress protein family protein | virulence, detoxification, adaptation |
| Rv1971 | *mce3F* | -0.038 | NA | -0.193 | NA | -0.175 | 1.000 | 0.324 | 1.000 | 0.052 | 0.949 | -1.323 | 0.007 | -0.304 | 0.701 | -1.105 | 0.006 | Mce-family protein Mce3F | virulence, detoxification, adaptation |
| Rv0549c | *vapC3* | 0.929 | NA | 0.091 | NA | -0.396 | 1.000 | 0.159 | 1.000 | -0.393 | 0.567 | -1.106 | 0.047 | -0.472 | 0.588 | -0.226 | 0.727 | Possible toxin VapC3 | virulence, detoxification, adaptation |
| Rv2865 | *relF* | 0.702 | NA | 0.603 | NA | -0.168 | 1.000 | 1.119 | 1.000 | -0.041 | 0.967 | -1.011 | 0.074 | -0.089 | 0.939 | -0.387 | 0.495 | Antitoxin RelF | virulence, detoxification, adaptation |
| Rv0277c | *vapC25* | -0.020 | 0.995 | -0.240 | 0.990 | -0.288 | 1.000 | 0.012 | 1.000 | -0.151 | 0.843 | -1.007 | 0.028 | -0.861 | 0.116 | -1.385 | 0.000 | Possible toxin VapC25. Contains PIN domain. | virulence, detoxification, adaptation |
| Rv1398c | *vapB10* | -0.158 | 0.714 | 0.080 | 0.990 | -0.066 | 1.000 | 0.147 | 1.000 | -1.249 | 0.000 | -0.875 | 0.000 | -0.453 | 0.039 | -1.158 | 0.000 | Possible antitoxin VapB10 | virulence, detoxification, adaptation |
| Rv1839c | *vapB13* | -0.251 | 0.707 | -0.144 | 0.990 | -0.062 | 1.000 | 0.398 | 1.000 | -0.983 | 0.005 | -0.871 | 0.053 | -0.607 | 0.242 | -1.028 | 0.005 | Possible antitoxin VapB13 | virulence, detoxification, adaptation |
| Rv0353 | *hspR* | -0.104 | 0.934 | 0.214 | 0.990 | 0.096 | 1.000 | 0.138 | 1.000 | -1.319 | 0.000 | -0.843 | 0.013 | -0.783 | 0.037 | -0.433 | 0.180 | Probable heat shock protein transcriptional repressor HspR (MerR family) | virulence, detoxification, adaptation |
| Rv0352 | *dnaJ1* | -0.167 | 0.882 | 0.161 | 0.990 | 0.085 | 1.000 | 0.133 | 1.000 | -1.040 | 0.004 | -0.759 | 0.052 | -0.547 | 0.238 | -0.233 | 0.573 | Probable chaperone protein DnaJ1 | virulence, detoxification, adaptation |
| Rv2104c | *vapB37* | 0.065 | 0.961 | 0.280 | 0.961 | -0.188 | 1.000 | 0.208 | 1.000 | -1.037 | 0.002 | -0.667 | 0.018 | -0.609 | 0.098 | -0.718 | 0.007 | Possible antitoxin VapB37 | virulence, detoxification, adaptation |
| Rv1636 | *TB15.3* | 0.336 | 0.624 | 0.194 | 0.990 | 0.038 | 1.000 | 0.031 | 1.000 | -0.101 | 0.862 | -0.528 | 0.191 | -0.505 | 0.304 | -1.114 | 0.001 | TB15.3 | virulence, detoxification, adaptation |
| Rv0599c | *vapB27* | 0.156 | 0.845 | 0.263 | 0.976 | -0.074 | 1.000 | 0.233 | 1.000 | -1.171 | 0.008 | -0.515 | 0.168 | 0.072 | 0.927 | -0.720 | 0.039 | Possible antitoxin VapB27 | virulence, detoxification, adaptation |
| Rv0384c | *clpB* | -0.503 | 0.696 | -0.112 | 0.992 | -0.104 | 1.000 | -0.076 | 1.000 | -1.627 | 0.006 | -0.429 | 0.594 | -0.011 | 0.998 | -0.099 | 0.907 | Probable endopeptidase ATP binding protein (chain B) ClpB (ClpB protein) (heat shock protein F84.1) | virulence, detoxification, adaptation |
| Rv3697A | *vapB48* | 0.054 | 0.981 | 0.341 | 0.990 | -0.114 | 1.000 | 0.316 | 1.000 | -1.159 | 0.008 | -0.419 | 0.504 | -0.232 | 0.765 | -0.713 | 0.107 | Possible antitoxin VapB48 | virulence, detoxification, adaptation |
| Rv0064A | *vapB1* | 0.055 | 0.972 | -0.018 | 0.998 | -0.161 | 1.000 | -0.020 | 1.000 | 0.385 | 0.379 | -0.395 | 0.585 | -0.296 | 0.667 | -1.049 | 0.006 | Possible antitoxin VapB1 | virulence, detoxification, adaptation |
| Rv3321c | *vapB44* | 0.230 | 0.778 | 0.114 | 0.990 | 0.235 | 1.000 | -0.021 | 1.000 | -0.047 | 0.956 | -0.393 | 0.438 | -0.618 | 0.206 | -1.362 | 0.000 | Possible antitoxin VapB44 | virulence, detoxification, adaptation |
| Rv3320c | *vapC44* | 0.863 | 0.025 | 0.362 | 0.868 | -0.025 | 1.000 | 0.217 | 1.000 | -0.098 | 0.884 | -0.363 | 0.463 | -0.371 | 0.485 | -1.188 | 0.000 | Possible toxin VapC44. Contains PIN domain. | virulence, detoxification, adaptation |
| Rv2547 | *vapB19* | 0.423 | NA | 0.459 | NA | 0.180 | 1.000 | 0.305 | 1.000 | -1.237 | 0.030 | -0.339 | 0.638 | 0.317 | 0.721 | -1.011 | 0.040 | Possible antitoxin VapB19 | virulence, detoxification, adaptation |
| Rv1740 | *vapB34* | -0.330 | NA | 0.088 | NA | -0.158 | 1.000 | 0.218 | 1.000 | -1.047 | NA | -0.257 | 0.791 | -0.447 | NA | -1.143 | 0.056 | Possible antitoxin VapB34 | virulence, detoxification, adaptation |
| Rv3385c | *vapB46* | -0.122 | NA | -0.047 | NA | 0.461 | 1.000 | -0.288 | 1.000 | -1.232 | 0.081 | -0.213 | 0.737 | -0.120 | 0.906 | 0.838 | 0.085 | Possible antitoxin VapB46 | virulence, detoxification, adaptation |
| Rv2527 | *vapC17* | -0.229 | 0.808 | 0.025 | 0.993 | 0.046 | 1.000 | 0.341 | 1.000 | -0.550 | 0.310 | 0.106 | 0.853 | -0.603 | 0.085 | -1.213 | 0.000 | Possible toxin VapC17 | virulence, detoxification, adaptation |
| Rv1965 | *yrbE3B* | 0.562 | NA | -0.662 | NA | 0.271 | 1.000 | 0.046 | 1.000 | 0.239 | 0.803 | 0.216 | 0.834 | 1.383 | 0.098 | 0.362 | 0.632 | Conserved hypothetical integral membrane protein YrbE3B | virulence, detoxification, adaptation |
| Rv2429 | *ahpD* | -0.831 | NA | -0.409 | NA | -0.272 | 1.000 | -0.417 | 1.000 | 0.103 | 0.932 | 0.240 | 0.799 | -0.368 | 0.698 | -1.128 | 0.050 | Alkyl hydroperoxide reductase D protein AhpD (alkyl hydroperoxidase D) | virulence, detoxification, adaptation |
| Rv3269 | *Rv3269* | 1.464 | 0.038 | 0.660 | 0.964 | 0.297 | 1.000 | 0.427 | 1.000 | 1.384 | 0.031 | 0.858 | 0.220 | 0.301 | 0.771 | 1.049 | 0.070 | Conserved protein | virulence, detoxification, adaptation |
| Rv2230c | *Rv2230c* | -0.228 | 0.827 | -0.101 | 0.990 | -0.083 | 1.000 | -0.083 | 1.000 | -0.042 | 0.945 | 0.881 | 0.010 | 0.389 | 0.404 | 1.031 | 0.000 | Possible toxin VapC16 | virulence, detoxification, adaptation |
| Rv2623 | *TB31.7* | -0.176 | 0.875 | 0.134 | 0.990 | 0.311 | 1.000 | 0.537 | 1.000 | 0.688 | 0.090 | 1.006 | 0.007 | 0.735 | 0.098 | 0.839 | 0.014 | Universal stress protein family protein TB31.7 | virulence, detoxification, adaptation |
| Rv2190c | *Rv2190c* | 0.070 | 0.968 | -0.402 | 0.835 | -0.108 | 1.000 | -0.286 | 1.000 | 1.139 | 0.000 | 1.048 | 0.000 | 0.435 | 0.278 | 0.947 | 0.000 | Conserved hypothetical protein | virulence, detoxification, adaptation |
| Rv2031c | *hspX* | -0.388 | 0.622 | 0.094 | 0.992 | 0.157 | 1.000 | 0.539 | 1.000 | 0.541 | 0.250 | 1.176 | 0.003 | 1.109 | 0.008 | 0.648 | 0.087 | Heat shock protein HspX (alpha-crystallin homolog) (14 kDa antigen) (HSP16.3) | virulence, detoxification, adaptation |
| Rv2624c | *Rv2624c* | 0.043 | 0.988 | -0.106 | 0.992 | 0.180 | 1.000 | 0.237 | 1.000 | 1.041 | 0.049 | 1.215 | 0.020 | 0.992 | 0.097 | 0.709 | 0.152 | Universal stress protein family protein | virulence, detoxification, adaptation |
| Rv3473c | *bpoA* | 0.074 | 0.969 | -2.069 | 0.598 | -0.073 | 1.000 | -0.282 | 1.000 | 0.074 | 0.911 | 1.679 | 0.044 | -0.148 | 0.849 | -0.265 | 0.532 | Possible peroxidase BpoA (non-haem peroxidase) | virulence, detoxification, adaptation |
| Rv0063a | *Rv0063a* | 0.508 | NA | -0.356 | NA | -0.063 | 1.000 | -0.020 | 1.000 | 0.329 | NA | -1.905 | 0.068 | -0.843 | NA | -0.448 | NA | | |
| RV1984a | *RV1984a* | 0.613 | NA | 0.532 | NA | -0.463 | 1.000 | -0.351 | 1.000 | -0.530 | NA | -1.367 | 0.037 | -0.757 | NA | -1.024 | 0.085 | | |
| Rv2386a | *Rv2386a* | -0.361 | 0.713 | -0.223 | 0.990 | 0.021 | 1.000 | -0.414 | 1.000 | -0.585 | 0.323 | -0.297 | 0.666 | -0.111 | 0.900 | -1.071 | 0.011 | | |
