## Supplemental Table S3 for "Exposure of *Mycobacterium tuberculosis* to human alveolar lining fluid shows temporal and strain-specific adaptation to the lung environment"

**Supplemental Table S3**. Pathways identified by gene enrichment analysis in ShinyGO 0.77. Pathways are listed based on enrichment FDR (significant pathways in bold), and information on fold enrichment, genes, and number of DEGs included in the pathway is provided.

| Enrichment FDR | nGenes | Pathway Genes | Fold Enrichment | Pathway | Genes |
|---|---|---|---|---|---|
| **0.0000** | **10** | **13** | **7.9612** | **Mixed, incl. animal organ development, and coenzyme a transferase family i** | RV3548C RV3549C RV3550 RV3551 RV3552 RV3553 RV3559C RV3560C RV3561 RV3563 |
| **0.0007** | **6** | **6** | **10.3496** | **Coenzyme A transferase family I, and animal organ development** | RV3548C RV3549C RV3550 RV3551 RV3552 RV3553 |
| **0.0007** | **6** | **6** | **10.3496** | **Response to acidic ph** | RV3083 RV3084 RV3085 RV3087 RV3088 RV3089 |
| **0.0079** | **23** | **92** | **2.5874** | **Response to abiotic stimulus** | RV0352 RV0384C RV0467 RV0560C RV0573C RV0981 RV1129C RV1221 RV2031C RV2246 RV2623 RV2626C RV2710 RV3083 RV3084 RV3085 RV3086 RV3087 RV3088 RV3089 RV3286C RV3414C RV3734C |
| **0.0122** | **11** | **29** | **3.9257** | **Mixed, incl. cholesterol metabolic process, and duf35 ob-fold domain, acyl-coa-associated** | RV3523 RV3548C RV3549C RV3550 RV3551 RV3552 RV3553 RV3559C RV3560C RV3561 RV3563 |
| **0.0122** | **19** | **74** | **2.6573** | **Mixed, incl. fatty acid biosynthetic process, and nonribosomal peptide biosynthetic process** | RV1181 RV1182 RV1183 RV1342C RV2243 RV2244 RV2245 RV2246 RV2378C RV2382C RV2383C RV2524C RV2933 RV2934 RV2946C RV2947C RV3820C RV3823C RV3824C |
| **0.0122** | **7** | **12** | **6.0373** | **Response to acidic ph, and helix-turn-helix domain of resolvase** | RV3083 RV3084 RV3085 RV3086 RV3087 RV3088 RV3089 |
| **0.0122** | **5** | **6** | **8.6247** | **Mixed, incl. mycofactocin, and heme carboxy lyase-like** | RV0692 RV0693 RV0694 RV0695 RV0696 |
| **0.0122** | **9** | **19** | **4.9024** | **Response to ph** | RV0467 RV0981 RV3083 RV3084 RV3085 RV3086 RV3087 RV3088 RV3089 |
| **0.0144** | **23** | **101** | **2.3568** | **Transferase activity, transferring acyl groups** | RV0032 RV0129C RV0262C RV0889C RV1131 RV1181 RV1182 RV1569 RV2243 RV2245 RV2246 RV2482C RV2933 RV2934 RV2946C RV3087 RV3088 RV3089 RV3419C RV3556C RV3734C RV3820C RV3824C |
| **0.0167** | **8** | **17** | **4.8704** | **Mixed, incl. response to acidic ph, and wax metabolic process** | RV3083 RV3084 RV3085 RV3086 RV3087 RV3088 RV3089 RV3742C |
| **0.0205** | **19** | **78** | **2.5211** | **Transferase activity, transferring acyl groups other than amino-acyl groups** | RV0032 RV0129C RV0262C RV1182 RV1569 RV2243 RV2245 RV2246 RV2482C RV2933 RV2934 RV3087 RV3088 RV3089 RV3419C RV3556C RV3734C RV3820C RV3824C |
| **0.0264** | **15** | **55** | **2.8226** | **Fatty acid biosynthetic process, and lipid transport** | RV1181 RV1182 RV1183 RV2243 RV2244 RV2245 RV2246 RV2524C RV2933 RV2934 RV2946C RV2947C RV3820C RV3823C RV3824C |
| **0.0294** | **21** | **94** | **2.3121** | **Mixed, incl. fatty acid biosynthetic process, and acyltransferase** | RV0032 RV1181 RV1182 RV1183 RV1342C RV2243 RV2244 RV2245 RV2246 RV2378C RV2382C RV2383C RV2524C RV2933 RV2934 RV2946C RV2947C RV3734C RV3820C RV3823C RV3824C |
| **0.0443** | **8** | **20** | **4.1398** | **Beta-ketoacyl synthase, N-terminal** | RV1181 RV2245 RV2246 RV2382C RV2524C RV2933 RV2934 RV2947C |
| **0.0443** | **8** | **20** | **4.1398** | **Beta-ketoacyl synthase, C-terminal** | RV1181 RV2245 RV2246 RV2382C RV2524C RV2933 RV2934 RV2947C |
| **0.0443** | **8** | **20** | **4.1398** | **Polyketide synthase, beta-ketoacyl synthase domain** | RV1181 RV2245 RV2246 RV2382C RV2524C RV2933 RV2934 RV2947C |
| 0.0532 | 4 | 5 | 8.2797 | Mixed, incl. l-lysine 6-transaminase, and domain of unknown function duf1338 | RV3288C RV3289C RV3290C RV3292 |
| 0.0664 | 35 | 206 | 1.7584 | Cellular lipid metabolic process | RV0129C RV0211 RV0468 RV0669C RV0888 RV1013 RV1129C RV1131 RV1181 RV1182 RV1497 RV2243 RV2244 RV2245 RV2246 RV2247 RV2383C RV2482C RV2523C RV2933 RV2934 RV2946C RV2947C RV2958C RV3087 RV3088 RV3089 RV3548C RV3556C RV3559C RV3734C RV3807C RV3820C RV3823C RV3824C |
| 0.0814 | 12 | 45 | 2.7599 | Mixed, incl. cholesterol metabolic process, and aromatic hydrocarbons catabolism | RV3523 RV3548C RV3549C RV3550 RV3551 RV3552 RV3553 RV3557C RV3559C RV3560C RV3561 RV3563 |
| 0.0814 | 6 | 13 | 4.7767 | Mixed, incl. sulfolipid metabolic process, and methyl-branched fatty acid biosynthetic process | RV1181 RV1182 RV1183 RV3820C RV3823C RV3824C |
| 0.0814 | 3 | 3 | 10.3496 | tRNA threonylcarbamoyladenosine modification | RV3419C RV3421C RV3422C |
| 0.0814 | 22 | 112 | 2.0330 | Fatty acid metabolic process | RV0211 RV0468 RV0669C RV1013 RV1129C RV1131 RV1181 RV2243 RV2244 RV2245 RV2246 RV2247 RV2383C RV2482C RV2523C RV2933 RV2934 RV2946C RV3089 RV3548C RV3556C RV3559C |
| 0.0814 | 3 | 3 | 10.3496 | tRNA threonylcarbamoyladenosine metabolic process | RV3419C RV3421C RV3422C |
| 0.0814 | 18 | 83 | 2.2445 | Monocarboxylic acid biosynthetic process | RV0032 RV0468 RV0669C RV1013 RV1181 RV1569 RV2225 RV2243 RV2244 RV2245 RV2246 RV2247 RV2383C RV2523C RV2933 RV2934 RV2946C RV3089 |
| 0.0814 | 3 | 3 | 10.3496 | SpoVT-AbrB domain | RV0599C RV2166C RV2595 |
| 0.0814 | 3 | 3 | 10.3496 | SpoVT-AbrB domain superfamily | RV0599C RV2166C RV2595 |
| 0.0903 | 4 | 6 | 6.8997 | Alkaloid biosynthetic process, and acyl-coa dehydrogenase/oxidase, n-terminal | RV3559C RV3560C RV3561 RV3563 |
| 0.0903 | 4 | 6 | 6.8997 | Mixed, incl. probable membrane protein mt1774/rv1733c-like, and uncharacterised conserved protein ucp006404, peptidase m50/cbs | RV2624C RV2625C RV2626C RV2628 |
| 0.0903 | 4 | 6 | 6.8997 | Protein secretion by the type vii secretion system, and regulation of protein secretion | RV3612C RV3614C RV3615C RV3616C |
| 0.0904 | 6 | 14 | 4.4355 | Mixed, incl. universal stress protein a family, and cbs domain | RV2031C RV2623 RV2624C RV2625C RV2626C RV2628 |
| 0.0904 | 21 | 108 | 2.0124 | Mixed, incl. ppe superfamily, and pe-pgrs family, n-terminal | RV0151C RV0160C RV0304C RV0442C RV1168C RV1169C RV1268C RV1375 RV1376 RV1430 RV1542C RV1690 RV1917C RV2352C RV2353C RV3136 RV3558 RV3612C RV3614C RV3615C RV3616C |
| 0.0904 | 28 | 161 | 1.7999 | Lipid biosynthetic process | RV0129C RV0211 RV0468 RV0669C RV1013 RV1181 RV1182 RV2243 RV2244 RV2245 RV2246 RV2247 RV2383C RV2482C RV2523C RV2933 RV2934 RV2946C RV2947C RV2958C RV2959C RV3087 RV3088 RV3089 RV3734C RV3820C RV3823C RV3824C |
| 0.0904 | 8 | 24 | 3.4499 | Response to organic substance | RV0211 RV0467 RV0560C RV0888 RV3086 RV3548C RV3553 RV3559C |
| 0.0904 | 6 | 14 | 4.4355 | Regulation of dna-templated transcription, initiation | RV1189 RV1221 RV2166C RV2710 RV3286C RV3414C |

| | | | | | |
|---|---|---|---|---|---|
| 0.0904 | 18 | 87 | 2.1413 | Acyltransferase | RV0032 RV0129C RV0262C RV1181 RV1182 RV1569 RV2243 RV2245 RV2246 RV2482C RV2946C RV3087 RV3088 RV3419C RV3556C RV3734C RV3820C RV3824C |
| 0.0955 | 15 | 67 | 2.3171 | Fatty acid biosynthetic process | RV0468 RV0669C RV1013 RV1181 RV2243 RV2244 RV2245 RV2246 RV2247 RV2383C RV2523C RV2933 RV2934 RV2946C RV3089 |
| 0.1096 | 39 | 253 | 1.5954 | Lipid metabolic process | RV0129C RV0211 RV0468 RV0669C RV0888 RV1013 RV1129C RV1131 RV1181 RV1182 RV1497 RV2243 RV2244 RV2245 RV2246 RV2247 RV2277C RV2383C RV2482C RV2523C RV2933 RV2934 RV2946C RV2947C RV2958C RV2959C RV3087 RV3088 RV3089 RV3286C RV3548C RV3556C RV3559C RV3561 RV3734C RV3807C RV3820C RV3823C RV3824C |
| 0.1096 | 7 | 20 | 3.6224 | Response to temperature stimulus | RV0352 RV0384C RV1221 RV2031C RV2710 RV3286C RV3414C |
| 0.1096 | 13 | 55 | 2.4463 | Response to decreased oxygen levels | RV0467 RV0560C RV0573C RV1129C RV2031C RV2246 RV2623 RV2626C RV2710 RV3087 RV3088 RV3286C RV3734C |
| 0.1096 | 7 | 20 | 3.6224 | ATPase family associated with various cellular activities (AAA) | RV0001 RV0384C RV2115C RV2457C RV2592C RV2897C RV3585 |
| 0.1096 | 8 | 25 | 3.3119 | Beta-ketoacyl synthase, N-terminal domain | RV2245 RV2246 RV2382C RV2524C RV2933 RV2934 RV2947C RV3556C |
| 0.1096 | 7 | 20 | 3.6224 | Beta-ketoacyl synthase, C-terminal domain | RV2245 RV2246 RV2382C RV2524C RV2933 RV2934 RV2947C |
| 0.1203 | 29 | 174 | 1.7249 | Symbiotic process | RV0211 RV0442C RV0467 RV0468 RV0549C RV0573C RV1129C RV1221 RV1349 RV1522C RV1917C RV2031C RV2383C RV2626C RV2659C RV3083 RV3084 RV3085 RV3086 RV3087 RV3088 RV3089 RV3151 RV3321C RV3551 RV3552 RV3615C RV3734C RV3810 |
| 0.1329 | 13 | 57 | 2.3604 | Response to oxygen levels | RV0467 RV0560C RV0573C RV1129C RV2031C RV2246 RV2623 RV2626C RV2710 RV3087 RV3088 RV3286C RV3734C |
| 0.1329 | 10 | 38 | 2.7236 | Thiolase-like | RV1181 RV2245 RV2246 RV2382C RV2524C RV2933 RV2934 RV2947C RV3523 RV3556C |
| 0.1537 | 28 | 170 | 1.7046 | Monocarboxylic acid metabolic process | RV0032 RV0211 RV0467 RV0468 RV0669C RV0694 RV1013 RV1129C RV1131 RV1181 RV1569 RV2225 RV2243 RV2244 RV2245 RV2246 RV2247 RV2383C RV2482C RV2523C RV2933 RV2934 RV2946C RV3089 RV3290C RV3548C RV3556C RV3559C |
| 0.1537 | 45 | 311 | 1.4975 | Interspecies interaction between organisms | RV0072 RV0211 RV0285 RV0288 RV0442C RV0467 RV0468 RV0549C RV0573C RV0867C RV0888 RV0981 RV1129C RV1198 RV1221 RV1349 RV1522C RV1917C RV2031C RV2115C RV2220 RV2383C RV2623 RV2626C RV2659C RV2958C RV3083 RV3084 RV3085 RV3086 RV3087 RV3088 RV3089 RV3151 RV3270 RV3303C RV3321C RV3414C RV3551 RV3552 RV3614C RV3615C RV3616C RV3734C RV3810 |
| 0.1567 | 9 | 33 | 2.8226 | Mixed, incl. stas domain, and sigma factor | RV0249C RV0250C RV0251C RV0516C RV0981 RV1904 RV3286C RV3287C RV3414C |
| 0.1573 | 12 | 52 | 2.3884 | Response to hypoxia | RV0467 RV0560C RV0573C RV1129C RV2031C RV2246 RV2623 RV2626C RV2710 RV3087 RV3088 RV3734C |
| 0.1671 | 14 | 66 | 2.1954 | Mixed, incl. pentapeptide repeats (8 copies), and pe-pgrs family, n-terminal | RV0151C RV0160C RV0304C RV0442C RV1268C RV1375 RV1376 RV1430 RV1690 RV1917C RV2352C RV2353C RV3136 RV3558 |
| 0.1827 | 55 | 403 | 1.4125 | Response to stimulus | RV0129C RV0211 RV0262C RV0352 RV0384C RV0442C RV0467 RV0468 RV0549C RV0560C RV0573C RV0821C RV0888 RV0981 RV1129C RV1221 RV1349 RV1522C RV1542C RV1636 RV1917C RV2031C RV2115C RV2136C RV2191 RV2246 RV2359 RV2383C RV2429 RV2592C RV2623 RV2626C RV2674 RV2710 RV2985 RV3083 RV3084 RV3085 RV3086 RV3087 RV3088 RV3089 RV3143 RV3219 RV3286C RV3321C RV3414C RV3548C RV3553 RV3556C RV3559C RV3585 RV3615C RV3734C RV3855 |
| 0.1890 | 4 | 8 | 5.1748 | Fatty acid elongation | RV0468 RV2245 RV2246 RV2247 |
| 0.1890 | 4 | 8 | 5.1748 | Condensation domain | RV1182 RV2383C RV3820C RV3824C |
| 0.1890 | 4 | 8 | 5.1748 | L-lysine 6-monooxygenase (NADPH-requiring) | RV2378C RV3049C RV3083 RV3303C |
| 0.1912 | 5 | 13 | 3.9806 | Fatty acid biosynthesis, and phenolic phthiocerol biosynthetic process | RV2243 RV2244 RV2245 RV2246 RV2524C |
| 0.1912 | 10 | 41 | 2.5243 | Mixed, incl. pentapeptide repeats (8 copies), and pe-ppe, c-terminal | RV0151C RV0160C RV0304C RV0442C RV1268C RV1430 RV1690 RV2352C RV2353C RV3136 |
| 0.1912 | 5 | 13 | 3.9806 | DNA-templated transcription, initiation | RV1189 RV1221 RV2710 RV3286C RV3414C |
| 0.1912 | 6 | 18 | 3.4499 | Response to heat | RV0352 RV0384C RV1221 RV2031C RV2710 RV3414C |
| 0.1912 | 5 | 13 | 3.9806 | Sigma factor activity | RV1189 RV1221 RV2710 RV3286C RV3414C |
| 0.1912 | 27 | 167 | 1.6733 | Response to chemical | RV0129C RV0211 RV0262C RV0467 RV0560C RV0821C RV0888 RV1221 RV1542C RV1636 RV2031C RV2115C RV2136C RV2359 RV2429 RV2710 RV3086 RV3087 RV3088 RV3219 RV3286C RV3548C RV3553 RV3556C RV3559C RV3734C RV3855 |
| 0.1912 | 12 | 55 | 2.2581 | Cellular response to chemical stimulus | RV0211 RV0467 RV0888 RV1542C RV2031C RV2115C RV2429 RV3086 RV3548C RV3553 RV3556C RV3559C |
| 0.1912 | 5 | 13 | 3.9806 | RNA polymerase sigma-70 region 2 | RV1189 RV1221 RV2710 RV3286C RV3414C |
| 0.1912 | 5 | 13 | 3.9806 | RNA polymerase sigma factor, region 3/4-like | RV1189 RV1221 RV2710 RV3286C RV3414C |
| 0.1912 | 5 | 13 | 3.9806 | RNA polymerase sigma factor, region 2 | RV1189 RV1221 RV2710 RV3286C RV3414C |
| 0.1912 | 5 | 13 | 3.9806 | RNA polymerase sigma-70 like domain | RV1189 RV1221 RV2710 RV3286C RV3414C |
| 0.1912 | 5 | 13 | 3.9806 | Sigma factor | RV1189 RV1221 RV2710 RV3286C RV3414C |
| 0.1912 | 5 | 13 | 3.9806 | Sigma-70 region 2 | RV1189 RV1221 RV2710 RV3286C RV3414C |
| 0.1958 | 30 | 194 | 1.6005 | Response to stress | RV0211 RV0352 RV0384C RV0467 RV0560C RV0573C RV1129C RV1221 RV1349 RV1542C RV2031C RV2115C RV2136C RV2191 RV2246 RV2383C RV2429 RV2592C RV2623 RV2626C RV2674 RV2710 RV2985 RV3087 RV3088 RV3219 RV3286C RV3414C RV3585 RV3734C |
