## Supplemental Table S4 for "Exposure of *Mycobacterium tuberculosis* to human alveolar lining fluid shows temporal and strain-specific adaptation to the lung environment"

**Supplemental Table S4**. Strain-specific DEGs at 15 min and 12 h post-ALF exposure. Locus tag, log2FC (ALF-exposed *vs* . unexposed *M.tb* , only significant DEGs are shown as defined in the methods, upregulated genes in yellow and downregulated genes in blue), product, functional category and function are provided for each of the DEGs.

| Locus tag | Name | log2FC | Product | Functional Category (Mycobrowser) | Function (Mycobrowser) |
|---|---|---|---|---|---|
| **CDC1551** | | | | | |
| Rv0867c | *rpfA* | **-1.154** | Possible resuscitation-promoting factor RpfA | cell wall and cell processes | Unknown. May promote the resuscitation and growth of dormant, nongrowing cell. |
| Rv0888 | *Rv0888* | **-1.389** | Probable exported protein | cell wall and cell processes | Unknown |
| Rv2136c | *Rv2136c or bacA or upk* | **1.015** | Possible conserved transmembrane protein | cell wall and cell processes | Unknown |
| Rv1078 | *Rv1078* | **-1.041** | Probable proline-rich antigen homolog | conserved hypotheticals | Unknown |
| Rv1535 | *Rv1535* | **-2.068** | Unknown protein | conserved hypotheticals | Unknown |
| Rv2042c | *Rv2042c* | **1.001** | Conserved protein | conserved hypotheticals | Unknown |
| Rv2225 | *panB* | **-1.148** | Conserved protein | conserved hypotheticals | Unknown |
| Rv2323c | *Rv2323c* | **-1.159** | Conserved protein | conserved hypotheticals | Function unknown |
| Rv2360c | *Rv2360c* | **1.004** | Unknown protein | conserved hypotheticals | Unknown |
| Rv3486 | *Rv3486* | **1.390** | Conserved protein | conserved hypotheticals | Function unknown |
| Rv3555c | *Rv3555c* | **1.088** | Conserved protein | conserved hypotheticals | Function unknown |
| Rv2441c | *rpmA* | **-1.029** | 50S ribosomal protein L27 RpmA | information pathways | Involved in translation mechanisms. |
| Rv3442c | *rpsI* | **-1.027** | 30S ribosomal protein S9 RpsI | information pathways | Involved in translation mechanism. This protein is one of the assembly proteins of the 50S ribosomal subunit. |
| Rv1882c | *Rv1882c* | **1.008** | Probable short-chain type dehydrogenase/reductase | int. metabolism & respiration | Function unknown; probably involved in cellular metabolism |
| Rv2220 | *glnA1* | **-1.046** | Glutamine synthetase GlnA1 (glutamine synthase) (GS-I) | int. metabolism & respiration | Involved in glutamine biosynthesis [catalytic activity: ATP + L-glutamate + NH(3) = ADP + glutamine + orthophosphate]. |
| Rv2368c | *phoH1* | **1.066** | Probable PHOH-like protein PhoH1 (phosphate starvation-inducible protein PSIH) | int. metabolism & respiration | Unknown |
| Rv0468 | *fadB2* | **-1.004** | 3-hydroxybutyryl-CoA dehydrogenase FadB2 (beta-hydroxybutyryl-CoA dehydrogenase) (BHBD) | lipid metabolism | Butyrate/butanol-producing pathway [catalytic activity: (S)-3-hydroxybutanoyl-CoA + NADP+ = 3-acetoacetyl-CoA + NADPH] |
| Rv1013 | *pks16* | **-1.039** | Putative polyketide synthase Pks16 | lipid metabolism | Potentially involved in some intermediate steps for the synthesis of a polyketide molecule which may be involved in secondary metabolism. |
| Rv1153c | *omt* | **1.008** | Probable O-methyltransferase Omt | lipid metabolism | Function unknown, but supposedly involved in lipid metabolism |

| | | | | | |
|---|---|---|---|---|---|
| Rv2244 | *acpM* | **-1.258** | 3-oxoacyl-[acyl-carrier protein] synthase 1 KasA (beta-ketoacyl-ACP synthase) (KAS I) | lipid metabolism | Involved in fatty acid biosynthesis (mycolic acids synthesis); involved in meromycolate extension. Catalyzes the condensation reaction of fatty acid synthesis by the addition to an acyl acceptor of two carbons from malonyl-ACP [catalytic activity: acyl-[acyl-carrier protein] + malonyl-[acyl-carrier protein] = 3-oxoacyl-[acyl-carrier protein] + [acyl-carrier protein] + CO(2)]. |
| Rv2245 | *kasA* | **-1.476** | 3-oxoacyl-[acyl-carrier protein] synthase 2 KasB (beta-ketoacyl-ACP synthase) (KAS I) | lipid metabolism | Involved in fatty acid biosynthesis (mycolic acids synthesis); involved in meromycolate extension. Catalyzes the condensation reaction of fatty acid synthesis by the addition to an acyl acceptor of two carbons from malonyl-ACP [catalytic activity: acyl-[acyl-carrier protein] + malonyl-[acyl-carrier protein] = 3-oxoacyl-[acyl-carrier protein] + [acyl-carrier protein] + CO(2)]. |
| Rv2246 | *kasB* | **-1.200** | Acetyl/propionyl-CoA carboxylase (beta subunit) AccD6 | lipid metabolism | Involved in fatty acid biosynthesis (mycolic acids synthesis) [catalytic activity: ATP + propionyl-CoA + CO(2) + H(2)O = ADP + orthophosphate + methylmalonyl-CoA]. |
| Rv3136 | *PPE51* | **-1.320** | PPE family protein PPE51 | PE/PPE | Function unknown |
| Rv2359 | *zur or fur-2 or furB* | **1.160** | Probable zinc uptake regulation protein Zur | regulatory proteins | Acts as a global negative controlling element, employing Zn(2+) as a cofactor to bind the operator of the repressed genes. |
| Rv3269 | *Rv3269* | **1.464** | Conserved protein | virulence, detoxification, adaptation | Function unknown. May be involved in a chaperoning process. |
| **H37Rv** | | | | | |
| Rv2959c | *Rv2959c* | **-1.193** | Possible methyltransferase (methylase) | int. metabolism & respiration | Thought to cause methylation. |
| Rv3559c | *Rv3559c* | **1.300** | Probable oxidoreductase | int. metabolism & respiration | Function unknown; probably involved in cellular metabolism. |
| Rv3560c | *fadE30* | **1.530** | Probable acyl-CoA dehydrogenase FadE30 | lipid metabolism | Function unknown, but involved in lipid degradation. |
| **HN878** | | | | | |
| Rv0072 | *Rv0072* | **5.728** | Probable glutamine-transport transmembrane protein ABC transporter | cell wall and cell processes | Thought to be involved in active transport of glutamine across the membrane (import). Responsible for the translocation of the substrate across the membrane. |
| **W-7642** | | | | | |
| Rv3086 | *adhD* | **1.088** | Probable zinc-type alcohol dehydrogenase AdhD (aldehyde reductase) | int. metabolism & respiration | Function unknown; generates an aldehyde (or perhaps a ketone) from an alcohol. |
| Rv3087 | *Rv3087* | **1.166** | Possible triacylglycerol synthase (diacylglycerol acyltransferase) | lipid metabolism | May be involved in synthesis of triacylglycerol |
| Rv3088 | *tgs4* | **1.155** | Putative triacylglycerol synthase (diacylglycerol acyltransferase) Tgs4 | lipid metabolism | May be involved in synthesis of triacylglycerol |

**Supplemental Table S4**. Strain-specific DEGs at 15 min and 12 h post-ALF exposure. Locus tag, log2FC (ALF-exposed *vs* . unexposed *M.tb* , only significant DEGs are shown as defined in the methods, upregulated genes in yellow and downregulated genes in blue), product, functional category and function are provided for each of the DEGs.

| Locus tag | Gene name | log2FC | Product | Functional Category (Mycobrowser) | Function (Mycobrowser) |
|---|---|---|---|---|---|
| **CDC1551** | | | | | |
| Rv0488 | *Rv0488* | **1.140** | Probable conserved integral membrane protein | cell wall and cell processes | Unknown; possibly involved in transport of lysine across the membrane. |
| Rv0531 | *Rv0531* | **-1.102** | Possible conserved membrane protein | cell wall and cell processes | Unknown |
| Rv0559c | *Rv0559c* | **-1.182** | Possible conserved secreted protein | cell wall and cell processes | Unknown |
| Rv0655 | *mkl* | **-1.465** | Possible ribonucleotide-transport ATP-binding protein ABC transporter Mkl | cell wall and cell processes | Thought to be involved in active transport of ribonucleotide across the membrane. Responsible for energy coupling to the |
| Rv0821c | *phoY2* | **-1.022** | Probable phosphate-transport system transcriptional regulatory protein PhoY2 | cell wall and cell processes | Involved in transcriptional regulation of active transport of inorganic phosphate across the membrane. |
| Rv1440 | *secG* | **-1.228** | Probable protein-export membrane protein (translocase subunit) SecG | cell wall and cell processes | Involved in protein export. Participates in a early event of protein translocation. |
| Rv3270 | *ctpC* | **1.636** | Probable metal cation-transporting P-type ATPase C CtpC | cell wall and cell processes | Metal cation-transporting ATPase; possibly catalyzes the transport of undetermined metal cation with the hydrolyse of ATP [catalytic activity: ATP + H(2)O + undetermined metal cation(in) = ADP + phosphate + undetermined metal |
| Rv3330 | *dacB1* | **1.307** | Probable penicillin-binding protein DacB1 (D-alanyl-D-alanine carboxypeptidase) (DD-peptidase) (DD-carboxypeptidase) (PBP) (DD-transpeptidase) (serine-type D-ala-D-ala carboxypeptidase) (D-amino acid hydrolase) | cell wall and cell processes | Involved in peptidoglycan synthesis (at final stages). Hydrolyzes the bound D-alanyl-D-alanine [catalytic activity: D-alanyl-D-alanine + H(2)O = 2 D-alanine]. |
| Rv3679 | *Rv3679* | **-1.251** | Probable anion transporter ATPase | cell wall and cell processes | Anion-transporting ATPase; supposedly catalyzes the extrusion of undetermined anions [catalytic activity: ATP + H(2)O + undetermined anion(in) = ADP + phosphate + undetermined |
| Rv0057 | *Rv0057* | **1.155** | Hypothetical protein | conserved hypotheticals | Unknown |
| Rv0059 | *Rv0059* | **1.132** | Hypothetical protein | conserved hypotheticals | Unknown |
| Rv0140 | *Rv0140* | **-1.410** | Conserved protein | conserved hypotheticals | Function unknown |
| Rv0424c | *Rv0424c* | **-1.213** | Hypothetical protein | conserved hypotheticals | Unknown |
| Rv0695 | *Rv0695 or mftE* | **-1.201** | Conserved hypothetical protein | conserved hypotheticals | Function unknown |
| Rv1171 | *Rv1171* | **-1.020** | Conserved hypothetical protein | conserved hypotheticals | Unknown |
| Rv1265 | *Rv1265* | **-1.106** | Unknown protein | conserved hypotheticals | Unknown. Seems to be expressed during macrophage |
| Rv1638A | *Rv1638A* | **-1.323** | Conserved hypothetical protein | conserved hypotheticals | Function unknown |
| Rv1907c | *Rv1907c* | **-1.102** | Hypothetical protein | conserved hypotheticals | Unknown |
| Rv2189c | *Rv2189c* | **1.709** | Conserved hypothetical protein | conserved hypotheticals | Unknown |
| Rv2331 | *Rv2331* | **-1.468** | Hypothetical protein | conserved hypotheticals | Unknown |
| Rv2407 | *Rv2407* | **1.144** | Conserved hypothetical protein | conserved hypotheticals | Function unknown |
| Rv2466c | *Rv2466c* | **-1.375** | Conserved protein | conserved hypotheticals | Function unknown. Seems regulated by sigh (Rv3223c |
| Rv2517c | *Rv2517c* | **-1.200** | Unknown protein | conserved hypotheticals | Unknown |
| Rv3190A | *Rv3190A* | **-1.100** | Conserved protein | conserved hypotheticals | Unknown |

| | | | | | |
|---|---|---|---|---|---|
| Rv3463 | *Rv3463* | **-1.211** | Conserved protein | conserved hypotheticals | Function unknown |
| Rv1221 | *sigE* | **-1.019** | Alternative RNA polymerase sigma factor SigE | information pathways | The sigma factor is an initiation factor that promotes attachment of the RNA polymerase to specific initiation sites and then is released. Seems to be regulated by sigh (Rv3223c product). Seems to regulate the heat-shock response. |
| Rv0605 | *Rv0605* | **-1.055** | Possible resolvase | insertion seqs and phages | Prevents the cointegration of foreign DNA before integration into the chromosome. |
| Rv1042c | *Rv1042c* | **-1.913** | Probable is like-2 transposase | insertion seqs and phages | Possibly required for the transposition of an insertion element. |
| Rv3467 | *Rv3467* | **1.430** | Conserved hypothetical protein | insertion seqs and phages | Unknown |
| Rv0694 | *Rv0694 or mftD* | **-2.172** | Possible L-lactate dehydrogenase (cytochrome) LldD1 | int. metabolism & respiration | Involved in respiration; catalyzes conversion of lactate into pyruvate [catalytic activity: (S)-lactate + 2 ferricytochrome C = pyruvate + 2 ferrocytochrome C]. |
| Rv0696 | *Rv0696 or mftE* | **-1.110** | Probable membrane sugar transferase | int. metabolism & respiration | Function unknown; probably involved in cellular metabolism. |
| Rv0865 | *mog* | **1.089** | Probable molybdopterin biosynthesis Mog protein | int. metabolism & respiration | Involved in molybdopterin biosynthesis; involved in the biosynthesis of a demolybdo-cofactor (molybdopterin), |
| Rv1652 | *argC* | **1.070** | Probable N-acetyl-gamma-glutamyl-phoshate reductase ArgC | int. metabolism & respiration | Involved in arginine biosynthesis (at the third step) [catalytic activity: N-acetyl-L-glutamate 5-semialdehyde + NADP(+) + phosphate = N-acetyl-5-glutamyl phosphate + NADPH]. |
| Rv2674 | *msrB* | **-1.105** | Probable peptide methionine sulfoxide reductase MsrB (protein-methionine-R-oxide reductase) (peptide met(O) reductase) | int. metabolism & respiration | Has an important function as a repair enzyme for proteins that have been inactivated by oxidation. Catalyzes the reversible oxidation-reduction of methionine sulfoxide in proteins to methionine [catalytic activity: protein L-methionine + oxidized thioredoxin + H2O = protein-L-methionine-(R)-S-oxide + |
| Rv2958c | *Rv2958c* | **1.259** | Possible glycosyl transferase | int. metabolism & respiration | Function unknown; probably involved in cellular metabolism. Possibly involved in resistance to killing by human |
| Rv2959c | *Rv2959c* | **1.138** | Possible methyltransferase (methylase) | int. metabolism & respiration | Thought to cause methylation. |
| Rv3145 | *nuoA* | **-1.205** | Probable NADH dehydrogenase I (chain A) NuoA (NADH-ubiquinone oxidoreductase chain A) | int. metabolism & respiration | Involved in aerobic\|anaerobic respiration [catalytic activity: NADH + ubiquinone = NAD(+) + ubiquinol]. |
| Rv1013 | *pks16* | **1.336** | Putative polyketide synthase Pks16 | lipid metabolism | Potentially involved in some intermediate steps for the synthesis of a polyketide molecule which may be involved in |
| Rv2243 | *fabD* | **1.184** | Meromycolate extension acyl carrier protein AcpM | lipid metabolism | Involved in fatty acid biosynthesis (mycolic acids synthesis); involved in meromycolate extension. |
| Rv2244 | *acpM* | **1.038** | 3-oxoacyl-[acyl-carrier protein] synthase 1 KasA (beta-ketoacyl-ACP synthase) (KAS I) | lipid metabolism | Involved in fatty acid biosynthesis (mycolic acids synthesis); involved in meromycolate extension. Catalyzes the condensation reaction of fatty acid synthesis by the addition to an acyl acceptor of two carbons from malonyl-ACP [catalytic activity: acyl-[acyl-carrier protein] + malonyl-[acyl-carrier |
| Rv2590 | *fadD9* | **-1.221** | Probable fatty-acid-CoA ligase FadD9 (fatty-acid-CoA synthetase) (fatty-acid-CoA synthase) | lipid metabolism | Function unknown, but involved in lipid degradation. |
| Rv3523 | *ltp3* | **-1.026** | Probable lipid carrier protein or keto acyl-CoA thiolase Ltp3 | lipid metabolism | Function unknown; probably involved in lipid metabolism. |
| Rv1168c | *PPE17* | **-1.448** | PPE family protein PPE17 | PE/PPE | Function unknown |
| Rv1169c | *lipX or PE11* | **-1.438** | PE family protein. Possible lipase LipX. | PE/PPE | Function unknown |

| | | | | | |
|---|---|---|---|---|---|
| Rv3746c | *PE34* | **-1.567** | Probable PE family protein PE34 (PE family-related protein) | PE/PPE | Function unknown |
| Rv1129c | *Rv1129c* | **-1.494** | Probable transcriptional regulator protein | regulatory proteins | Involved in transcriptional mechanism |
| Rv1534 | *Rv1534* | **-1.312** | Probable transcriptional regulator | regulatory proteins | Possibly involved in a transcriptional mechanism |
| Rv1776c | *Rv1776c* | **1.116** | Possible transcriptional regulatory protein | regulatory proteins | Involved in transcriptional mechanism. |
| Rv3557c | *Rv3557c* | **1.204** | Transcriptional regulatory protein (probably TetR-family) | regulatory proteins | Involved in transcriptional mechanism. |
| Rv3833 | *Rv3833* | **-1.276** | Transcriptional regulatory protein (probably AraC-family) | regulatory proteins | Involved in a transcriptional mechanism. |
| Rv0352 | *dnaJ1* | **-1.040** | Probable chaperone protein DnaJ1 | virulence, detoxification, adaptation | Acts as a co-chaperone. Stimulates, jointly with GRPE\|Rv0351, the ATPase activity of DNAK\|Rv0350. Seems to be regulated negatively by HSPR (Rv0353 product). |
| Rv0353 | *hspR* | **-1.319** | Probable heat shock protein transcriptional repressor HspR (MerR family) | virulence, detoxification, adaptation | Involved in transcriptional regulation (repression) of heat shock proteins e.g. DNAK\|Rv0350, GRPE\|Rv0351, DNAJ1\|Rv0352. Binds to three inverted repeats (IR1-IR3) in the promoter region of the DNAK operon. Induction: by heat shock. |
| Rv0384c | *clpB or htpM* | **-1.627** | Probable endopeptidase ATP binding protein (chain B) ClpB (ClpB protein) (heat shock protein F84.1) | virulence, detoxification, adaptation | Thought to be an ATPase subunit of an intracellular ATP-dependent protease. Seems to be regulated positively by sigh (Rv3223c product) and negatively by HSPR (Rv0353 product). |
| Rv0599c | *vapB27* | **-1.171** | Possible antitoxin VapB27 | virulence, detoxification, adaptation | Unknown |
| Rv2104c | *vapB37* | **-1.037** | Possible antitoxin VapB37 | virulence, detoxification, adaptation | Unknown |
| Rv3385c | *vapB46* | **-1.232** | Possible antitoxin VapB46 | virulence, detoxification, adaptation | Unknown |
| Rv3697A | *vapB48* | **-1.159** | Possible antitoxin VapB48 | virulence, detoxification, adaptation | Unknown |
| **H37Rv** | | | | | |
| Rv0063a | *Rv0063a* | **-1.905** | | | |
| Rv0188 | *Rv0188* | **-1.309** | Probable conserved transmembrane protein | cell wall and cell processes | Unknown |
| Rv0309 | *Rv0309* | **-1.020** | Possible conserved exported protein | cell wall and cell processes | Unknown |
| Rv0870c | *Rv0870c* | **-1.004** | Possible conserved integral membrane protein | cell wall and cell processes | Unknown |
| Rv1349 | *irtB* | **1.112** | Iron-regulated transporter IrtB | cell wall and cell processes | Involved in iron homeostasis. Responsible for energy coupling to the transport system and for the translocation of the substrate across the membrane. |
| Rv2625c | *Rv2625c* | **1.052** | Probable conserved transmembrane alanine and leucine rich protein | cell wall and cell processes | Unknown |
| Rv2709 | *Rv2709* | **-1.109** | Probable conserved transmembrane protein | cell wall and cell processes | Unknown |
| Rv0762c | *Rv0762c* | **-1.016** | Conserved hypothetical protein | conserved hypotheticals | Function unknown |
| Rv1913 | *Rv1913* | **1.228** | Conserved hypothetical protein | conserved hypotheticals | Unknown |
| Rv2626c | *hrp1* | **1.404** | Hypoxic response protein 1 Hrp1 | conserved hypotheticals | Function unknown |
| Rv3026c | *Rv3026c* | **1.210** | Conserved hypothetical protein | conserved hypotheticals | Function unknown |
| Rv3354 | *Rv3354* | **-1.854** | Conserved hypothetical protein | conserved hypotheticals | Function unknown |
| Rv0717 | *rpsN1* | **-1.077** | 30S ribosomal protein S14 RpsN1 | information pathways | Known to be required for the assembly of 30S particles and may also be responsible for determining the conformation of |

| | | | | | |
|---|---|---|---|---|---|
| Rv2710 | *sigB or mysB* | **-1.107** | RNA polymerase sigma factor SigB | information pathways | The sigma factor is an initiation factor that promotes attachment of the RNA polymerase to specific initiation sites and then is released. May control the regulons of stationary phase and general stress resistance. Seems to be regulated by sigh (Rv3223c product) and SIGE (Rv1221 product). Seems to [illegible] |
| Rv3414c | *sigD* | **-1.004** | Probable alternative RNA polymerase sigma-D factor SigD | information pathways | The sigma factor is an initiation factor that promotes attachment of the RNA polymerase to specific initiation sites |
| Rv3750c | *Rv3750c* | **-1.093** | Possible excisionase | insertion seqs and phages | Sequence excision. |
| Rv0428c | *Rv0428c* | **1.330** | GCN5-related N-acetyltransferase | int. metabolism & respiration | Acetylation, substrate unknown |
| Rv0573c | *pncB2* | **1.020** | Nicotinic acid phosphoribosyltransferase PncB2 | int. metabolism & respiration | Involved in NAD salvage. Phosphoribosylation of nicotinic acid. [catalytic activity: nicotinate + 5-phosphoribosyl 1-pyrophosphate = nicotinate mononucleotide + diphosphate] |
| Rv0770 | *Rv0770* | **-1.112** | Probable dehydrogenase/reductase | int. metabolism & respiration | Function unknown; 3-hydroxyisobutyrate dehydrogenase family protein probably involved in cellular metabolism. |
| Rv1050 | *Rv1050* | **1.868** | Probable oxidoreductase | int. metabolism & respiration | Function unknown; probably involved in cellular metabolism |
| Rv2111c | *pup* | **-1.233** | Prokaryotic ubiquitin-like protein Pup | int. metabolism & respiration | Involved in proteasomal degradation. Covalently binds to protein substrates. |
| Rv2121c | *hisG* | **1.360** | ATP phosphoribosyltransferase HisG | int. metabolism & respiration | Involved in histidine biosynthesis. |
| Rv2277c | *Rv2277c* | **2.530** | Possible glycerolphosphodiesterase | int. metabolism & respiration | Unknown |
| Rv3293 | *pcd or aldB* | **-1.104** | Probable piperideine-6-carboxilic acid dehydrogenase Pcd (piperideine-6-carboxylate dehydrogenase) | int. metabolism & respiration | Involved in L-alpha-aminoadipic acid (L-AAA) biosynthesis (in the second step; the first step is promoted by lat enzyme. |
| Rv3303c | *lpdA* | **1.012** | NAD(P)H quinone reductase LpdA | int. metabolism & respiration | Involved in energy metabolism. Can catalyze the reduction of electron acceptors such as 2,6-dimethyl-1,4-benzoquinone (DMBQ) and 5-hydroxy-1,4-napthaquinone (5-HNQ) [catalytic activity: NAD(P)H + a quinone + H+ -> a quinol + NAD(P)+]. |
| Rv0119 | *fadD7* | **1.090** | Probable fatty-acid-CoA ligase FadD7 (fatty-acid-CoA synthetase) (fatty-acid-CoA synthase) | lipid metabolism | Function unknown, but involved in lipid degradation. |
| Rv2378c | *mbtG* | **1.073** | Lysine-N-oxygenase MbtG (L-lysine 6-monooxygenase) (lysine N6-hydroxylase) | lipid metabolism | Involved in the biogenesis of the hydroxyphenyloxazoline-containing siderophore mycobactins. This hydroxylase is possibly required for N-hydroxylation of the two lysine residues at some stage during mycobactin assembly [catalytic activity: L-lysine + O(2) = N6-hydroxy-L-lysine + H(2)O. No information can be found if this enzyme is NADPH dependent or [illegible] |
| Rv2352c | *PPE38* | **-1.131** | PPE family protein PPE38 | PE/PPE | Function unknown |
| Rv2034 | *Rv2034* | **-1.173** | ArsR repressor protein | regulatory proteins | Involved in transcriptional regulation. |
| Rv0549c | *vapC3* | **-1.106** | Possible toxin VapC3 | virulence, detoxification, adaptation | Unknown |
| Rv0959A | *vapB9* | **-1.402** | Possible antitoxin VapB9 | virulence, detoxification, adaptation | Unknown |
| Rv1720c | *vapC12* | **-1.474** | Possible toxin VapC12 | virulence, detoxification, adaptation | Unknown |
| Rv2319c | *Rv2319c* | **-1.337** | Universal stress protein family protein | virulence, detoxification, adaptation | Unknown |
| Rv2595 | *vapB40* | **-1.428** | Possible antitoxin VapB40 | virulence, detoxification, adaptation | Unknown |
| Rv2623 | *TB31.7* | **1.006** | Universal stress protein family protein TB31.7 | virulence, detoxification, adaptation | Function unknown |
| Rv2865 | *relF or relB2* | **-1.011** | Antitoxin RelF | virulence, detoxification, adaptation | Function unknown |

| | | | | | |
|---|---|---|---|---|---|
| Rv3473c | *bpoA* | **1.679** | Possible peroxidase BpoA (non-haem peroxidase) | virulence, detoxification, adaptation | Supposedly involved in detoxification reactions. |
| **HN878** | | | | | |
| Rv0114 | *gmhB* | **1.017** | Possible D-alpha,beta-D-heptose-1,7-biphosphate phosphatase GmhB (D-glycero-D-manno-heptose 7-phosphate kinase) | cell wall and cell processes | Involved in biosynthesis of nucleotide-activated glycero-manno-heptose. Involved in two pathways, D-alpha-D pathway [catalytic activity: D-glycero-alpha-D-manno-heptose 1,7-biphosphate = D-glycero-alpha-D-manno-heptose 1-phosphate] and L-beta-D pathway [catalytic activity: D-glycero-beta-D-manno-heptose 1,7-biphosphate = D-glycero-beta-D-manno- [illegible] |
| Rv0267 | *narU* | **1.897** | Probable integral membrane nitrite extrusion protein NarU (nitrite facilitator) | cell wall and cell processes | Involved in excretion of nitrite produced by the dissimilatory reduction of nitrate. Responsible for the translocation of the substrate across the membrane. |
| Rv1037c | *esxI* | **1.515** | Putative ESAT-6 like protein EsxI (ESAT-6 like protein 1) | cell wall and cell processes | Unknown |
| Rv1517 | *Rv1517* | **-1.021** | Conserved hypothetical transmembrane protein | cell wall and cell processes | Unknown |
| Rv1686c | *Rv1686c* | **2.378** | Probable conserved integral membrane protein ABC transporter | cell wall and cell processes | Thought to be involved in active transport of undetermined substrate (possibly drug) across the membrane. Responsible for the translocation of the substrate across the membrane. |
| Rv1687c | *Rv1687c* | **2.859** | Probable conserved ATP-binding protein ABC transporter | cell wall and cell processes | Thought to be involved in active transport of undetermined substrate (possibly drug) across the membrane. Responsible for energy coupling to the transport system. |
| Rv2080 | *lppJ* | **1.714** | Lipoprotein LppJ | cell wall and cell processes | Unknown |
| Rv2254c | *Rv2254c* | **-1.104** | Probable integral membrane protein | cell wall and cell processes | Unknown |
| Rv0358 | *Rv0358* | **1.046** | Conserved protein | conserved hypotheticals | Function unknown |
| Rv0470A | *Rv0470A* | **-1.153** | Hypothetical protein | conserved hypotheticals | Unknown |
| Rv0963c | *Rv0963c* | **1.770** | Conserved hypothetical protein | conserved hypotheticals | Function unknown |
| Rv2288 | *Rv2288* | **1.762** | Hypothetical protein | conserved hypotheticals | Unknown |
| Rv2917 | *Rv2917* | **1.666** | Conserved hypothetical alanine and arginine rich protein | conserved hypotheticals | Function unknown |
| Rv2659c | *Rv2659c* | **1.685** | Probable PhiRv2 prophage integrase | insertion seqs and phages | Sequence integration. Integrase is necessary for integration of a phage into the host genome by site-specific recombination. In conjunction with excisionase, integrase is also necessary for excision of the prophage from the host genome. |
| Rv0032 | *bioF2* | **1.023** | Possible 8-amino-7-oxononanoate synthase BioF2 (AONS) (8-amino-7-ketopelargonate synthase) (7-keto-8-amino-pelargonic acid synthetase) (7-KAP synthetase) (L-alanine--pimelyl CoA ligase) | int. metabolism & respiration | Could be involved in biotin biosynthesis (at the first step) [catalytic activity: 6-carboxyhexanoyl-CoA + L-alanine = 8-amino-7-oxononanoate + CoA + CO2]. |
| Rv0161 | *Rv0161* | **1.220** | Possible oxidoreductase | int. metabolism & respiration | Function unknown; probably involved in cellular metabolism. |
| Rv0327c | *cyp135A1* | **1.487** | Possible cytochrome P450 135A1 Cyp135A1 | int. metabolism & respiration | Cytochromes P450 are a group of heme-thiolate monooxygenases. They oxidize a variety of structurally unrelated compounds, including steroids, fatty acids, and |

| | | | | | |
|---|---|---|---|---|---|
| Rv0669c | *Rv0669c* | **1.062** | Possible hydrolase | int. metabolism & respiration | Function unknown; hydrolytic enzyme probably involved in cellular metabolism. |
| Rv1432 | *Rv1432* | **1.303** | Probable dehydrogenase | int. metabolism & respiration | Function unknown; probably involved in cellular metabolism |
| Rv1542c | *glbN* | **1.338** | Hemoglobin GlbN | int. metabolism & respiration | Oxygen transport |
| Rv2363 | *amiA2* | **1.286** | Probable amidase AmiA2 (aminohydrolase) | int. metabolism & respiration | Generates monocarboxylate from monocarboxylic acid amide [catalytic activity: a monocarboxylic acid amide + H(2)O = a monocarboxylate + NH(3)]. |
| Rv3109 | *moaA1* | **1.871** | Probable molybdenum cofactor biosynthesis protein A MoaA1 | int. metabolism & respiration | Involved in molybdenum cofactor biosynthesis; involved in the biosynthesis of molybdopterin precursor Z from guanosine. |
| Rv3379c | *dxs2* | **1.621** | Probable 1-deoxy-D-xylulose 5-phosphate synthase Dxs2 (1-deoxyxylulose-5-phosphate synthase) (DXP synthase) (DXPS) | int. metabolism & respiration | Catalyzes the acyloin condensation reaction between C atoms 2 and 3 of pyruvate and glyceraldehyde 3-phosphate to yield 1-deoxy-D-xylulose-5-phosphate (DXP). Possibly involved in deoxyxylulose-5-phosphate pathway (DXP) of isoprenoid biosynthesis (at the first step), and biosynthetic pathway to |
| Rv0672 | *fadE8* | **-1.049** | Probable acyl-CoA dehydrogenase FadE8 | lipid metabolism | Function unknown, but involved in lipid degradation. |
| Rv3820c | *papA2* | **1.209** | Possible conserved polyketide synthase associated protein PapA2 | lipid metabolism | Involved in sulfolipid-1 (SL-1) biosynthesis |
| Rv1430 | *PE16* | **1.098** | PE family protein PE16 | PE/PPE | Function unknown |
| Rv1917c | *PPE34* | **1.239** | PPE family protein PPE34 | PE/PPE | Function unknown |
| Rv1965 | *yrbE3B* | **1.383** | Conserved hypothetical integral membrane protein YrbE3B | virulence, detoxification, adaptation | Unknown |
| **W-7642** | | | | | |
| Rv0236c | *aftD* | **1.103** | Possible arabinofuranosyltransferase AftD | cell wall and cell processes | Involved in the biosynthesis of the mycobacterial cell wall |
| Rv0261c | *narK3* | **-1.551** | Probable integral membrane nitrite extrusion protein NarK3 (nitrite facilitator) | cell wall and cell processes | Involved in excretion of nitrite produced by the dissimilatory reduction of nitrate. Responsible for the translocation of the substrate across the membrane. |
| Rv0288 | *esxH or cfp7* | **1.049** | Low molecular weight protein antigen 7 EsxH (10 kDa antigen) (CFP-7) (protein TB10.4) | cell wall and cell processes | Function unknown. May be involved in virulence. |
| Rv0290 | *eccD3* | **1.115** | ESX conserved component EccD3. ESX-3 type VII secretion system protein. Probable transmembrane protein. | cell wall and cell processes | Unknown |
| Rv0522 | *gabP* | **-1.018** | Probable GABA permease GabP (4-amino butyrate transport carrier) (GAMA-aminobutyrate permease) | cell wall and cell processes | Involved in 4-aminobutyrate (GABA) degradation pathway. Transporter for GABA. Responsible for the translocation of the substrate across the membrane. |
| Rv0528 | *Rv0528* | **1.117** | Probable conserved transmembrane protein | cell wall and cell processes | Unknown |
| Rv0888 | *Rv0888* | **-1.439** | Probable exported protein | cell wall and cell processes | Unknown |
| Rv0935 | *pstC1* | **1.037** | Phosphate-transport integral membrane ABC transporter PstC1 | cell wall and cell processes | Involved in active transport of inorganic phosphate across the membrane (import); responsible for the translocation of the substrate across the membrane. This is one of the proteins required for binding-protein-mediated phosphate transport. |
| Rv1038c | *esxJ* | **-1.098** | ESAT-6 like protein EsxJ (ESAT-6 like protein 2) | cell wall and cell processes | Function unknown |
| Rv1198 | *esxL* | **1.100** | Putative ESAT-6 like protein EsxL (ESAT-6 like protein 4) | cell wall and cell processes | Unknown |
| Rv1226c | *Rv1226c* | **1.080** | Probable transmembrane protein | cell wall and cell processes | Unknown |

| | | | | | |
|---|---|---|---|---|---|
| Rv1227c | *Rv1227c* | **1.039** | Probable transmembrane protein | cell wall and cell processes | Unknown |
| Rv1342c | *Rv1342c* | **-1.075** | Conserved membrane protein | cell wall and cell processes | Unknown |
| Rv1362c | *Rv1362c* | **-1.042** | Possible membrane protein | cell wall and cell processes | Function unknown |
| Rv2094c | *tatA* | **-1.175** | Sec-independent protein translocase membrane-bound protein TatA | cell wall and cell processes | Involved in protein export: required for correct localization of precursor proteins bearing signal peptides with the twin arginine conserved motif S/T-R-R-X-F-L-K. This sec-independent pathway is termed tat for twin-arginine translocation system. This system mainly transports proteins [illegible] |
| Rv2157c | *murF* | **1.155** | Probable UDP-N-acetylmuramoylalanyl-D-glutamyl-2,6-diaminopimelate-D-alanyl-D-alanyl ligase MurF | cell wall and cell processes | Involved in cell wall formation; peptidoglycan biosynthesis. |
| Rv2158c | *murE* | **1.031** | Probable UDP-N-acetylmuramoylalanyl-D-glutamate-2,6-diaminopimelate ligase MurE | cell wall and cell processes | Involved in cell wall formation; peptidoglycan biosynthesis. |
| Rv2163c | *pbpB or ftsI* | **-1.056** | Probable penicillin-binding membrane protein PbpB | cell wall and cell processes | Involved in peptidoglycan biosynthesis |
| Rv2169c | *Rv2169c* | **-1.007** | Probable conserved transmembrane protein | cell wall and cell processes | Unknown |
| Rv2347c | *esxP* | **-1.144** | Putative ESAT-6 like protein EsxP (ESAT-6 like protein 7) | cell wall and cell processes | Function unknown |
| Rv3238c | *Rv3238c* | **-1.104** | Probable conserved integral membrane protein | cell wall and cell processes | Unknown |
| Rv3614c | *espD or snm10* | **-1.426** | ESX-1 secretion-associated protein EspD | cell wall and cell processes | Function unknown |
| Rv3615c | *espC or snm9* | **-1.369** | ESX-1 secretion-associated protein EspC | cell wall and cell processes | Function unknown |
| Rv3616c | *espA* | **-1.075** | ESX-1 secretion-associated protein A, EspA | cell wall and cell processes | Function unknown |
| Rv3807c | *Rv3807c* | **1.081** | Possible conserved transmembrane protein | cell wall and cell processes | Unknown |
| Rv3891c | *esxD* | **-1.256** | Possible ESAT-6 like protein EsxD | cell wall and cell processes | Function unknown |
| Rv0028 | *Rv0028* | **-1.535** | Conserved hypothetical protein | conserved hypotheticals | Unknown |
| Rv0460 | *Rv0460* | **-1.566** | Conserved hydrophobic protein | conserved hypotheticals | Function unknown |
| Rv0500A | *Rv0500A* | **-1.010** | Conserved protein | conserved hypotheticals | Function unknown |
| Rv0580c | *Rv0580c* | **-1.096** | Conserved protein | conserved hypotheticals | Function unknown |
| Rv0679c | *Rv0679c* | **-1.181** | Conserved threonine rich protein | conserved hypotheticals | Function unknown |
| Rv0739 | *Rv0739* | **-1.156** | Conserved hypothetical protein | conserved hypotheticals | Function unknown |
| Rv0856 | *Rv0856* | **-1.050** | Conserved hypothetical protein | conserved hypotheticals | Function unknown |
| Rv1190 | *Rv1190* | **-1.008** | Conserved hypothetical protein | conserved hypotheticals | Function unknown |
| Rv1268c | *Rv1268c* | **-1.286** | Hypothetical protein | conserved hypotheticals | Unknown |
| Rv1489A | *Rv1489A* | **-1.299** | Conserved protein | conserved hypotheticals | Function unknown |
| Rv1501 | *Rv1501* | **-1.319** | Conserved hypothetical protein | conserved hypotheticals | Function unknown |
| Rv1571 | *Rv1571* | **1.017** | Conserved protein | conserved hypotheticals | Function unknown |
| Rv1716 | *Rv1716* | **1.320** | Conserved hypothetical protein | conserved hypotheticals | Function unknown |
| Rv1718 | *Rv1718* | **1.076** | Conserved hypothetical protein | conserved hypotheticals | Function unknown |
| Rv1724c | *Rv1724c* | **-1.202** | Hypothetical protein | conserved hypotheticals | Unknown |
| Rv1754c | *Rv1754c* | **-1.022** | Conserved protein | conserved hypotheticals | Function unknown |
| Rv1810 | *Rv1810* | **-1.445** | Conserved protein | conserved hypotheticals | Function unknown |
| Rv1883c | *Rv1883c* | **-1.572** | Conserved hypothetical protein | conserved hypotheticals | Function unknown |
| Rv1904 | *Rv1904* | **-1.244** | Conserved hypothetical protein | conserved hypotheticals | Function unknown |

| | | | | | |
|---|---|---|---|---|---|
| Rv1995 | *Rv1995* | **-1.098** | Unknown protein | conserved hypotheticals | Unknown |
| Rv2023c | *Rv2023c* | **-1.006** | Hypothetical protein | conserved hypotheticals | Unknown |
| Rv2047c | *Rv2047c* | **1.051** | Conserved hypothetical protein | conserved hypotheticals | Unknown |
| Rv2143 | *Rv2143* | **-1.068** | Conserved hypothetical protein | conserved hypotheticals | Unknown |
| Rv2166c | *Rv2166c* | **-1.158** | Conserved protein | conserved hypotheticals | Unknown |
| Rv2247 | *accD6* | **1.012** | Conserved hypothetical protein | conserved hypotheticals | Unknown |
| Rv2293c | *Rv2293c* | **-1.038** | Conserved hypothetical protein | conserved hypotheticals | Unknown |
| Rv2522c | *Rv2522c* | **1.037** | Conserved hypothetical protein | conserved hypotheticals | Function unknown |
| Rv2664 | *Rv2664* | **-1.166** | Hypothetical protein | conserved hypotheticals | Unknown |
| Rv2670c | *Rv2670c* | **1.023** | Conserved hypothetical protein | conserved hypotheticals | Function unknown |
| Rv2897c | *Rv2897c* | **1.217** | Conserved hypothetical protein | conserved hypotheticals | Function unknown |
| Rv2974c | *Rv2974c* | **1.092** | Conserved hypothetical alanine rich protein | conserved hypotheticals | Function unknown |
| Rv3005c | *Rv3005c* | **-1.008** | Conserved hypothetical protein | conserved hypotheticals | Function unknown |
| Rv3031 | *Rv3031* | **1.015** | Conserved protein | conserved hypotheticals | Function unknown |
| Rv3046c | *Rv3046c* | **-1.259** | Conserved protein | conserved hypotheticals | Function unknown |
| Rv3073c | *Rv3073c* | **-1.042** | Conserved hypothetical protein | conserved hypotheticals | Function unknown |
| Rv3237c | *Rv3237c* | **-1.089** | Conserved protein | conserved hypotheticals | Function unknown |
| Rv3292 | *Rv3292* | **-1.039** | Conserved hypothetical protein | conserved hypotheticals | Function unknown |
| Rv3412 | *Rv3412* | **-1.034** | Conserved hypothetical protein | conserved hypotheticals | Function unknown |
| Rv3421c | *Rv3421c* | **1.067** | Conserved hypothetical protein | conserved hypotheticals | Function unknown |
| Rv3422c | *Rv3422c* | **1.127** | Conserved hypothetical protein | conserved hypotheticals | Function unknown |
| Rv3433c | *Rv3433c* | **1.266** | Conserved protein | conserved hypotheticals | Function unknown |
| Rv3555c | *Rv3555c* | **1.146** | Conserved protein | conserved hypotheticals | Function unknown |
| Rv3633 | *Rv3633* | **-1.422** | Conserved protein | conserved hypotheticals | Function unknown |
| Rv3717 | *Rv3717* | **-1.084** | Conserved hypothetical protein | conserved hypotheticals | Function unknown |
| Rv3733c | *Rv3733c* | **-1.387** | Conserved hypothetical protein | conserved hypotheticals | Function unknown |
| Rv0001 | *dnaA* | **-1.118** | Chromosomal replication initiator protein DnaA | information pathways | Plays an important role in the initiation and regulation of chromosomal replication. Binds to the origin of replication; it binds specifically double-stranded DNA at a 9 bp consensus (DNAA box): 5'-TTATC(C/A)A(C/A)A-3'. DNAA binds to ATP and to acidic phospholipids. DNAA protein binds the origin of replication (oriC), ATP and ADP, and exhibited weak ATPase [illegible] |
| Rv1189 | *sigI* | **-1.550** | Possible alternative RNA polymerase sigma factor SigI | information pathways | The sigma factor is an initiation factor that promotes attachment of the RNA polymerase to specific initiation sites |
| Rv2191 | *Rv2191* | **1.242** | Conserved hypothetical protein | information pathways | Unknown |
| Rv2592c | *ruvB* | **1.103** | Probable holliday junction DNA helicase RuvB | information pathways | forms a complex with RUVA. RUVB could possess weak ATPase activity, which will be stimulated by the RUVA protein in the presence of DNA. The RUVA-RUVB complex in the presence of ATP renatures cruciform structure in supercoiled DNA with palindromic sequence, indicating that it may promote strand exchange reactions in homologous recombination. RUVAB is an helicase that mediates the [illegible] |
| Rv2985 | *mutT1* | **-1.007** | Possible hydrolase MutT1 | information pathways | Function unknown; hydrolytic enzyme. Possibly involved in removal of damaged nucleotide. |

| | | | | | |
|---|---|---|---|---|---|
| Rv3201c | *Rv3201c* | **1.146** | Probable ATP-dependent DNA helicase | information pathways | Has both ATPase and helicase activities |
| Rv3241c | *Rv3241c* | **-1.018** | Conserved protein | information pathways | Function unknown, but may be involved in transduction |
| Rv3286c | *sigF* | **-1.309** | Alternative RNA polymerase sigma factor SigF | information pathways | The sigma factor is an initiation factor that promotes attachment of the RNA polymerase to specific initiation sites and then is released. Thought to be involved in survival and proliferation in lung granulomas during infection. Thought to be involved in virulence and persistence processes. Modulates expression of the 16 KDa alpha-crystallin homologue/Rv2031c. [illegible] |
| Rv3287c | *rsbW or usfX* | **-1.243** | Anti-sigma factor RsbW (sigma negative effector) | information pathways | Binds to sigma and blocks its ability to form an RNA polymerase holoenzyme. Regulates negatively SIGF\|Rv3286c, and negatively regulated by Rv1365c\|RSFA and |
| Rv3585 | *radA* | **1.147** | DNA repair protein RadA (DNA repair protein SMS) | information pathways | Involved in genetic recombination. May play a role in the repair of endogenous alkylation damage. |
| Rv0148 | *Rv0148* | **-1.005** | Probable short-chain type dehydrogenase/reductase | int. metabolism & respiration | Function unknown; possibly involved in cellular metabolism. |
| Rv0149 | *Rv0149* | **-1.150** | Possible quinone oxidoreductase (NADPH:quinone oxidoreductase) (zeta-crystallin) | int. metabolism & respiration | Possibly binds NADP and acts through a one-electron transfer process. Quinones are supposed to be the best substrates. May act in the detoxification of xenobiotics [catalytic activity: NADPH + quinone = NADP+ + semiquinone] |
| Rv0156 | *pntAb* | **1.238** | Probable NAD(P) transhydrogenase (subunit alpha) PntAb [second part; integral membrane protein] (pyridine nucleotide transhydrogenase subunit alpha) (nicotinamide nucleotide transhydrogenase subunit alpha) | int. metabolism & respiration | The transhydrogenation between NADH and NADP is coupled to respiration and ATP hydrolysis and functions as a proton pump across the membrane [catalytic activity: NADPH + NAD+ = NADP+ + NADH]. |
| Rv0157 | *pntB* | **1.191** | Probable NAD(P) transhydrogenase (subunit beta) PntB [integral membrane protein] (pyridine nucleotide transhydrogenase subunit beta) (nicotinamide nucleotide transhydrogenase subunit beta) | int. metabolism & respiration | The transhydrogenation between NADH and NADP is coupled to respiration and ATP hydrolysis and functions as a proton pump across the membrane [catalytic activity: NADPH + NAD+ = NADP+ + NADH]. |
| Rv0186 | *bglS* | **1.015** | Probable beta-glucosidase BglS (gentiobiase) (cellobiase) (beta-D-glucoside glucohydrolase) | int. metabolism & respiration | Possibly involved in degradation [catalytic activity: hydrolysis of terminal, non-reducing beta-D-glucose residues with release |
| Rv0211 | *pckA or pck1 or p* | **-1.157** | Probable iron-regulated phosphoenolpyruvate carboxykinase [GTP] PckA (phosphoenolpyruvate carboxylase) (PEPCK)(pep carboxykinase) | int. metabolism & respiration | Rate-limiting gluconeogenic enzyme [catalytic activity: GTP + oxaloacetate = GDP + phosphoenolpyruvate + CO2]. |
| Rv0291 | *mycP3* | **1.100** | Probable membrane-anchored mycosin MycP3 (serine protease) (subtilisin-like protease) (subtilase-like) (mycosin-3) | int. metabolism & respiration | Thought to have proteolytic activity. |
| Rv0524 | *hemL* | **1.036** | Probable glutamate-1-semialdehyde 2,1-aminomutase HemL (GSA) (glutamate-1-semialdehyde aminotransferase) (GSA-at) | int. metabolism & respiration | Involved in porphyrin biosynthesis by the C5 pathway (at the second step) [catalytic activity: (S)-4-amino-5-oxopentanoate = 5-aminolevulinate]. |
| Rv0526 | *Rv0526* | **1.035** | Possible thioredoxin protein (thiol-disulfide interchange protein) | int. metabolism & respiration | Possibly acts on thioredoxin |

| | | | | | |
|---|---|---|---|---|---|
| Rv0527 | *ccdA* | **1.203** | Possible cytochrome C-type biogenesis protein CcdA | int. metabolism & respiration | Possibly involved in cytochrome C synthesis. Might transfer reducing equivalents across the cytoplasmic membrane, promoting efficient disulfide bond isomerization of proteins localized on the outer surface of the membrane. |
| Rv0886 | *fprB* | **-1.128** | Probable NADPH:adrenodoxin oxidoreductase FprB (adrenodoxin reductase) (AR) (ferredoxin-NADP(+) reductase) | int. metabolism & respiration | Serves as the first electron transfer protein in all the P450 systems [catalytic activity: reduced adrenodoxin + NADP+ = oxidized adrenodoxin + NADPH]. |
| Rv0889c | *citA* | **1.000** | Probable citrate synthase II CitA | int. metabolism & respiration | Involved in tricarboxylic acid cycle (KREBS cycle) [catalytic activity: citrate + CoA = acetyl-CoA + H2O + oxaloacetate]. |
| Rv1076 | *lipU* | **-1.153** | Possible lipase LipU | int. metabolism & respiration | Hydrolyses lipids |
| Rv1188 | *Rv1188* | **-1.684** | Probable proline dehydrogenase | int. metabolism & respiration | Oxidizes proline to glutamate for use as a carbon and nitrogen source [catalytic activity: L-proline + acceptor + H2O = (S)-1-pyrroline-5-carboxylate + reduced acceptor] |
| Rv1569 | *bioF1* | **1.081** | Probable 8-amino-7-oxononanoate synthase BioF1 (AONS) (8-amino-7-ketopelargonate synthase) (7-keto-8-amino-pelargonic acid synthetase) (7-KAP synthetase) (L-alanine--pimelyl CoA ligase) | int. metabolism & respiration | Involved in biotin biosynthesis (at the first step) [catalytic activity: 6-carboxyhexanoyl-CoA + L-alanine = 8-amino-7-oxononanoate + CoA + CO2]. |
| Rv1613 | *trpA* | **1.063** | Probable tryptophan synthase, alpha subunit TrpA | int. metabolism & respiration | Tryptophan biosynthesis pathway (fifth - last step). The alpha subunit is responsible for the ALDOL cleavage of indoleglycerol phosphate to indole and glyceraldehyde 3- phosphate. [catalytic activity: L-serine + 1-(indol-3-YL)glycerol 3-phosphate = L-tryptophan + glyceraldehyde 3-phosphate + H(2)O.] |
| Rv1655 | *argD* | **1.064** | Probable acetylornithine aminotransferase ArgD | int. metabolism & respiration | Arginine biosynthesis (fourth step) [catalytic activity: N2-acetyl-L-ornithine + 2-oxoglutarate = N-acetyl-L-glutamate 5- |
| Rv1714 | *Rv1714* | **1.154** | Probable oxidoreductase | int. metabolism & respiration | Function unknown; probably involved in cellular metabolism |
| Rv1882c | *Rv1882c* | **-1.371** | Probable short-chain type dehydrogenase/reductase | int. metabolism & respiration | Function unknown; probably involved in cellular metabolism |
| Rv2457c | *clpX* | **-1.007** | Probable ATP-dependent CLP protease ATP-binding subunit ClpX | int. metabolism & respiration | ATP-dependent specificity component of the CLP protease. It directs the protease to specific substrates. Can perform chaperone functions in the absence of CLPP). |
| Rv2678c | *hemE* | **1.105** | Probable uroporphyrinogen decarboxylase HemE (uroporphyrinogen III decarboxylase) (URO-D) (UPD) | int. metabolism & respiration | Involved in porphyrin biosynthesis [catalytic activity: uroporphyrinogen III = coproporphyrinogen + 4 CO(2)]. |
| Rv2739c | *Rv2739c* | **1.045** | Possible alanine rich transferase | int. metabolism & respiration | Function unknown; probably involved in cellular metabolism. |
| Rv2987c | *leuD* | **1.141** | Probable 3-isopropylmalate dehydratase (small subunit) LeuD (isopropylmalate isomerase) (alpha-IPM isomerase) (IPMI) | int. metabolism & respiration | Involved in leucine biosynthesis (at the second step) [catalytic activity: 3-isopropylmalate = 2-isopropylmaleate + H(2)O (also catalyses 2-isopropylmaleate + H(2)O = 3-hydroxy-4-methyl-3- |
| Rv2988c | *leuC* | **1.091** | Probable 3-isopropylmalate dehydratase (large subunit) LeuC (isopropylmalate isomerase) (alpha-IPM isomerase) (IPMI) | int. metabolism & respiration | Involved in leucine biosynthesis (at the second step) [catalytic activity: 3-isopropylmalate = 2-isopropylmaleate + H(2)O (also catalyses 2-isopropylmaleate + H(2)O = 3-hydroxy-4-methyl-3- |
| Rv3151 | *nuoG* | **1.126** | Probable NADH dehydrogenase I (chain G) NuoG (NADH-ubiquinone oxidoreductase chain G) | int. metabolism & respiration | Involved in aerobic\|anaerobic respiration [catalytic activity: NADH + ubiquinone = NAD(+) + ubiquinol]. |

| | | | | | |
|---|---|---|---|---|---|
| Rv3322c | *Rv3322c* | **-1.059** | Possible methyltransferase | int. metabolism & respiration | Could cause methylation. |
| Rv3323c | *moaX* | **-1.145** | Probable MoaD-MoaE fusion protein MoaX | int. metabolism & respiration | Thought to be involved in molybdenum cofactor biosynthesis. |
| Rv3324c | *moaC3* | **-1.211** | Probable molybdenum cofactor biosynthesis protein C 3 MoaC3 | int. metabolism & respiration | Thought to be involved in the biosynthesis of molybdopterin. |
| Rv3419c | *gcp* | **1.024** | Probable O-sialoglycoprotein endopeptidase Gcp (glycoprotease) | int. metabolism & respiration | Hydrolysis of O-sialoglycoproteins; cleaves 31-ARG-\|-asp-32 bond in glycophorin A. Does not cleave unglycosylated proteins, desialylated glycoproteins or glycoproteins that are only N-glycosylated. Could be a metalloprotease. |
| Rv3742c | *Rv3742c* | **-1.428** | Possible oxidoreductase | int. metabolism & respiration | Function unknown; probably involved in cellular metabolism. |
| Rv3854c | *ethA* | **-1.191** | Monooxygenase EthA | int. metabolism & respiration | Activates the pro-drug ethionamide (ETH); induced ETH sensitivity when overexpressed in Mycobacterium tuberculosis. |
| Rv0129c | *fbpC or mpt45* | **-1.043** | Secreted antigen 85-C FbpC (85C) (antigen 85 complex C) (AG58C) (mycolyl transferase 85C) (fibronectin-binding protein C) | lipid metabolism | Proteins of the antigen 85 complex are responsible for the high affinity of mycobacteria to fibronectin. Possesses a mycolyltransferase activity required for the biogenesis of trehalose dimycolate (cord factor), a dominant structure |
| Rv2246 | *kasB* | **1.030** | Acetyl/propionyl-CoA carboxylase (beta subunit) AccD6 | lipid metabolism | Involved in fatty acid biosynthesis (mycolic acids synthesis) [catalytic activity: ATP + propionyl-CoA + CO(2) + H(2)O = ADP + orthophosphate + methylmalonyl-CoA]. |
| Rv2482c | *plsB2* | **-1.076** | Probable glycerol-3-phosphate acyltransferase PlsB2 (GPAT) | lipid metabolism | Involved in phospholipid biosynthesis (at the first step). May also function in the regulation of membrane biogenesis [catalytic activity: acyl-CoA + SN-glycerol 3-phosphate = CoA + |
| Rv2523c | *acpS* | **1.238** | holo-[acyl-carrier protein] synthase AcpS (holo-ACP synthase) (CoA:APO-[ACP]pantetheinephosphotransferase) (CoA:APO-[acyl-carrier protein]pantetheinephosphotransferase) | lipid metabolism | Biosynthesis of fatty acids and lipids. Transfers the 4'-phosphopantetheine moiety from coenzyme A to a SER of acyl-carrier protein. Catalyzes the formation of holo-ACP, which mediates the transfer of acyl fatty-acid intermediates during the biosynthesis of fatty acids and lipids [catalytic activity: CoA + APO-[acyl-carrier protein] = adenosine 3',5'-bisphosphate + [illegible] |
| Rv2524c | *fas* | **1.310** | Probable fatty acid synthase Fas (fatty acid synthetase) | lipid metabolism | Involved in lipid metabolism. Fatty acid synthetase catalyzes the formation of long-chain fatty acids from acetyl-CoA, |
| Rv2933 | *ppsC* | **1.210** | Phenolpthiocerol synthesis type-I polyketide synthase PpsC | lipid metabolism | Involved in phenolpthiocerol and phthiocerol dimycocerosate (dim) biosynthesis: extension with malony CoA (complete |
| Rv2934 | *ppsD* | **1.107** | Phenolpthiocerol synthesis type-I polyketide synthase PpsD | lipid metabolism | Involved in phenolpthiocerol and phthiocerol dimycocerosate (dim) biosynthesis: extension with methylmalony CoA (partial |
| Rv2947c | *pks15* | **1.035** | Probable polyketide synthase Pks15 | lipid metabolism | Polyketide synthase possibly involved in lipid synthesis |
| Rv3556c | *fadA6* | **1.228** | Probable acetyl-CoA acetyltransferase FadA6 (acetoacetyl-CoA thiolase) | lipid metabolism | Function unknown, but involved in lipid degradation [catalytic activity: 2 acetyl-CoA = CoA + acetoacetyl-CoA]. |
| Rv3563 | *fadE32* | **1.206** | Probable acyl-CoA dehydrogenase FadE32 | lipid metabolism | Function unknown, but involved in lipid degradation. |
| Rv3734c | *tgs2* | **-1.447** | Putative triacylglycerol synthase (diacylglycerol acyltransferase) Tgs2 | lipid metabolism | May be involved in synthesis of triacylglycerol |
| Rv0151c | *PE1* | **-1.040** | PE family protein PE1 | PE/PPE | Function unknown |
| Rv0160c | *PE4* | **-1.291** | PE family protein PE4 | PE/PPE | Function unknown |
| Rv0285 | *PE5* | **1.006** | PE family protein PE5 | PE/PPE | Function unknown |
| Rv0304c | *PPE5* | **-1.078** | PPE family protein PPE5 | PE/PPE | Function unknown |
| Rv1067c | *PE_PGRS19* | **1.048** | PE-PGRS family protein PE_PGRS19 | PE/PPE | Function unknown |

| | | | | | |
|---|---|---|---|---|---|
| Rv1068c | *PE_PGRS20* | **1.021** | PE-PGRS family protein PE_PGRS20 | PE/PPE | Function unknown |
| Rv1325c | *PE_PGRS24* | **1.091** | PE-PGRS family protein PE_PGRS24 | PE/PPE | Function unknown |
| Rv1646 | *PE17* | **-1.377** | PE family protein PE17 | PE/PPE | Function unknown |
| Rv1808 | *PPE32* | **-1.010** | PPE family protein PPE32 | PE/PPE | Function unknown |
| Rv1809 | *PPE33* | **-1.634** | PPE family protein PPE33 | PE/PPE | Function unknown |
| Rv2353c | *PPE39* | **-1.093** | PPE family protein PPE39 | PE/PPE | Function unknown |
| Rv3135 | *PPE50* | **1.314** | PPE family protein PPE50 | PE/PPE | Function unknown |
| Rv0165c | *mce1R* | **-1.068** | Probable transcriptional regulatory protein Mce1R (probably GntR-family) | regulatory proteins | Involved in transcriptional mechanism |
| Rv1556 | *Rv1556* | **-1.157** | Possible regulatory protein | regulatory proteins | Possibly involved in a transcriptional mechanism |
| Rv2231c | *cobC* | **1.107** | Protein tyrosine kinase transcriptional regulatory protein PtkA | regulatory proteins | Involved in signal transduction (via phosphorylation). Can phosphorylate PTPA\|Rv2234 and the peptide substrate myelin basic protein (MBP) [catalytic activity: ATP + a protein = ADP + |
| Rv2989 | *Rv2989* | **1.019** | Probable transcriptional regulatory protein | regulatory proteins | Involved in transcriptional mechanism. |
| Rv3219 | *whiB1* | **-1.192** | Transcriptional regulatory protein WhiB-like WhiB1. Contains [4FE-4S]2+ cluster. | regulatory proteins | Involved in transcriptional mechanism. |
| Rv3855 | *ethR* | **-1.055** | Transcriptional regulatory repressor protein (TetR-family) EthR | regulatory proteins | Regulates negatively the production of ETHA. Induced ETH resistance when overexpressed in Mycobacterium tuberculosis. |
| RVnc0013 | *mcr11* | **1.206** | | Stable rnas | |
| RVnc0015 | *mcr15* | **-1.085** | | Stable rnas | |
| Rvnr01 | *rrs* | **1.203** | | Stable rnas | |
| Rvnr02 | *rrl* | **1.152** | | Stable rnas | |
| Rv0064A | *vapB1* | **-1.049** | Possible antitoxin VapB1 | virulence, detoxification, adaptation | Unknown |
| Rv1636 | *TB15.3* | **-1.114** | Iron-regulated universal stress protein family protein TB15.3 | virulence, detoxification, adaptation | Function unknown |
| Rv1740 | *vapB34* | **-1.143** | Possible antitoxin VapB34 | virulence, detoxification, adaptation | Unknown |
| Rv1839c | *vapB13* | **-1.028** | Possible antitoxin VapB13 | virulence, detoxification, adaptation | Unknown |
| Rv2230c | *Rv2230c* | **1.031** | Possible toxin VapC16 | virulence, detoxification, adaptation | Unknown |
| Rv2429 | *ahpD* | **-1.128** | Alkyl hydroperoxide reductase D protein AhpD (alkyl hydroperoxidase D) | virulence, detoxification, adaptation | Involved in oxidative stress response. LPDC\|Rv0462, DLAT\|Rv2215, AHPD\|Rv2429, and AHPC\|Rv2428 constitute an NADH-dependent peroxidase and peroxynitrite reductase that provides protection against oxidative stress. |
| Rv2527 | *vapC17* | **-1.213** | Possible toxin VapC17 | virulence, detoxification, adaptation | Unknown |
| Rv3320c | *vapC44* | **-1.188** | Possible toxin VapC44. Contains PIN domain. | virulence, detoxification, adaptation | Unknown |
| Rv3321c | *vapB44* | **-1.362** | Possible antitoxin VapB44 | virulence, detoxification, adaptation | Unknown |
| Rv2386a | *Rv2386a* | **-1.071** | | | |
