## Supplemental Table S5 for "Exposure of *Mycobacterium tuberculosis* to human alveolar lining fluid shows temporal and strain-specific adaptation to the lung environment"

**Supplemental Table S5**. RT-qPCR primers for selected genes used for RNA-seq validation.

| Gene | Locus tag | Operon/function | Primers (sequence 5'-3') | Amplicon (bp) | Reference |
|---|---|---|---|---|---|
| ***echA20*** | *Rv3550* | KstR2 operon | echA20-F: ATGCGACGGCTGTTCTTTAC<br>echA20-R: AAACCTTGCTCCATCCGGTA | 224 | This study |
| ***Rv3557c*** | *Rv3557c* | KstR2 operon | KstR2-F: TCTACCAGGATGAAGCGCAA<br>KstR2-R: AGGTGGTGTCACGGATGAAT | 177 | This study |
| ***mymA*** | *Rv3083* | Maintaining mycolic acid composition | mymA-F: ATCGCTGGGTCTACGAGTTT<br>mymA-R: CAGGTAGTTCGGGGTGAAGT | 167 | This study |
| ***papA1*** | *Rv3824c* | DAT/PAT and SL-1 | papA1-F: GAGCTTCGAGATACCGACCA<br>papA1-R: GCTGGCATAGAACGTGAAGG | 198 | This study |
| ***papA3*** | *Rv1182* | DAT/PAT and SL-1 | papA3-F: GGGTTGTTTTCGGTTTGGGA<br>papA3-R: TGCGTACATCAGGTGGAACT | 127 | This study |
| ***mmpl10*** | *Rv1183* | DAT/PAT and SL-1 | mmpl10-F: CCGTGCGGTACTTCATTCAG<br>mmpl10-R: TCAGCCAGGGAGGTATTTGG | 115 | This study |
| ***mmpl8*** | *Rv3823c* | DAT/PAT and SL-1 | mmpl8-F: GGGCTGAATGTAGACCAAC<br>mmpl8-R: ATCCGACGACAGACACCTTG | 199 | Garima et al. 2015 |
| ***mprA*** | *Rv0981* | 2 component system, stress resistance, establishment and maintenance of persistent infection, associated with hypoxia, starvation and iron metabolism | mprA-F: GTCGCTTTCCTTCAATGGCT<br>mprA-R: CATGACATCCAGGACCAACG | 106 | This study |
| ***sigF*** | *Rv3286c* | Sigma factor, associated with virulence and persistence | sigF-F: CTGCATCTGCGGCTAGGTA<br>sigF-R: TCGATGGACAAGGTGTGGTA | 155 | Williams et al. 2007 |
| ***espC*** | *Rv3615c* | Required for ESX-1 function. Required for either stability or expression of EspA. Part of the Rv3612c-Rv3616c operon regulated by espR | espC-F: TGTGTACTTGACTGCCCACA<br>espC-R: AACAACCCGTCGATAGCCTT | 138 | This study |
| ***esxH*** | *Rv0288* | Part of the ESX-3. With EsxG, impairs host phagosome maturation, promoting intracellular bacterial growth | esxH-F: GTCACGCCGGGGATATGG<br>esxH-R: GCTGGACATCGCATGATAGG | 185 | This study |
